# Overcoming EGFR resistance by monovalent and bident inhibitors targeting Cys775

**DOI:** 10.64898/2025.11.28.691243

**Authors:** Zhengnian Li, Jie Jiang, Yaning Wang, Scott B. Ficarro, Stephen J. Collins, Tyler S. Beyett, Ilse K. Schaeffner, Isidoro Tavares, Felix H. Gottlieb, Dhiraj Suda, Prafulla C. Gokhale, Leah M. Black-Holmes, Dimitris Gazgalis, Michael J. Eck, Pasi A. Jänne, Jarrod A. Marto, Jianwei Che, Nathanael S. Gray, Tinghu Zhang

**Author notes:** Authors contributed equally to this work (Zhengnian Li, Jie Jiang, Yaning Wang). Corresponding authors: Tinghu Zhang Stanford School of Medicine 290 Jane Stanford Way Stanford, CA 94305 Nathanael S. Gray Stanford School of Medicine 290 Jane Stanford Way Stanford, CA 94305 Jianwei Che Dana-Farber Cancer Institute 450 Brookline Ave, Boston, MA 02215.

## Abstract

Covalent targeting of EGFR cysteine 797 by osimertinib is one of the most successful breakthroughs in targeted therapy, fundamentally transforming the treatment landscape for non-small cell lung cancer (NSCLC) patients. However, resistance driven by mutation of C797 remains a major clinical challenge. Developing novel covalent strategies beyond C797 targeting presents a compelling opportunity for next-generation EGFR inhibitors. We first demonstrated that cysteine 775, located deep within the ATP-binding pocket, is accessible by a rationally designed covalent molecule ZNL-3, which as the first-in-class covalent cysteine 775 inhibitor exhibited strong efficacy in osimertinib-resistant mouse models. To further enhance resilience to resistance-causing mutations, we developed a dual-warhead, bident compound—YNW-1—which covalently targets both cysteine 775 and 797 simultaneously. YNW-1 is the first intramolecular lock to exhibit balanced reactive efficiency on both cysteines, rendering single-site mutations ineffective to confer resistance. The discovery of ZNL-3 and YNW-1 represents significant advancements in EGFR-targeted drug development, and further optimization toward clinical translation is a worthwhile strategy.

SIGNIFICANCE: This study establishes the therapeutic potential of an EGFR covalent inhibitor through unprecedented targeting of cysteine 775 and provides the first demonstration that dual cysteine engagement offers superior efficacy over conventional covalent inhibitors by delaying resistance.

## Introduction

Mutations in the epidermal growth factor receptor (EGFR) gene are major drivers in non-small cell lung cancer (NSCLC), which accounts for over 80% of all lung cancer cases^1,2^. Targeting EGFR with covalent small-molecule inhibitors has been one of the most successful strategies in precision oncology. Osimertinib, a third-generation covalent EGFR inhibitor, selectively modifies cysteine 797 (C797) and has demonstrated superior potency and efficacy compared to first-generation reversible inhibitors such as gefitinib and erlotinib, which target EGFR mutations like L858R^3^. Its broad activity across a range of EGFR mutations is largely attributed to its covalent binding mechanism, which confers prolonged target engagement and enhanced pharmacodynamic effects^4–6^. However, resistance to osimertinib inevitably emerges. Among EGFR-dependent mechanisms, the C797S mutation, which abolishes the covalent binding site, is the most prevalent and represents a major unmet need^7–9^. Although several fourth-generation strategies have been pursued, including allosteric inhibitors, PROTAC degraders, and reversible C797S-active inhibitors^10–14^, each approach faces limitations in mutation coverage, drug-like properties, or kinase selectivity, and convincing clinical progress remains limited.

To address resistance driven by loss of C797 engagement, we explored alternative nucleophilic residue within the EGFR protein that could serve as an additional covalent anchoring point. This led to the discovery of cysteine 775 (C775), located in the back pocket of the active site^15^. Compared to C797, C775 is more deeply buried and spatially constrained, less nucleophilic due to the lack of a polarizing environment that make it challenging to target but also offer the potential for highly selective covalent inhibition as it is not common across the kinome^15^. Our prior study demonstrated that C775 is targetable through a bidentate design (ZNL-0056) linking C797 and C775, offering initial proof-of-concept, although the resulting inhibitor remained largely dependent on bonding with C797^15^.

In this study, we pursued a reverse design strategy, beginning with optimized of C775. This approach led to the identification of ZNL-3, a potent mutant EGFR inhibitor primarily driven by covalency with C775. Building upon this scaffold, we further developed a bidentate compound YNW-1, which is mechanistically distinct from ZNL-0056 and exhibits equal efficiency on both C797 and C775, offering enhanced resilience to mutation-driven resistance. Furthermore, we demonstrated that the single-warhead compound ZNL-3 exhibits high oral dosing efficacy in a xenograft mouse model harboring the L858R/C797S EGFR mutation, highlighting its promising therapeutic potential for patients who progress on Osimertinib treatment. For the bidentate compound YNW-1, in vitro resistance studies showed a lower mutation frequency at either cysteine compared to Osimertinib under the same conditions. This suggests that YNW-1 may offer enhanced durability against EGFR mutation–induced resistance in clinical practice.

## Results

### Evolution of Designs on Cysteine775-Targeting Covalent Inhibitor

The wealth of publicly accessible co-crystal structures of EGFR provides the opportunity for structure-guided design of many potential C775 targeted covalent inhibitors. We previously reported a two-step approach utilizing a benzimidazole scaffold in which initial reaction with C797 would tether the compound to the ATP-binding site thereby achieving sufficient induced proximity for a slower covalent bond formation to the less reactive C775 residue. While this strategy successfully delivered the first dual covalent C775/C797 inhibitor, ZNL-0056, it renders the compound highly susceptible to resistance through mutation of C797. The question of whether targeting C775 itself is a viable strategy for achieving a desired pharmacology remains to be addressed. We suspected that poor accessibility and lower reactivity of C775 relative to C797 might contribute to the challenge and indeed C775 has not scored as a ligandable cysteine in any reported covalent fragment screening chemoproteomics studies^16^. We therefore decided to explore the reverse approach of designing compounds that would target C775 preferentially over C797. Using available liganded EGFR co-structures we designed and synthesized a variety of molecules bearing electrophiles that were predicted to project towards C775. We developed a mass-spectrometry based protein labeling assay with Cys797Ser mutant EGFR to enable detection of even weak covalent binders. Although many designs were unsuccessful, a focused medicinal chemistry campaign based on a dihydropyrimido[4,5-d]pyrimidin-4-one scaffold led to the identification of a promising hit compound, ZNL-1(Figure 1A). Among over 50 designed analogs, ZNL-1 demonstrated robust target engagement, binding over 90% of the EGFR^L858R^ mutant in mass-spectrometry analysis (Figure 1B). ZNL-1 inhibited EGFR kinase activity in a homogeneous time resolved fluorescence (HTRF) kinase assay with an apparent IC_50_ of 17.6 nM, representing a 16-fold less potent than Osimertinib (Figure 1C). To enable structure-based optimization, we solved the co-crystal structure of ZNL-1 bond to EGFR. As shown in Figure 1D, ZNL-1 forms the expected two hydrogen bonds with methionine 793 in the hinge through 2-aminopyrimide moiety and the methylated piperazine was exposed to the solvent region. The benzyl group that possesses an acrylamide warhead adapted a vertical pose to the urea ring and was well accommodated into the hydrophobic back pocket, with the acrylamide warhead forming a covalent bond to C775. The amide group adopts a non-canonical cis configuration to form two hydrogen bonds with aspartic acid 855 and the methyl group on second N of urea reaches to the sugar pocket (Figure S1 and Table S1). Given that covalent bond formation is a typically slow step and requires proper geometry to form transition state could improve the kinetics of nucleophilic reaction with thiol group of cysteine. To test this hypothesis, we have systematically modified ZNL-1 to improve reversible interactions and the warhead orientation toward C775, leading to more than 100 designed analogs. Among this series, ZNL-2 — featuring a cyclized ring bridging the benzyl group and the acrylamide NH — proved to be the most effective in enhancing potency, exhibiting an apparent biochemical IC_50_ of 0.49 nM, while maintaining high labeling efficiency (67%) on L858R EGFR (Figure 1A and 1B). Next, a stereo selective synthetic route was developed to enable the synthesis of the enantiomeric pure compounds, ZNL-2S and ZNL-2R. The S enantiomer exhibited approximately 40-fold lower apparent IC_50_ in the HTRF kinase assay as compared to the R enantiomer (Figure 1C). Further optimization on the core heterocycle resulted in the discovery of ZNL-3 which is with pyrido[2,3-*d*] pyrimidin-7(8*H*)-one scaffold and equipped with methyl on phenyl group which was expected improve compound stability and to modulate the warhead reactivity (Figure 1A). Labeling by intact MS was >90% with single digit nanomolar apparent IC_50_ in HTRF assay respectively (Figure 1B and 1C), while the reversible counterpart ZNL-3r lost significant potency (∼79-fold). More importantly, ZNL-3 demonstrated markedly improved metabolic stability in both hepatocyte and whole blood assay. In the mouse whole blood, the half-life (T_1/2_) of ZNL-3 increased to 205 min, a more than 16-fold improvement over ZNL-2, which displayed a T_1/2_ of only 12.4 min (Table S2).

**Figure 1.**
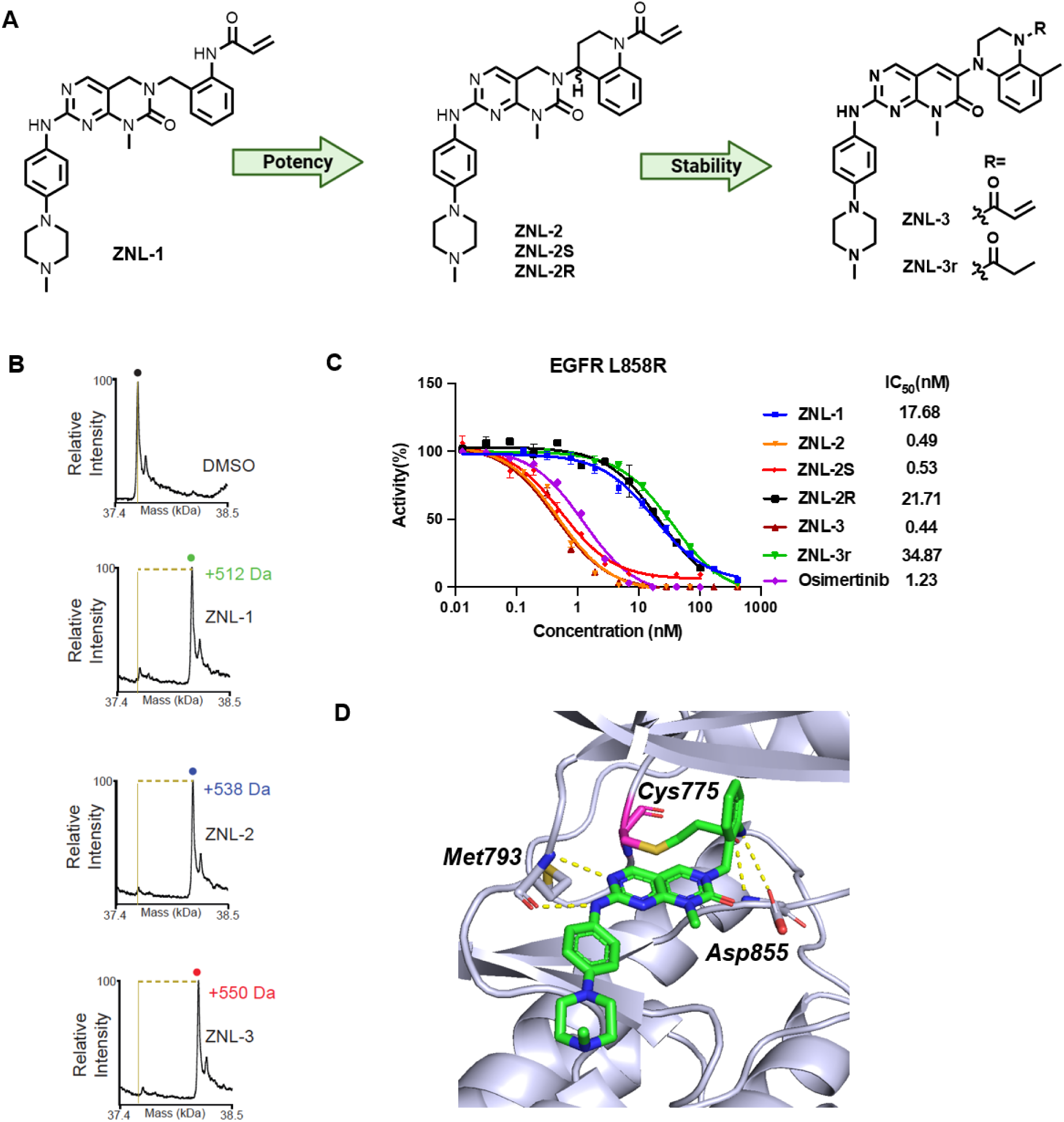
Evolution of designs on cysteine775-targeting covalent inhibitor **A.** Compound structures for EGFR cysteine-targeting molecules. **B.** Intact spectrometry analysis of EGFR L858R treated with indicated compound or DMSO. **C.** Biochemical IC_50_s for the indicated compounds against EGFR L858R. Data are presented as the mean±SEM. Representative data from three independent experiments are shown **D.** Cocrystal structure of ZNL-1 with WT EGFR

### ZNL-3 Covalently Inhibits EGFR Cys797-Ser Mutation and is Efficacious in L858R/C797S Xenograft Mouse Model

EGFR-dependent Ba/F3 cells are a well-established and widely used platform in the EGFR-targeted drug discovery. Accordingly, we employed three engineered Ba/F3 cells carrying single or double EGFR mutations - L858R, L858R/C797S and L858R/C775S - to evaluate the anti-proliferation activity of ZNL-3. At 72-hour timepoint, ZNL-3 inhibited cell growth with a IC_50_ below 1 nM in L858R Ba/F3 cells, comparable to Osimertinib, whereas negative control ZNL-3r remarkably lost this activity (Figure 2A), supporting a covalent mechanism. Notably, mutation of cysteine 797 to a serine, ZNL-3 maintained its potency, while Osimertinib lost nearly 1000-fold activity (Figure 2B). The mutation of C775 to serine resulted in an opposite observation and the activity for ZNL-3 significantly diminished, reminiscent of a covalent dependency as reflected by ZNL-3r (Figure 2C). To examine target engagement, two biotin probes conjugating Osimertinib or ZNL-0013^15^ which can covalently label on cysteine 797 and cysteine 775 respectively in cell lysate were used to quantify the unmodified EGFR protein under pre-treatment of ZNL-3 in Ba/F3 cells. As exhibited in Figure 2D and Figure S2A, ZNL-3 achieved nearly complete engagement of EGFR at a concentration of 50 nM in both L858R and L858R/C797S cells. As an orthogonal assay, measuring downstream signaling on p-EGFR, p-ERK and p-AKT collectively highlighted on-target effects with a starting concentration from 10 to 50 nM (Figure 2E and Figure S2B).

**Figure 2.**
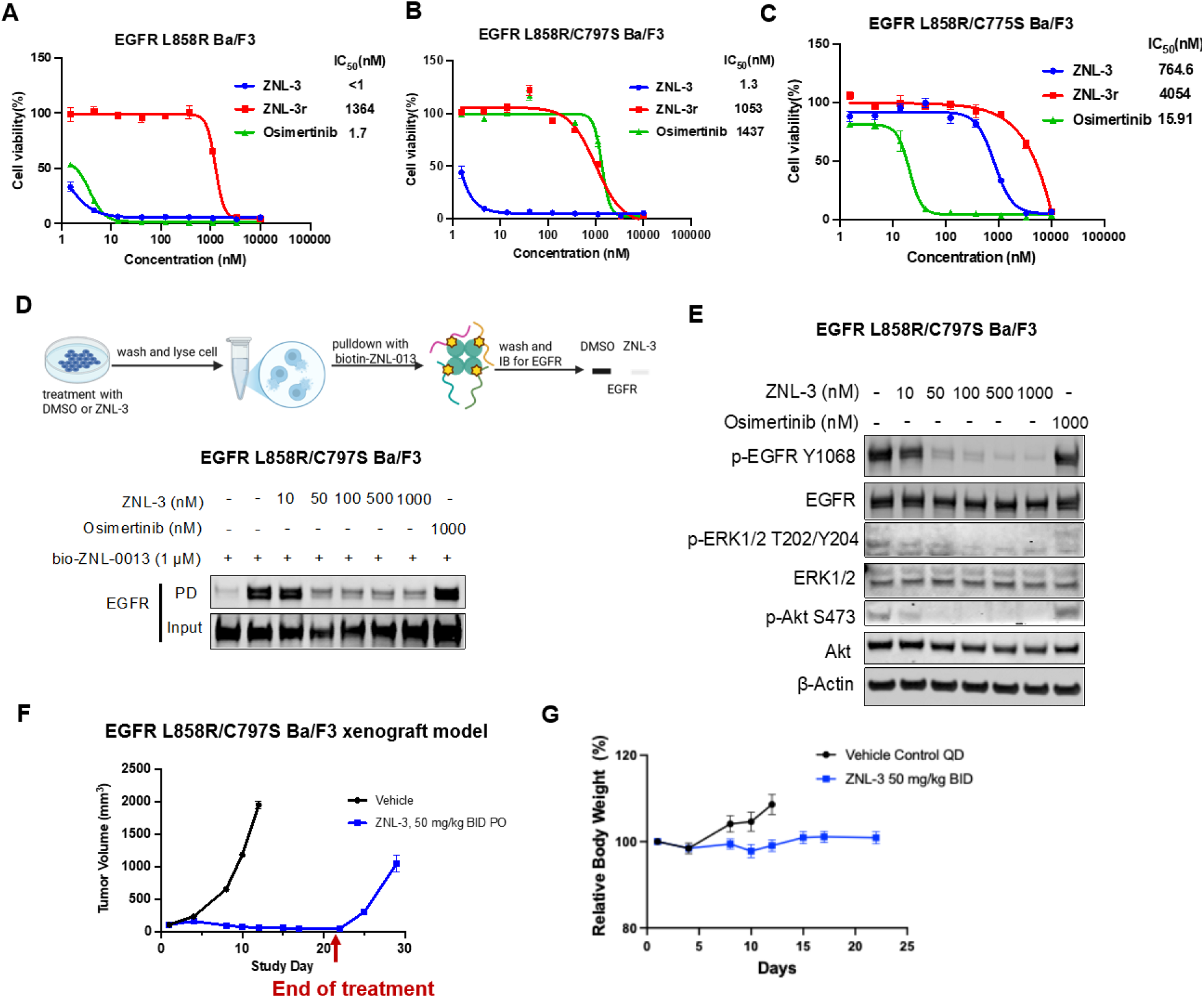
ZNL-3 covalently inhibits EGFR Cys797-Ser mutation and is efficacious in L858R/C797S xenograft mouse model **A.** Dose-response curves for ZNL-3, ZNL-3r and osimertinib in EGFR L858R Ba/F3 cells following 72 h of treatment. Cell viability was assessed with CellTiter-Glo. Data are presented as the mean±SEM. Representative data from three independent experiments are shown. **B.** Dose-response curves for ZNL-3, ZNL-3r and osimertinib in EGFR L858R/C797S Ba/F3 cells following 72 h of treatment. Cell viability was assessed with CellTiter-Glo. Data are presented as the mean ±SEM. Representative data from three independent experiments are shown. **C.** Dose-response curves for ZNL-3, ZNL-3r and osimertinib in EGFR L858R/C775S Ba/F3 cells following 72 h of treatment. Cell viability was assessed with CellTiter-Glo. Data are presented as the mean ±SEM. Representative data from three independent experiments are shown. **D.** Competitive pull-down assay in L858R/C797S Ba/F3 cells treated with ZNL-3 or Osimertinib at the indicated concentrations for 6 h. Cell lysates were incubated with bio-ZNL-0013 **E**. Effects of ZNL-3 on EGFR signaling pathway in EGFR L858R/C797S Ba/F3 after 6 h treatment. **F.** The *in vivo* antitumor efficacy of ZNL-3 in EGFR L858R/C797S-driven tumors (n =10 per group). Data are presented as the mean ±SEM. **G**. The body weight of EGFR L858R/C797S xenograft model (n =10 per group). Data are presented as the mean ±SEM.

Supported by biochemical and cellular data, ZNL-3 was advanced to a mouse pharmacokinetic (PK) study and an in vivo efficacy study with a xenograft model of L858R/C797S Ba/F3 cells. In the PK study, mice were dosed with 1 mg/kg and 10 mg/kg of IV and PO respectively. Analysis of mice plasma data demonstrated that ZNL-3 exhibits a moderate IV half-life of approximately 1 hour and achieved greater than 100% oral bioavailability (Table S3). As shown in Figure 2F, ZNL-3 treatment with a regiment of 50mg/kg twice daily (BID) resulted in significant tumor growth inhibition with regression during the treatment period. Tumor regrowth was only observed in the days after the drug was withdrawn. The compound was well tolerated with an average body weight loss of <3% (Figure 2F and 2G).

ZNL-3 also exhibits an antiproliferative EC_50_ of less than 100 nM against the established human tumor derived cancer cell lines H3255 and PC-9 which harbor L858R and Del-19, respectively (Figure 3A). Furthermore, ZNL-3 inhibits phosphorylation of p-EGFR, p-ERK and p-AKT with a dose-dependent manner in H3255 and PC9, with effects observed at concentrations between 10 to 50 nM (Figure 3B).

**Figure 3.**
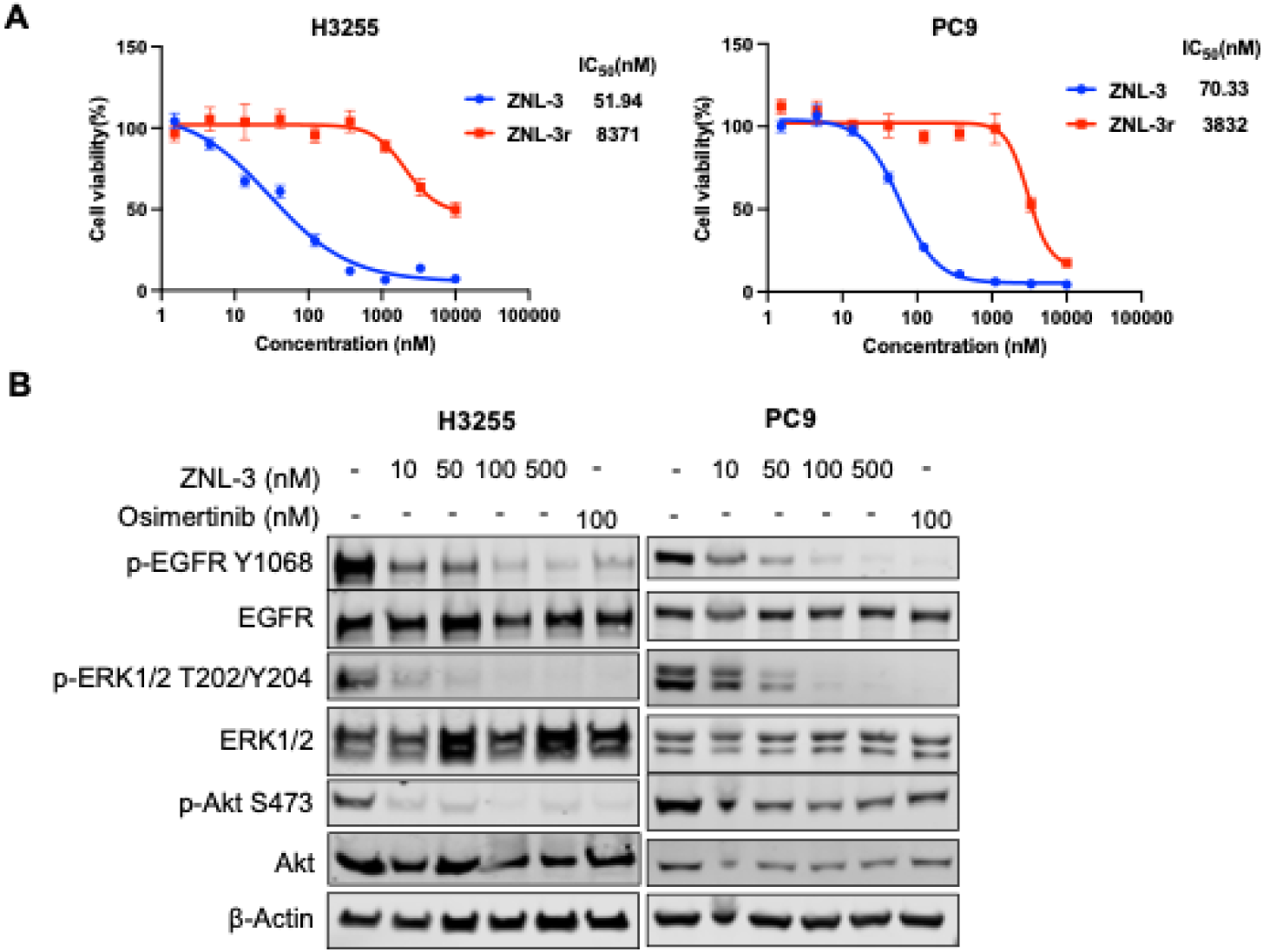
Antiproliferative and downstream inhibitory effect of ZNL-3 in human NSCLC cells. **A.** Dose-response curves for ZNL-3, ZNL-3r in H3255 and PC9 following 72 h of treatment. Cell viability was assessed with CellTiter-Glo. Data are presented as the mean±SEM. Representative data from three independent experiments are shown. **B.** Effects of ZNL-3 on EGFR signaling pathway in H3255 and PC9 after 6 h treatment. Representative data from three independent experiments are shown.

### Cysteine 775-Directed Approach for the Design of Bident EGFR Inhibitor YNW-1

The first generation bident EGFR inhibitor ZNL-0056 was designed from a molecule that labels front C797, and the optimization did not yield a potent covalent inhibitor through targeting C775 alone^15^. Therefore, ZNL-0056 is inadequate for inhibition of C797S mutant. The robust covalent engagement of C775 led us to envision that ZNL-3 might represent a novel starting point for a reverse approach to install a second acrylamide warhead for C797. With this hypothesis, we introduced acrylamide-equipped pyrrolidine, piperidine or phenyl ring to the sugar pocket and reduced the molecular size by eliminating the solvent-exposed phenylpiperazine group, leading to compounds ZNL-4, ZNL-5 and ZNL-6 (Figure 4A). All these designs maintained high anti-proliferative activity in L858R Ba/F3 cells with EC_50_s below 100 nM. To elucidate if these bident compounds can exert the activity through independent engagement on either cysteine, we evaluated their activity in Ba/F3 cells expressing either C797S or C775S mutations. Unfortunately, these compounds lost between 20 and 140-folds activity against both mutations with C797S having the greatest impact (Figure 4B and Figure S3A). These results suggest that covalent engagement at C775 is substantially less efficient than at C797. We hypothesized that improving the reversible binding for the residence time independent of covalency might achieve this balance. To improve the reversible binding, we re-introduced the hinge binding anilinopiperazine groups to generate ZNL-7 and ZNL-8 which exhibited sub 100 nM EC_50_ against both C797S and C775 mutations (Figure 4A, 4B and Figure S3A).

**Figure 4.**
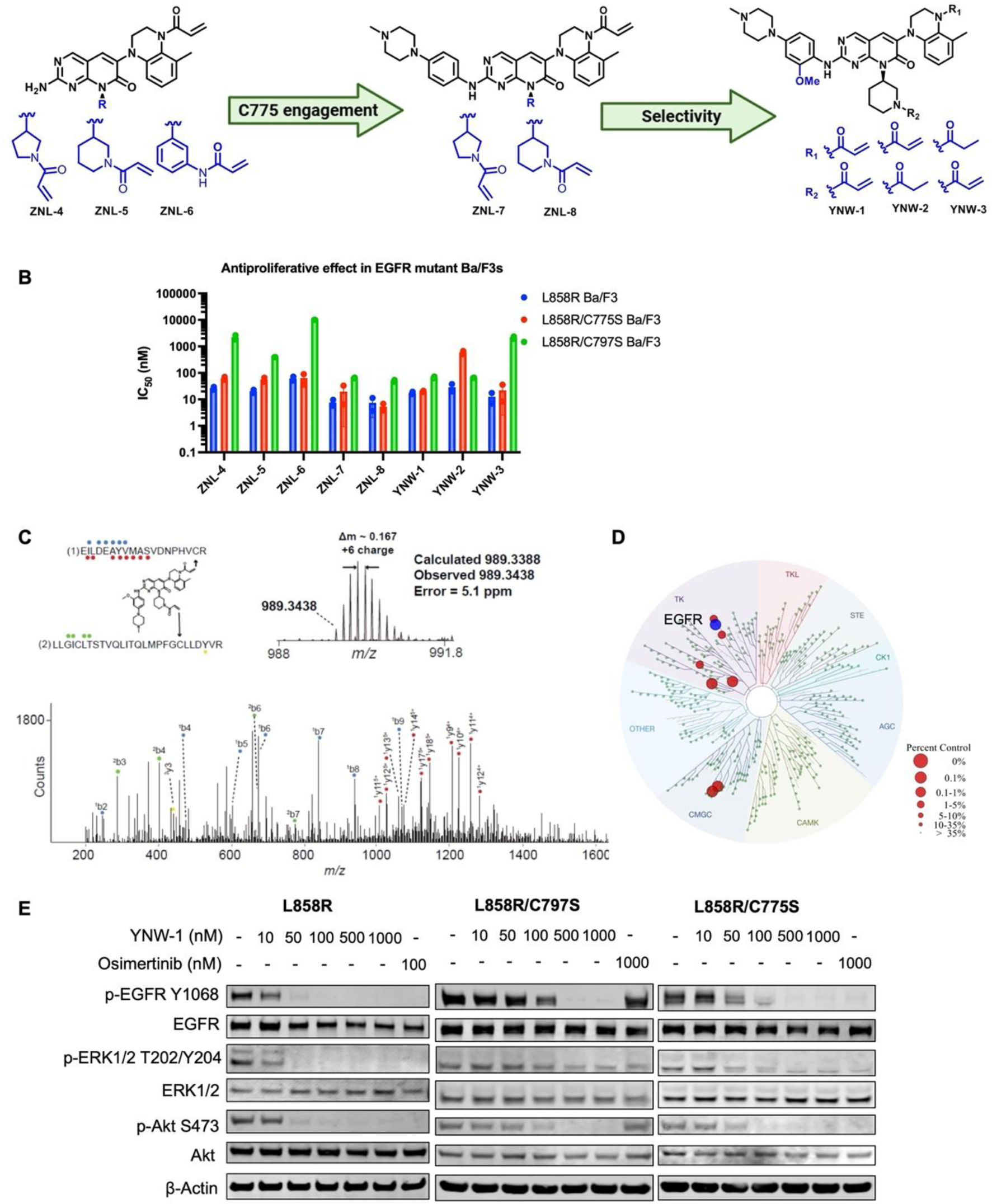
Cysteine 775-directed approach for the design of bident EGFR inhibitor YNW-1. **A.** Compound structures for EGFR dual cysteine-targeting molecules. **B**. The average IC50 values (mean±SEM) of the indicated compounds in EGFR L858R Ba/F3, EGFR L858R/C775S Ba/F3, and EGFR L858R/C797S Ba/F3 cells after 72 h treatment. Data are presented as the mean±SEM. Representative data from three independent experiments are shown. **C.** MS/MS spectrum for YNW-1 covalently modifying Cys775 and Cys797 of EGFR. Red and yellow circles indicate ions of type y for sequences marked (1) and (2), respectively. Blue circles indicate ions of b for sequence (1). *, carbamidomethyl cysteine, Δ, oxidized methionine. **D.** The KINOMEscan profiling of YNW-1 at 1µM. **E.** Effects of YNW-1 on EGFR signaling pathway in EGFR L858R Ba/F3, EGFR L858R/C775S Ba/F3, and EGFR L858R/C797S Ba/F3 after 6 h treatment. Representative data from three independent experiments are shown.

One of concerns for bident compound is the potential for increased promiscuity due to the presence of a second electrophile warhead. To assess this, we conducted KinomeScan profiling. At 1 µM, ZNL-8 (Figure S3B) demonstrated moderate selectivity, with 37 off-targets based on the fraction of kinases inhibited by >90% (Table S4). YNW-1 was designed to incorporate a 2-methoxy group to the aniline which is a well-known kinase selectivity enhancing modifications (Figure 4A). To confirm the covalent bond formation with Cys797 and Cys775 by YNW-1, we conducted mass-spectrometry with trypsin digestion of purified recombinant EGFR^L858R^ kinase domain treated by YNW-1, identified cross-linked peptide corresponding to Cys797 and Cys775, supporting the conclusion that YNW-1 forms simultaneous, intramolecular covalent bonds with both residues (Figure 4C). YNW-1 demonstrated a highly improved selectivity profile across the whole kinome and only 7 kinases were detected as potential off-targets (>90% inhibition) (Figure 4D and Table S5), none of which contains a targetable cysteine (Table S6) in the kinase domain. Furthermore, YNW-1 exhibited robust mutant EGFR inhibition with an apparent IC_50_ below 10 nM (Figure 4B and S3C). The anti-proliferative EC_50_ in Ba/F3 remained potent (19 nM and 62 nM) regardless of the mutations on either C775 or C797 (Figure S3A). but became inactive once the simultaneous mutations on both cysteines occurred (Figure S3D). This equal independence on cysteines was further evidenced by the evaluation of two control compounds YNW-2 and YNW-3 which were designed as mono-covalent inhibitors targeting C775 and C797, respectively (Figure 4A). They unambiguously demonstrated the necessity of cysteine-targeting as its anti-proliferation activity was significantly diminished once its designated cysteine was mutated (Figure 4B and Figure S3A). Consistent with anti-proliferation activity, the downstream singling including p-EGFR/ERK and AKT were inhibited by 50-100nM of YNW-1 with background of L858R, L8585/C797S and L858R/C775S EGFR mutants (Figure 4E).

### YNW-1 has Broad Activity against EGFR Mutants and Delayed Acquired Mutations

The unique mechanism of action for YNW-1 prompted us to investigate two potential advantages: resilience to a broad spectrum of EGFR mutants and suppression of acquired point mutation. First, we tested YNW-1 in Ba/F3 cells engineered with a Del19 background. As shown in Figure 5A, unveiled a similar activity across Del19, Del19/C775S and Del19/C797S. Second, the compound was active against three exon 20 insertion mutations (insASV, insSVD and insFQEA) which universally displayed sensitivity with an EC_50_ ranging from 100 nM to 250 nM (Figure 5B). Other rarely occurred mutations including E790K, L861Q and G719A/E709K were also susceptible to YNW-1 (Figure 5C). While del18, G719A, S768I and G719A/T790M are less susceptible to YNW-1(Figure S4). Notably, YNW-2 and YNW-3 showed reduced activity across these mutations, consistent with the enhanced potency conferred by the dual covalent-bonding mechanism of YNW-1.

**Figure 5.**
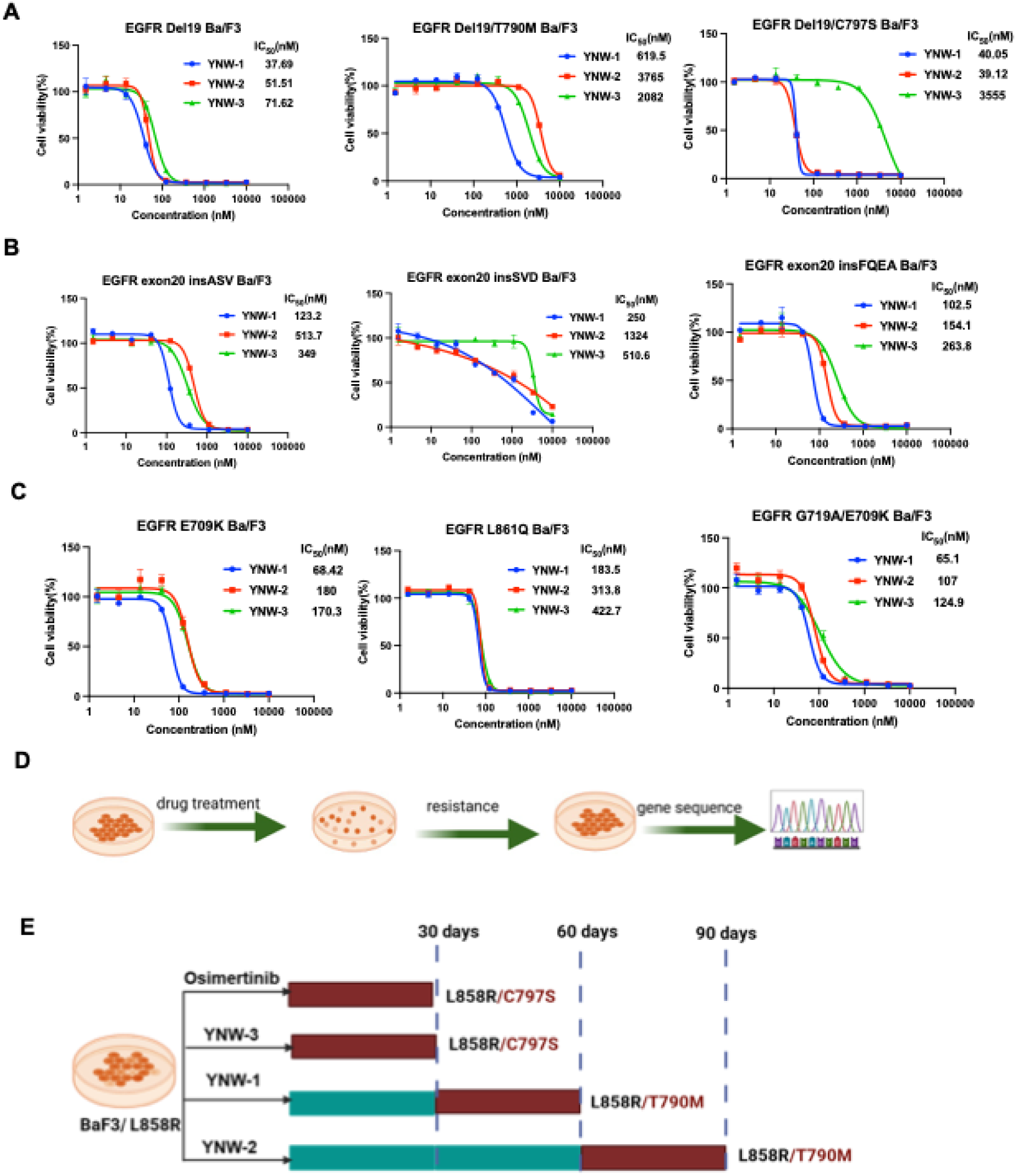
YNW-1 has broad activity against EGFR mutants and delayed acquired mutations **A.** Dose-response curves for YNW-1, YNW-2, YNW-3 in EGFR Del19 Ba/F3, EGFR Del19/T790M Ba/F3 and EGFR Del19/C797S Ba/F3 after 72 h treatment. Cell viability was assessed with CellTiter-Glo. Data are presented as mean±SEM. Representative data from three independent experiments are shown. **B.** Dose-response curves for YNW-1, YNW-2, YNW-3 in EGFR exon20 insASV Ba/F3, EGFR exon20 insSVD Ba/F3 and EGFR exon20 insFQEA Ba/F3 after 72 h treatment. Cell viability was assessed with CellTiter-Glo. Data are presented as the mean±SEM. Representative data from three independent experiments are shown. **C.** Dose-response curves for YNW-1, YNW-2, YNW-3 in EGFR E709K Ba/F3, EGFR L861Q Ba/F3 and EGFR G719A/E709K Ba/F3 after 72 h treatment. Cell viability was assessed with CellTiter-Glo. Data are presented as mean±SEM. Representative data from three independent experiments are shown. **D.** Schematic illustration of chronic exposure for YNW-1 acquired resistant mutation **E.** Mutations found in EGFR kinase domain for YNW-1, YNW-2, YNW-3 and osimertinib in EGFR L858R Ba/F3-resistant cells.

We next performed a chronic drug-exposure study, gradually escalating the concentrations of YNW-1, YNW-2, and YNW-3 until cells were able to proliferate normally at 500 nM. To assess whether resistance mechanisms differ between bident and monovalent inhibitors (Figure 5D). We harvested persistent cells for sequence analysis on EGFR after one-, two-and three-month continuous drug exposure. As shown in Figure 5E and S5, osimertinib and YNW-3 rapidly induced the C797S mutation within one month, whereas cells treated with the bident inhibitors YNW-1 and YNW-2 remained in the wild-type state. Interestingly, the bident compound began to induce the gatekeeper T790M mutation instead of the mutation on cysteines in two months. However, this mutation was absent with YNW-2 at this time point but emerged only after three-month treatments. The results indicate that targeting C775 might have a slow onset of acquired resistance through EGFR mutation compared to Osimertinib. The underline mechanism is under investigation.

## Discussion

The emergence of resistance to the front-line therapy Osimertinib through both EGFR-dependent and -independent mechanisms remains as an unmet medical need. Although multiple drug-resistant EGFR mutations have been identified, the C797S mutation accounts for a substantial proportion of patients who relapse following Osimertinib treatment. ZNL-3, a first-in-class cysteine 775-targeted covalent inhibitor for EGFR, has demonstrated promising therapeutic potential to overcome this resistance. Both *in vitro* and *in vivo* efficacy study suggested that ZNL-3 potently inhibited mutant EGFR signaling and drove tumor regression with good tolerability. Compared to current drug discovery efforts focusing on allosteric, reversible inhibitors and degraders, ZNL-3 offers a completely new direction for clinical translation with its robust antitumor activity, selectivity, and efficiency in the drug development process. Its mode action differentiates it from reversible attempts by retaining the remarkable efficacy of covalent inhibitors for mutant EGFR NSCLC demonstrated by Osimertinib while addressing the C797S resistance. While ZNL-3 serves as a promising lead compound targeting the L858R and L858R/C797S mutations, the activity suffers from the T790M mutation, displaying IC_50_ values above 1 μM in Ba/F3 cells (Figure S6A), and losing activity in H3255GR and DFCI52 cells that harbor the T790M gatekeeper mutation (Figure S6B). Further optimization of ZNL-3 to overcome this limitation is under way. One the other hand, T790M in the real world mEGFR NSCLC patient population is diminishing due to the first line treatment of Osimertinib, clinical candidates stemmed from ZNL-3 can be readily developed to address the drug-resistant mutants as the second line therapy. In addition, it can be used in combination with Osimertinib to delay the occurrence of drug resistance even though both compounds are mutually exclusive when binding to an EGFR protein, since a double mutation (C797S/C775S) in cis would be required to confer resistance to the combination therapy.

The bident compound built upon ZNL-3 provides a new strategy and a breakthrough in the discovery of new modality. YNW-1 is the first bident molecule disclosed so far with a nearly equal efficiency on both cysteines despite drastically different protein microenvironment, demonstrating small molecules can mitigate different nucleophilic reactivity within one molecule, a first in its class. The unambiguous characterization of YNW-1 with high selectivity, broad activity across EGFR mutants and more importantly, slow onset of acquired point mutation compared to Osimertinib is intriguing. Here, we observed that the gatekeeper mutation T790M emerged rather than either of cysteines for bident compound YNW-1 which is consistent with the observed EC_50_ on L858R/T790M (136 nM) (Figure S7A and S7B) compared to L858R/C797S (62 nM) and L858R/C775S (19 nM) (Figure S3A). It points to the direction to further optimize the inhibition of T790M for YNW-1chemical series. We believe this is an addressable issue with medicinal chemistry effort rather than intrinsic weakness of the modality. Since single mutation on either cysteine is incapable of causing resistance, the point mutation on EGFR is predicted as a minor resistance mechanism for a potent L858R/T790M bident inhibitor. Finally, the observation of delayed mutagenesis on T790M for YNW-3 was encouraging and might imply a potential therapeutic superiority to Osimertinib. It may also pose advantage over Osimertinib combination therapy by less drug load to reduce any potential side effects. Unquestionably, the development of an optimized version of YNW-1 or ZNL-3 that overcomes T790M will be a major next step and offers a new strategy to manage mEGFR driven non-small cell lung cancer, which is an on-going effort in our laboratory and will be reported in a following publication.

## Methods

### Crystallization and Structure Determination

Wild-type EGFR (residues 696-1022) was purified as described for other variants. Protein at ∼3 mg/mL was incubated with 0.5 mM ZNL-1 and incubated for 1 hr at room temperature. Sitting drop screens were set with drops containing 0.2 µL protein and 0.2 µL reservoir solution adjacent to 50 µL reservoir solution. Screens were incubated at room temperature and crystals of two morphologies (rods and diamonds) observed after ∼1 month in condition C1 (3.5 M sodium formate pH 7.0) of the Index-HT screen (Hampton Research). Crystals were harvested directly from the screening tray and cryoprotected in a solution containing 3.5 M sodium formate (pH 7.0) and 25% ethylene glycol prior to flash freezing in liquid nitrogen.

Diffraction data were collected under a cryostream at NE-CAT beamlines at the Advanced Photon Source at Argonne National Lab. Crystals with rod morphology diffracted significantly better than diamond crystals. Data were processed using Dials via xia2 and SBGrid.^17,18^ The structure was phased via molecular replacement with PDB 2GS2 and the N-lobe of the kinase manually rebuilt due to large rotational conformational changes.^19^Refinement was performed using Phenix and manual model building in Coot.^20,21^ Custom ligand restraints were generated using eLBOW in Phenix with AM1 quantum mechanical optimization.^22^ The resulting structure has been deposited in the Protein Data Bank (PDB) with the accession code 00009PWI.

### Mass spectrometry analysis

EGFR protein was treated with a 5-fold molar excess of indicated probe or DMSO for 2 hours at 30 °C in 50 mM HEPES buffer 100 mM NaCl pH 7.5. Reactions were analyzed by LC-MS with a Shimadzu HPLC system (Shimadzu, Marlborough, MA) interfaced to an LTQ ion trap mass spectrometer (Thermofisher Scientific, San Jose, CA) operated in profile mode (m/z 300-2000, spray voltage = 5 kV). Proteins were desalted for 1 min with solvent A before eluting to the mass spectrometer with a ballistic gradient (0-100% B in 1 minute, A=0.1% formic acid in water, B=0.1% formic acid in acetonitrile). Raw data were deconvoluted with Magtran 1.03b software (PMID: 9879360).

To identify sites of dual warhead crosslinking, protein was treated as described above and then reduced (10 mM dithiothreitol, 37 °C, 30 minutes), alkylated (22.5 mM iodoacetamide, 30 minutes, room temperature, protected from light), and digested overnight with trypsin (1:20 trypsin:EGFR; PROMEGA, Madison, WI) at 37 °C. Peptides were loaded onto EVO tips and analyzed with a 20-SPD method on an EVO Sep One (column=75 µm I.D. packed with 15 cm 1.5 µm Dr. Maisch C18 [ESI Source Solutions, Woburn, MA] with integrated emitter tip (PMID: 19331382)) coupled to a Bruker timsTOF Pro2 mass spectrometer operated in DDA-PASEF mode. Dual warhead targets were identified by calculating *m/z* values for expected cross-linked peptides using the PepCalc tool in mzStudio (PMID: 28763045). This tool was then used to make extracted ion chromatograms (XICs) for expected charge states of the cross-linked peptides (typically +4-+7). We then examined MS/MS spectra within XIC elution profiles that were identified with the “Find Precursors” function of PepCalc. We identified the cross-linked peptides by mapping y and b ions onto MS spectra as shown in Figure S4E.

### Cell culture

H3255(RRID: CVCL_6831) and H3255GR were cultured in ACL-4 media^23^ containing 10% fetal bovine serum (GeminiBio, Cat #100-106) and 1% Penicillin/Streptomycin. The EGFR mutant Ba/F3 cells were generated and characterized as described previously^10^, and cultured in in RPMI media (Life technologies, Cat# 11875119) containing 10% fetal bovine serum and 1% Penicillin/Streptomycin. All the cell lines were cultured at 37°C in 5% CO_2_ humidified air and tested for mycoplasma negative.

### Immunoblotting and Antibodies

Cells were lysed in RIPA buffer (150 mM NaCl, 1.0% IGEPAL® CA-630, 0.5% sodium deoxycholate, 0.1% SDS, 50 mM Tris, pH 8.0) (Sigma, Cat# R0278) with protease inhibitor and phosphatase inhibitor (Roche). The protein concentrations were measured by BCA analysis (Thermo Fisher Scientific, Cat # PI23225). Equal amounts of protein were resolved by 4-12% Tris-Base gels (Life Technologies) and then transferred to the Immuno-Blot PVDF membrane (BioRad, cat # 1620177). Proteins were probed with appropriate primary antibodies at 4 ℃ overnight and then with IRDye®800-labeled goat anti-rabbit IgG (LICOR Biosciences, cat # 926-32211, RRID: AB_621843), IRDye®800-labeled goat anti-mouse IgG (LICOR Biosciences, cat # 926-32210, RRID: AB_621842) or IRDye 680RD goat anti-Mouse IgG (LICOR Biosciences, Cat # 926-68070, RRID: AB_10956588) secondary antibodies at room temperature for 1 hour. The membranes were detected on the Odyssey CLx system.

Antibodies used in this study include anti-following proteins: EGFR (Cell signaling Technology, 4267S, 1:1000, RRID: AB_2895042), p-EGFR (Tyr1068) (Cell signaling Technology, #3777S, 1:1000, RRID: AB_2096270), ERK1/2 (Cell signaling Technology, 4696S, #1:0000, RRID: AB_390780), p-ERK1/2 (Cell Signaling Technology, 4370S, 1;1000, RRID: AB_2315112), Akt (Cell signaling Technology, # 9272L, 1:1000, RRID: AB_329827), p-Akt (Cell Signaling Technology, #4060S, 1;1000, RRID: AB_2315049) and β-Actin (Cell Signaling Technology, #3700, 1:1000, AB_10828322).

### Antiproliferation Assay

Cells were seeded at the density of 1000 cells/well for 384-well plates. For adherent cell lines, cells were cultured for 16 hours before compounds were added into the media for 72 hours treatment. Cell viability was determined by using CellTiter-Glo (Promega #G7571) according to the manufacturer’s instructions, measuring luminescence using an FLUOstar Omega plate reader (BMG LABTEC). Dose-response curves were generated using non-linear regression curve fit in GraphPad Prism (RRID: SCR_002798).

### Cellular Target Engagement Assays

After 6 hours treatment, cells were pelleted, washed with PBS once and lysed with IP lysis buffer (25 mMTris-HCl pH 7.4, 150 mM NaCl, 1 mM EDTA, 1% NP-40 and 5% glycerol) (Thermo Fisher Scientific, Cat#87788) containing protease/phosphatase inhibitor cocktail (Roche). The protein concentrations were measured by BCA analysis (Pierce). Cell lysates were incubated with 1 μM of biotin conjugated probe (bio-Osimertinib or ZNL-0130) at 4 ℃ overnight, and incubated for 3 more hours at room temperature. Lysates with probe were then incubated with streptavidin beads (Thermo Fisher, #20349) for 2 hours at 4 ℃. The protein-probe complexes on the beads were then subjected to immunoblotting.

### Protein Expression and Purification

The human EGFR kinase domain (residues 696–1022) harboring the L858R, L858R/T790M, L858R/C775S, or L858R/C797S mutations was cloned into a pFastBac vector with an N-terminal His-GST tag containing a TEV protease cleavage site. Recombinant proteins were expressed in Spodoptera frugiperda (SF9) insect cells using baculovirus generated via the pFastBac system (Invitrogen). SF9 cells were cultured to a density of approximately 1.5-2 × 10⁶ cells/mL and infected with 1%-2% (v/v) virus stock. After 68-72 hours of incubation, cells were harvested via centrifugation and the resulting cell pellets were stored at −80 °C.

For protein extraction, frozen cell pellets were resuspended and lysed in buffer A comprised of 50 mM Tris-HCl, pH 8.0, 500 mM NaCl, 5% glycerol, 0.5 mM Tris(2-carboxyethyl) phosphine [TCEP]). The resulting solution was lysed via sonication. Lysates were clarified by ultracentrifugation at >200,000 × g for 1 hour at 4 °C. The resulting lysate was filtered with a 2um glass filter before being applied to Ni-NTA agarose resin by gravity flow. The resin was washed with buffer A supplemented with 40 mM imidazole and the bound protein was eluted with buffer A containing 200 mM imidazole.

To cleave the His-GST tag, a stoichiometric excess of in-house purified TEV protease was added to the eluate, and the mixture was dialyzed overnight at 4 °C in ∼1L of buffer A. Following protease digestion, the sample was re-applied to Ni-NTA resin to remove the cleaved tag and uncleaved protein. The kinase domain was then further purified by size-exclusion chromatography on a Superdex 200 (S200) column (Cytiva) equilibrated in buffer A. Fractions containing the target protein with a purity of ≥95% (confirmed by SDS-PAGE) were pooled and concentrated for subsequent experiments.

### Kinase Inhibition Assay

Kinase inhibition assays were conducted using the HTRF® KinEASE™ TK assay kit (Cisbio) following the manufacturer’s instructions. Small-molecule inhibitors were prepared from 10 mM DMSO stock solutions and dispensed into black 384-well plates using a D300e Digital Dispenser (Hewlett-Packard), normalized to a final DMSO concentration of 1% (v/v) in all wells.

Purified EGFR kinase domain was diluted to a final concentration of 0.2 nM in assay buffer and dispensed into the plates using a Multidrop Combi dispenser (Thermo Fisher Scientific). Plates were incubated at room temperature for 60 minutes to allow pre-equilibration with the inhibitors. Reactions were initiated by adding ATP to a final concentration of 100 µM and incubated for an additional 30 minutes at room temperature.

Reactions were quenched with the detection reagent provided in the KinEASE kit. Fluorescence resonance energy transfer (FRET) signals were measured at 620 nm and 665 nm using a PHERAstar microplate reader (BMG LABTECH). Data were analyzed in GraphPad Prism 10 and fitted to a three-parameter dose–response model to determine inhibitor potency.

### Efficacy study with mice

The efficacy study was conducted at Dana-Farber Cancer Institute with the approval of the Institutional Animal Care and Use Committee in an AAALAC accredited vivarium.

The L858R/C797S Ba/F3 cells, grown in RPMI-1640 with 10% FBS, were harvested and 2 million cells with 30% Matrigel implanted subcutaneously in female NOD.Cg-*Prkdc^scid^ Il2rg^tm1Wjl^*/SzJ (NSG) mice from The Jackson Laboratories (ME). Tumors were allowed to establish to 104.9 ± 11.9 mm^3^ in size before randomization into treatment groups (Studylog, CA software) with n=9/group as: vehicle control (10% N-methyl-2-pyrrolidone and 90% PEG 300) or ZNL-3 at 50 mg/kg administered twice daily. Mice were treated orally for 21 days. Tumor volumes were determined from caliper measurements by using the formula, Tumor volume = (length x width^2^)/2. Tumor volumes and body weights were measured twice weekly

### Author Contributions

N.S.G., T.Z, J.C, Z.L. conceived this work; Chemistry, the synthesis of the compounds and structure-activity analysis by Z.L. Y.W. and D.G.; Coordinating with outsourced chemistry, scale-up of key compounds and physicochemical property measurement by Z.L and Y.W., Cellular experiments and data analysis by J.J., F.H.G., and L.M.B; Mass spectrometry experiments by S.B.F and I.T.; Protein expression, biochemical assay and crystallography by T.S.B., S.J.C. and I.K.S; *In vivo* efficacy study by D.S. and P.C.G.; Scientific guidance on experimental setup and data interpretation: N.S.G., T.Z., J.C., J.A.M., M.J.E., P.A.J. Writing original draft by T.Z.; Draft reviewing and editing by N.S.G., J.J., J.C., Z.L. and Y.W..

### Declarations

The authors declare the following competing financial interest(s): N.S.G. is a founder, science advisory board member (SAB), and equity holder in Syros, C4, Allorion, Lighthorse, Voronoi, Inception, Matchpoint, CobroVentures, GSK, Shenandoah (board member), Larkspur (board member), and Soltego (board member). The Gray laboratory receives or has received research funding from Novartis, Takeda, Astellas, Taiho, Jansen, Kinogen, Arbella, Deerfield, Springworks, Interline, Sanofi and Simcere. T.Z. is a scientific founder, equity holder, and consultant of Matchpoint, equity holder of Shenandoah. P.A.J. has received consulting fees from AbbVie, Accutar Biotech, Allorion Therapeutics, AstraZeneca, Bayer, Biocartis, Boehringer Ingelheim, Chugai Pharmaceutical Co., Daiichi Sankyo, Duality, Eisai, Eli Lilly, Frontier Medicines, Hongyun Biotechnology, Merus, Mirati Therapeutics, Monte Rosa, Novartis, Nuvalent, Pfizer, Roche/Genentech, Scorpion Therapeutics, SFJ Pharmaceuticals, Silicon Therapeutics, Syndax, Takeda Oncology, Transcenta, and Voronoi; sponsored research support from AstraZeneca, Boehringer Ingelheim, Daiichi Sankyo, Eli Lilly, Puma Technology, Revolution Medicines, and Takeda Oncology; post-marketing royalties from a DFCI owned patent on EGFR mutation licensed to Lab Corp; and has stock in Gatekeeper Pharmaceuticals. J.A.M. is a founder, equity holder, and advisor to Entact Bio, serves on the SAB of 908 Devices, and receives or has received sponsored research funding from Vertex, AstraZeneca, Taiho, Springworks, and TUO Therapeutics. J.C. is a scientific co-founder M3Bioinformatics& Technology Inc., and consultant and equity holder for Matchpoint, Soltego and Allorion. J.C. had received sponsored research support from Springworks and Deerfield. M.J.E. has received sponsored research support from Novartis, Takeda, Springworks, and Sanofi and consulting income from Novartis. All other authors declare no competing interests.

## Acknowledgments

We thank SpringWorks Therapeutics and Antidote Health Foundation for Cure of Cancer for financial support. We thank Stephen Gwaltney from SpringWorks Therapeutics for the consultation on the compound design. This work is based upon research conducted at the Northeastern Collaborative Access Team beamlines (P30 GM124165, P41 GM103403) utilizing resources of the Advanced Photon Source at the Argonne National Laboratory (DE-AC02-06CH11357).

## Supplementary Information

**Figure S1.**
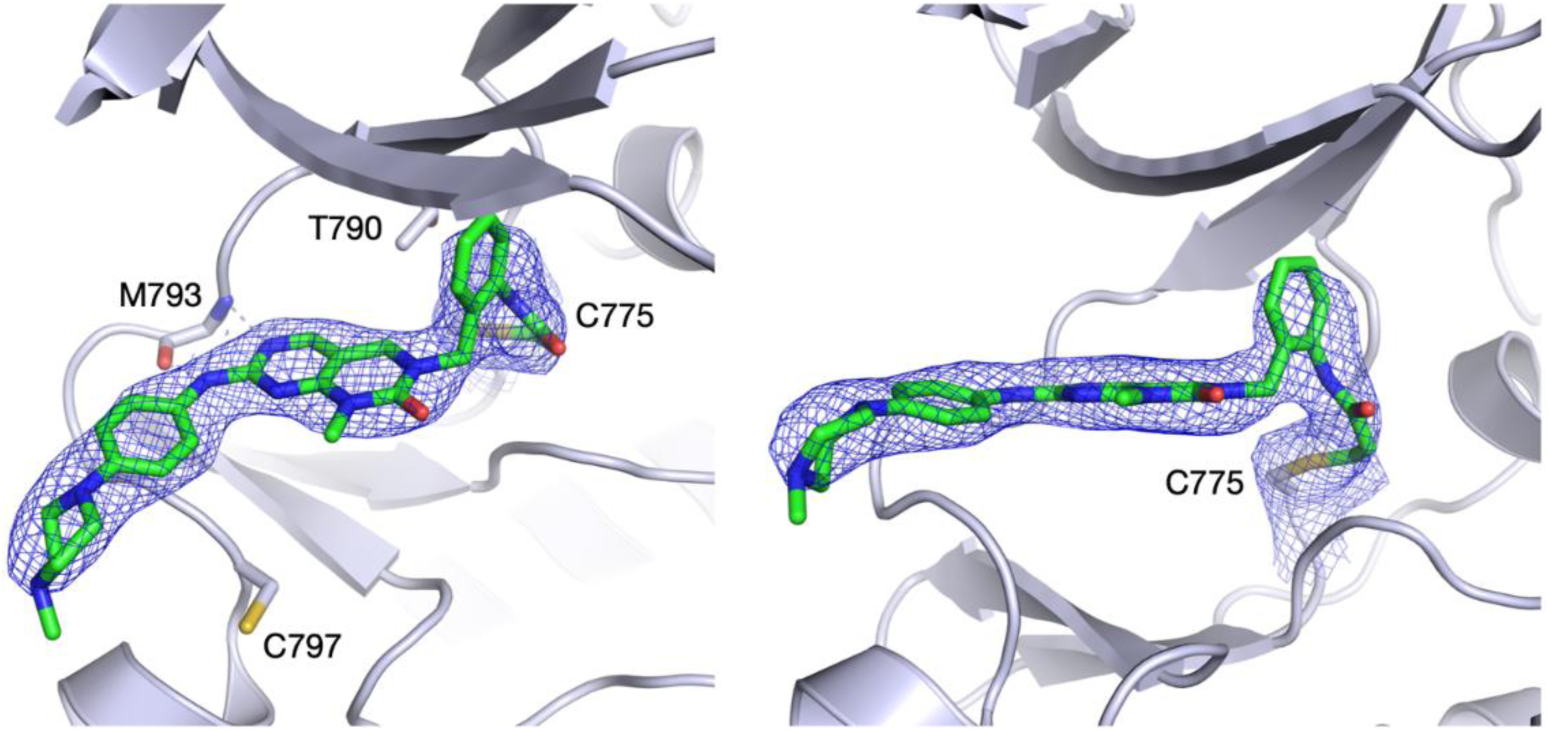
Electron density for ZNL-1 bound to EGFR. Mesh shown is 2F_o_-F_c_ contoured at 1 sigma. Density shown includes covalent adduct with C775.

**Figure S2.**
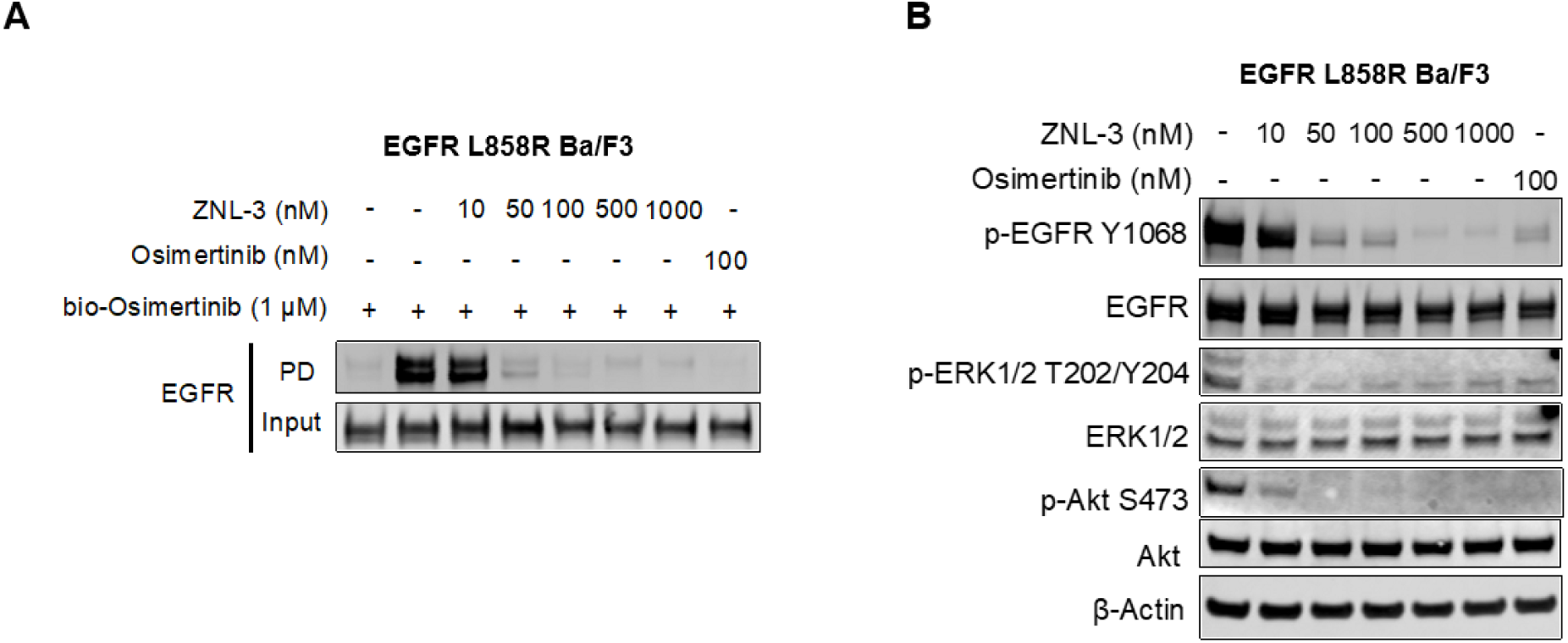
**A**. Competitive pull-down assay in EGFR L858R Ba/F3 cells treated with ZNL-3 or Osimertinib at the indicated concentrations for 6 h. Cell lysates were incubated with bio-osimertinib. **B.** Effects of ZNL-3 on EGFR signaling pathway in EGFR L858R Ba/F3 after 6 h treatment. Representative data from three independent experiments are shown

**Figure S3.**
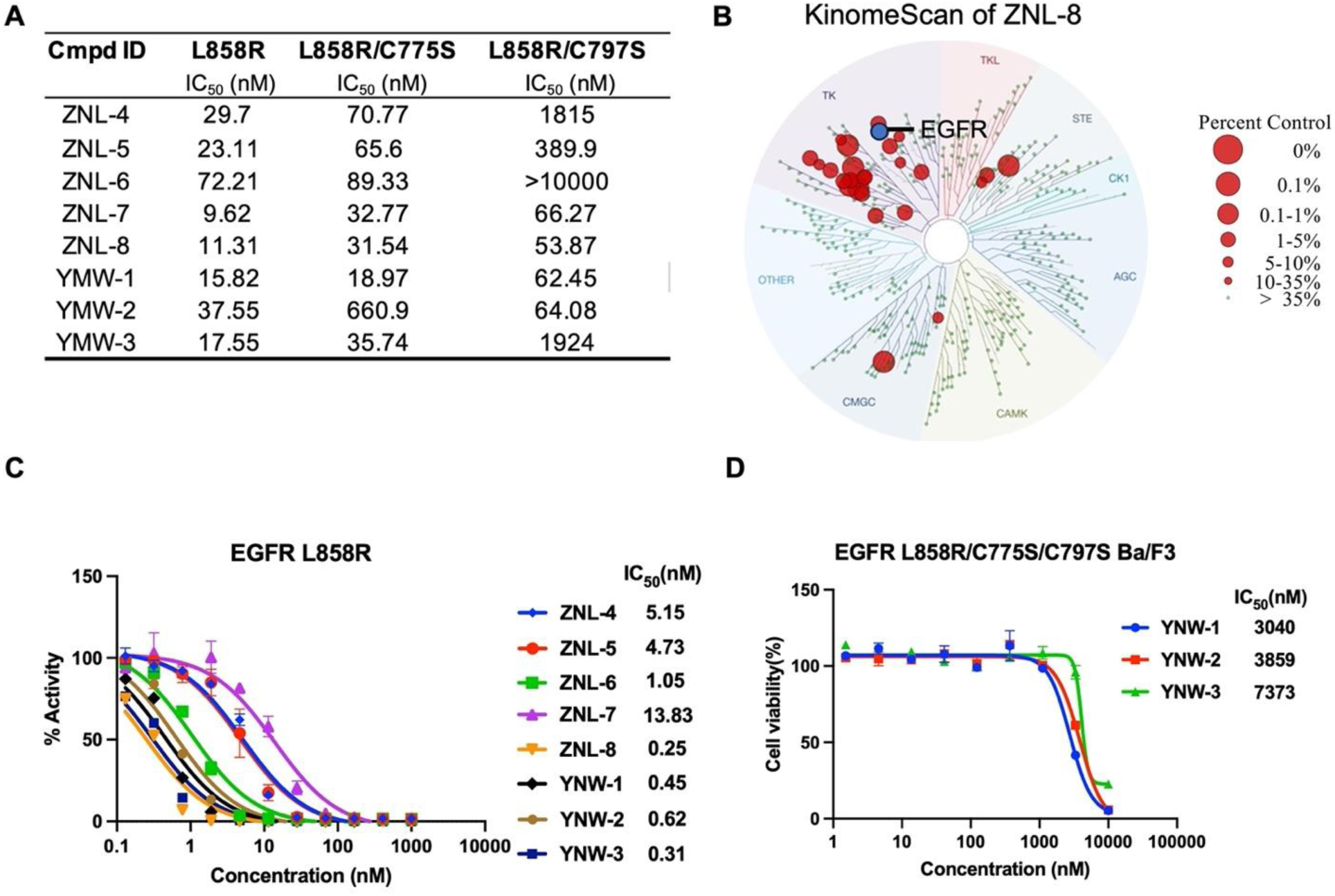
**A.** IC_50_s of ZNL series compounds and YNW-1,2,3 in EGFR L858R, EGFR L858R/C775S and L858R/C797S Ba/F3. Cell viability was assessed with CellTiter-Glo. Representative data from three independent experiments are shown. **B.** The KINOMEscan profiling of ZNL-8 at 1µM. **C.** Biochemical IC50s for the indicated compounds against EGFR L858R. **D.** Dose-response curves for YNW-1, YNW-2 and YNW-3 in EGFR L858R/C775S/C797S Ba/F3 after 72 h treatment. Cell viability was assessed with CellTiter-Glo. Data are presented as the mean±SEM. Representative data from three independent experiments are shown.

**Figure S4.**
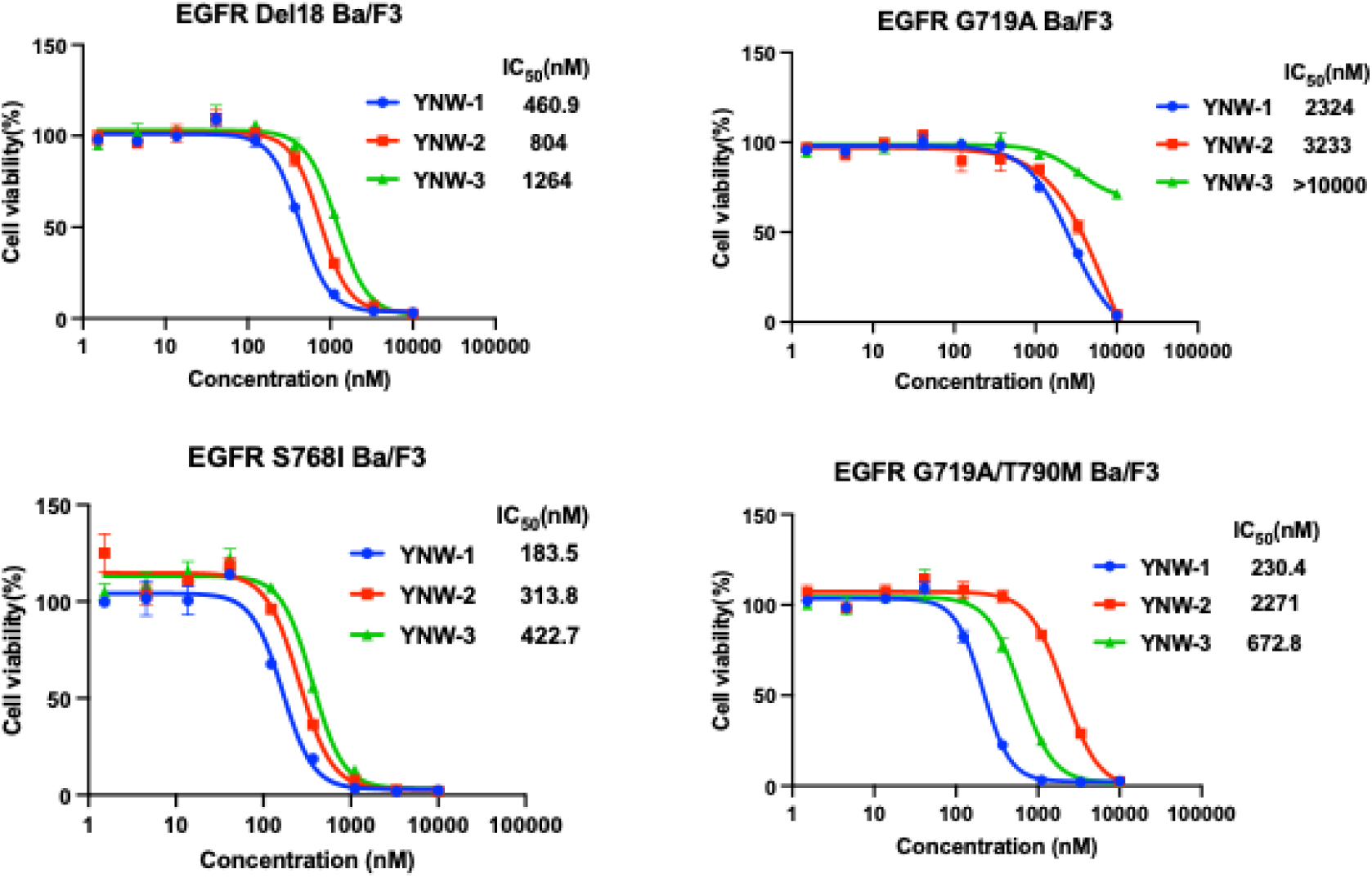
Dose-response curves for YNW-1, YNW-2 and YNW-3 in the indicated EGFR rare mutant Ba/F3 cells after 72 h treatment. Cell viability was assessed with CellTiter-Glo. Data are presented as the mean±SEM. Representative data from three independent experiments are shown.

**Figure S5.**
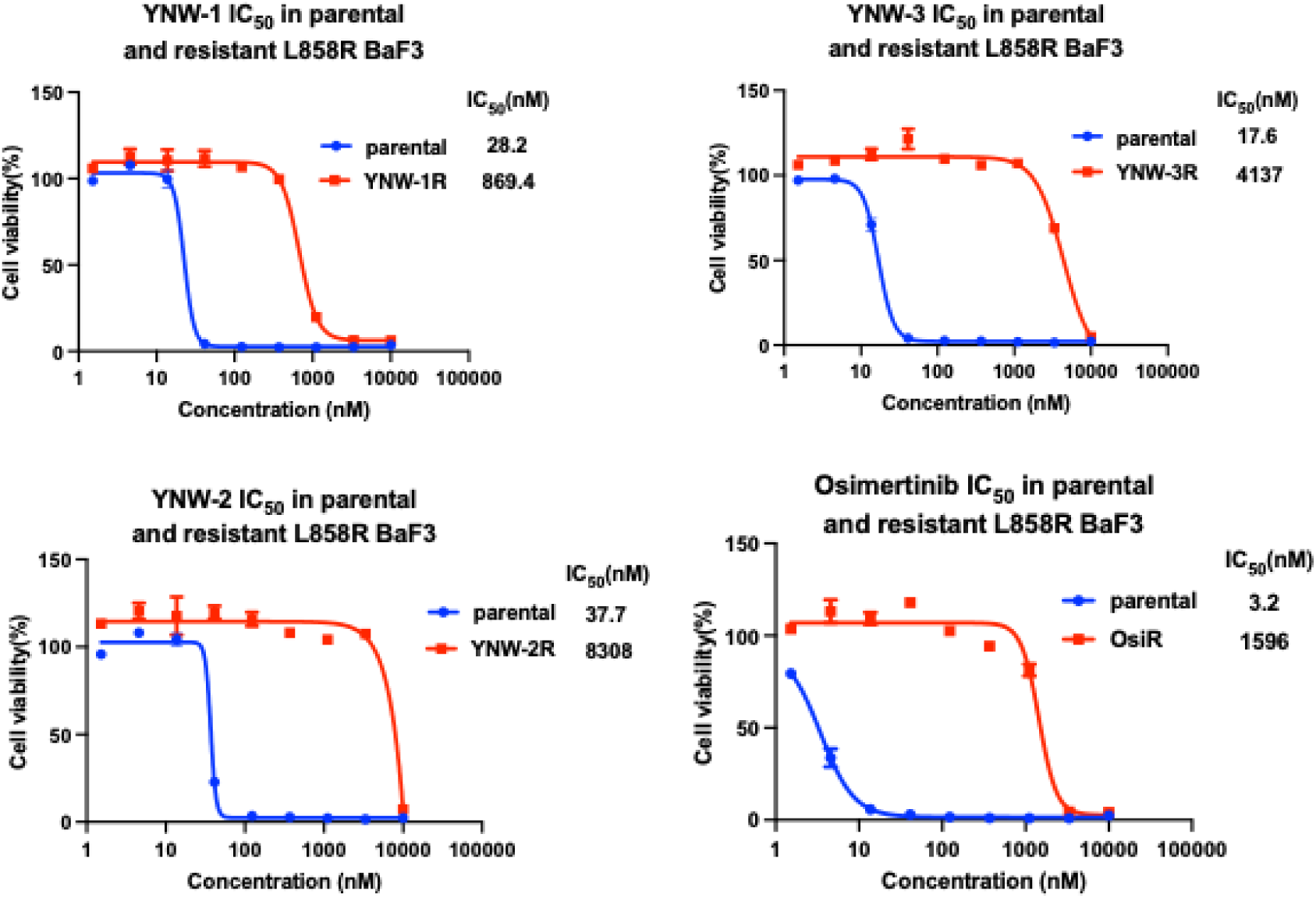
Dose-response curves for YNW-1, YNW-2, YNW-3 and osimertinib in parental or resistant EGFR L858R Ba/F3 after 72 h treatment. Cell viability was assessed with CellTiter-Glo. Data are presented as the mean±SEM. Representative data from three independent experiments are shown.

**Figure S6.**
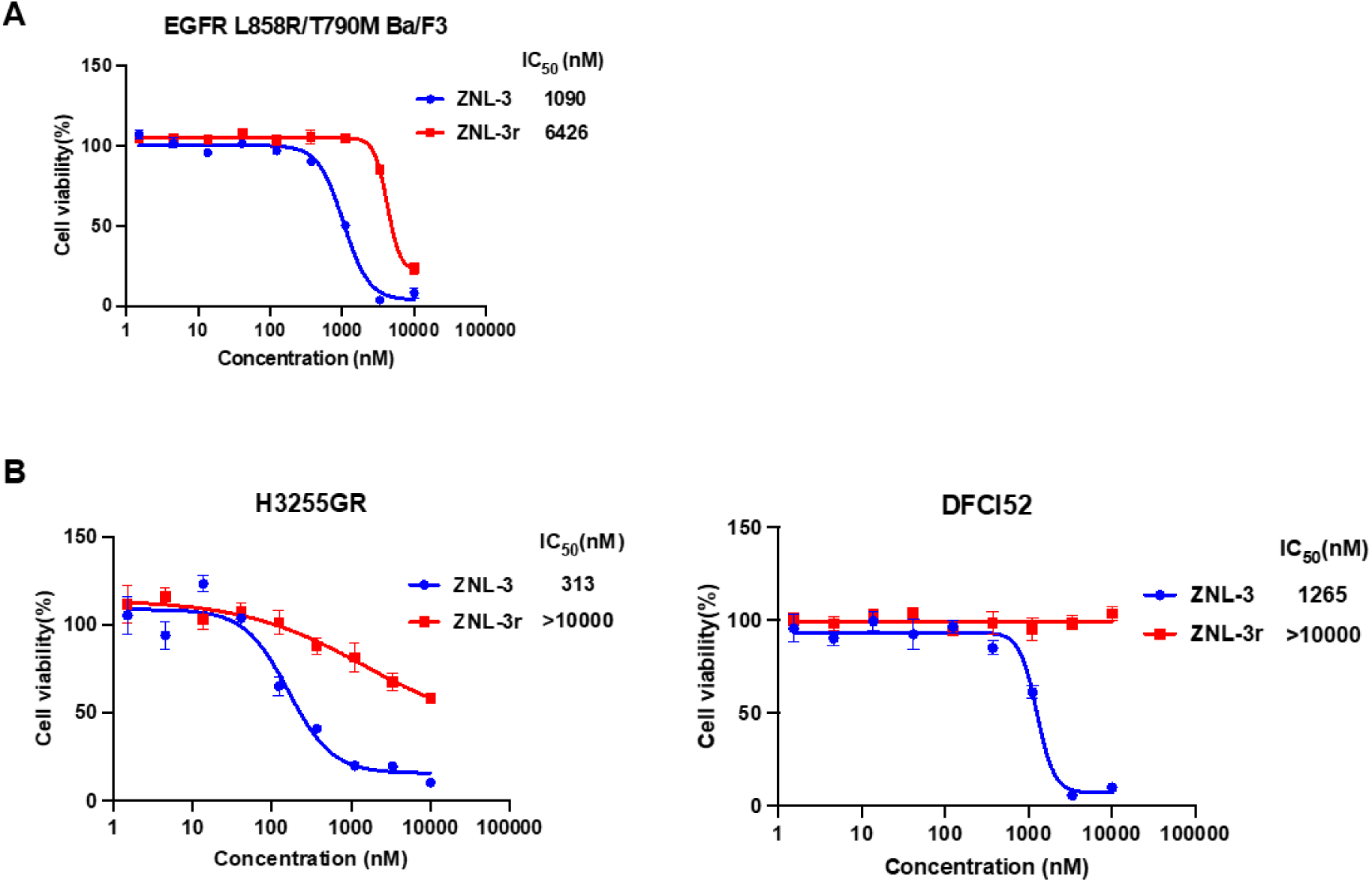
**A.** Dose-response curves for ZNL-3 and ZNL-3r in EGFR L858R/T790M Ba/F3 after 72 h treatment. Cell viability was assessed with CellTiter-Glo. Data are presented as mean±SEM. Representative data from three independent experiments are shown. **B**. Dose-response curves for ZNL-3 and ZNL-3r in H3255GR and DFCI52 after 72 h treatment. Cell viability was assessed with CellTiter-Glo. Data are presented as mean±SEM. Representative data from three independent experiments are shown.

**Figure S7.**
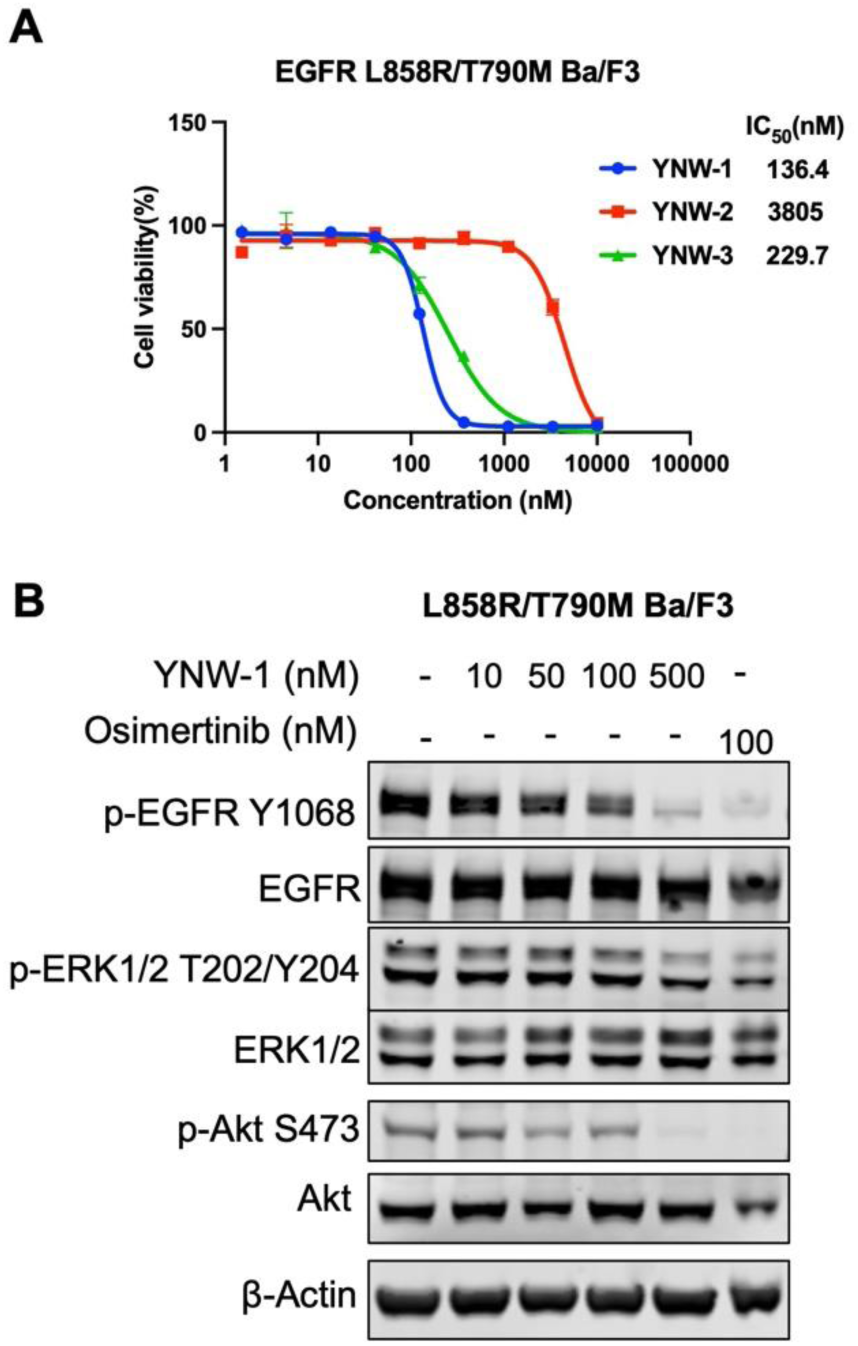
**A.** Dose-response curves for YNW-1, YNW-2, YNW-3 in EGFR L858R/T790M Ba/F3 and EGFR L858R/T790M/C797S Ba/F3 after 72 h treatment. Cell viability was assessed with CellTiter-Glo. Data are presented as mean±SEM. Representative data from three independent experiments are shown. **B**. Effects of YNW-1 on EGFR signaling pathway in EGFR L858R/T790M Ba/F3 after 6 h treatment. Representative data from three independent experiments are shown.

**Table S1.** Data collection and refinement statistics.

| **Data Collection** |  |
| --- | --- |
| PDB Accession | 00009PWI |
| Resolution range | 64.54 - 2.6 (2.86 - 2.6) |
| Space group | P 4<sub>3</sub> 2<sub>1</sub> 2 |
| Unit cell dimensions (Å) | 70.39 70.39 161.72 |
| Unit cell angles (°) | 90 90 90 |
| Total reflections | 84961 (21759) |
| Unique reflections | 13148 (3200) |
| Multiplicity | 6.5 (6.8) |
| Completeness (%) | 99.51 (99.44) |
| Mean I/sigma(I) | 9.03 (1.03) |
| Wilson B-factor | 71.24 |
| R<sub>merge</sub> | 0.0906 (0.8069) |
| R<sub>meas</sub> | 0.0989 (0.8739) |
| R<sub>pim</sub> | 0.0389 (0.3309) |
| CC<sub>1/2</sub> | 0.997 (0.882) |
| **Data Refinement** |  |
| Reflections used in refinement | 13108 (3183) |
| Reflections used for R<sub>free</sub> | 605 (141) |
| R<sub>work</sub> | 0.224 (0.334) |
| R<sub>free</sub> | 0.254 (0.358) |
| Number of non-hydrogen atoms | 2243 |
| macromolecules | 2202 |
| ligand | 38 |
| solvent | 3 |
| Protein residues | 274 |
| RMS (bonds) | 0.177 |
| RMS (angles) | 3.18 |
| Ramachandran favored (%) | 94.1 |
| Ramachandran allowed (%) | 5.5 |
| Ramachandran outliers (%) | 0.4 |
| Rotamer outliers (%) | 4.5 |
| Average B-factor | 84.4 |
| macromolecules | 84.4 |
| ligands | 84.1 |

*Statistics for the highest-resolution shell are shown in parentheses.*

**Table S2.** Comparison of *in vitro* metabolic stability for compound ZNL-2S and ZNL-3

| Cmpd ID | Whole blood stability- mouse:T1/2(min) | Whole blood stability- human:T1/2(min) | Hepatocytes Stability- mouse T1/2(min) | Hepatocytes Stability- mouse Clint (hep) (μL/min/10e6) | Hepatocytes Stability- mouse Clint (liver) (μL/min/kg) | Hepatocytes Stability- human T1/2(min) | Hepatocytes Stability- human CLint (hep) (uL/min/10e6) | Hepatocytes Stability- human CLint (live) (uL/min/10e6) |
| --- | --- | --- | --- | --- | --- | --- | --- | --- |
| ZNL-2S | 12.4 | 105 | 3.8 | 363 | 4310 | 24.3 | 56.9 | 158 |
| ZNL-3 | 205 | 214 | 11.2 | 124 | 1470 | 31.2 | 44.5 | 124 |

**Table S3.** Pharmacokinetics in mouse for ZNL-3

| Pharmacokinetics of ZNL-3 (ng/mL) |  |  |  |  |  |  |  |  |  |
| --- | --- | --- | --- | --- | --- | --- | --- | --- | --- |
| IV bolus1 |  |  |  |  | PO2 |  |  |  |  |
| PK Parameters | M01 | M02 | M03 | Mean | PK Parameters | M04 | M05 | M06 | Mean |
| Rs<sub>q</sub>_adj | 0.996 | 0.996 | 0.988 | — | Rs<sub>q</sub>_adj | 0.998 | 0.997 | 0.979 | — |
| No. points used for T<sub>1/2</sub> | 3.00 | 3.00 | 3.00 | 3.00 | No. points used for T<sub>1/2</sub> | 3.00 | 3.00 | 3.00 | 3.00 |
| C<sub>0</sub> (ng/mL) | 527 | 519 | 779 | 608 | C<sub>max</sub> (ng/mL) | 119 | 738 | 186 | 348 |
| T<sub>1/2</sub> (h) | 1.36 | 1.10 | 0.798 | 1.09 | T<sub>max</sub> (h) | 0.500 | 1.00 | 0.250 | 0.583 |
| V<sub>d</sub><sub>ss</sub> (L/kg) | 3.35 | 4.50 | 2.60 | 3.48 | T<sub>1/2</sub> (h) | 1.42 | 1.49 | 2.05 | 1.65 |
| Cl (mL/min/kg) | 54.7 | 69.8 | 56.7 | 60.4 | T<sub>last</sub> (h) | 8.00 | 8.00 | 8.00 | 8.00 |
| T<sub>last</sub> (h) | 8.00 | 4.00 | 4.00 | ND | AUC<sub>0-last</sub> (ng.h/mL) | 360 | 1768 | 381 | 836 |
| AUC<sub>0-last</sub> (ng.h/mL) | 302 | 227 | 288 | 272 | AUC<sub>0-inf</sub> (ng.h/mL) | 370 | 1813 | 407 | 863 |
| AUC<sub>0-inf</sub> (ng.h/mL) | 305 | 239 | 294 | 279 | MRT<sub>0-last</sub> (h) | 2.42 | 2.02 | 2.21 | 2.22 |
| MRT<sub>0-last</sub> (h) | 0.954 | 0.842 | 0.677 | 0.824 | MRT<sub>0-inf</sub> (h) | 2.62 | 2.22 | 2.77 | 2.54 |
| MRT<sub>0-inf</sub> (h) | 1.02 | 1.08 | 0.764 | 0.955 | AUC<sub>Extra</sub> (%) | 2.70 | 2.49 | 6.37 | 3.85 |
| AUC<sub>Extra</sub> (%) | 0.727 | 4.93 | 1.95 | 2.54 | AUMC<sub>Extra</sub> (%) | 10.4 | 11.4 | 25.2 | 15.7 |
| AUMC<sub>Extra</sub> (%) | 7.10 | 25.6 | 13.2 | 15.3 | Bioavailability (%)<sup>a</sup> | - | - | - | 103 |

Note: Mouse #5 was treated as an outlier and bioavailability was re-calculated

**Table S4.**
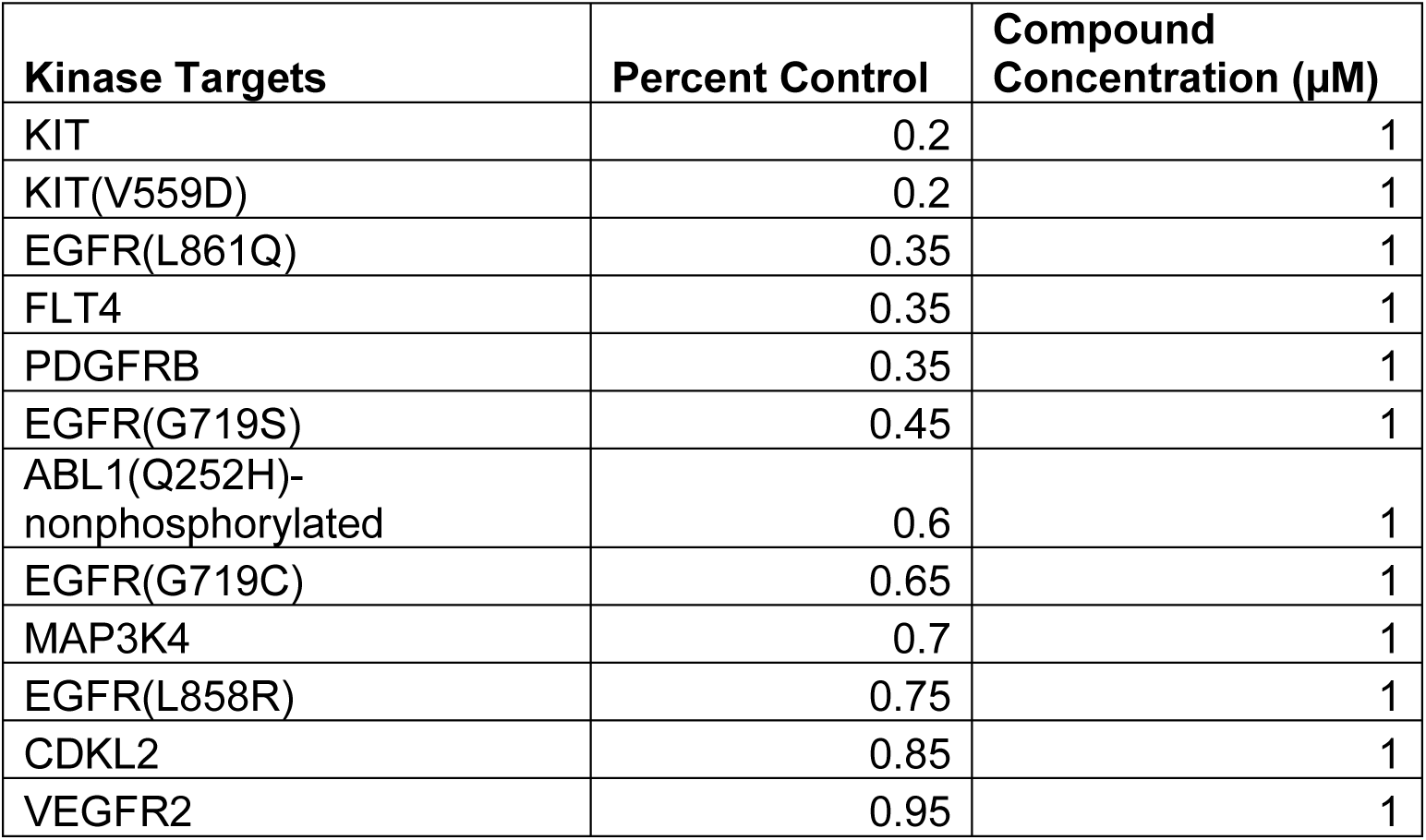
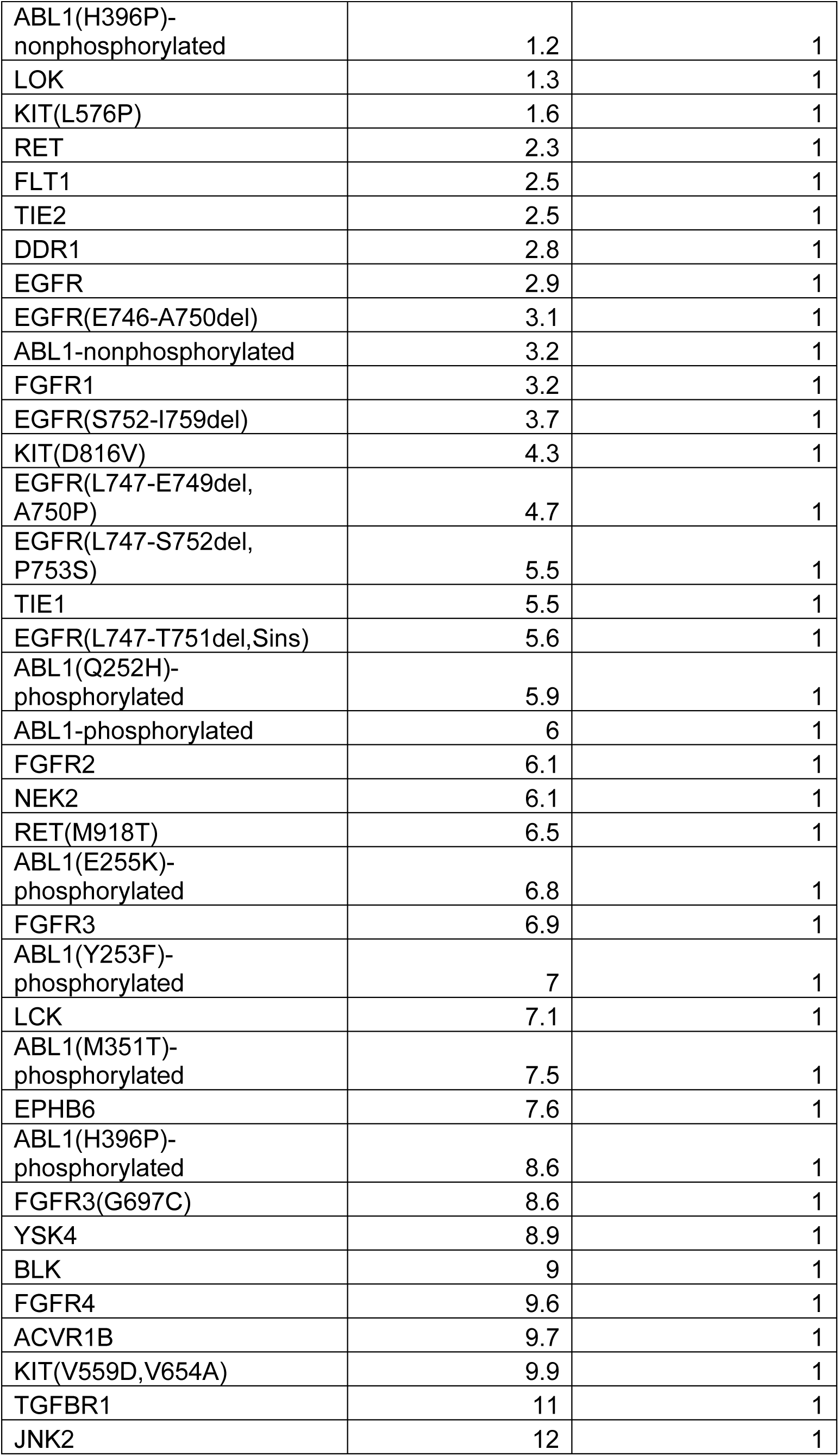
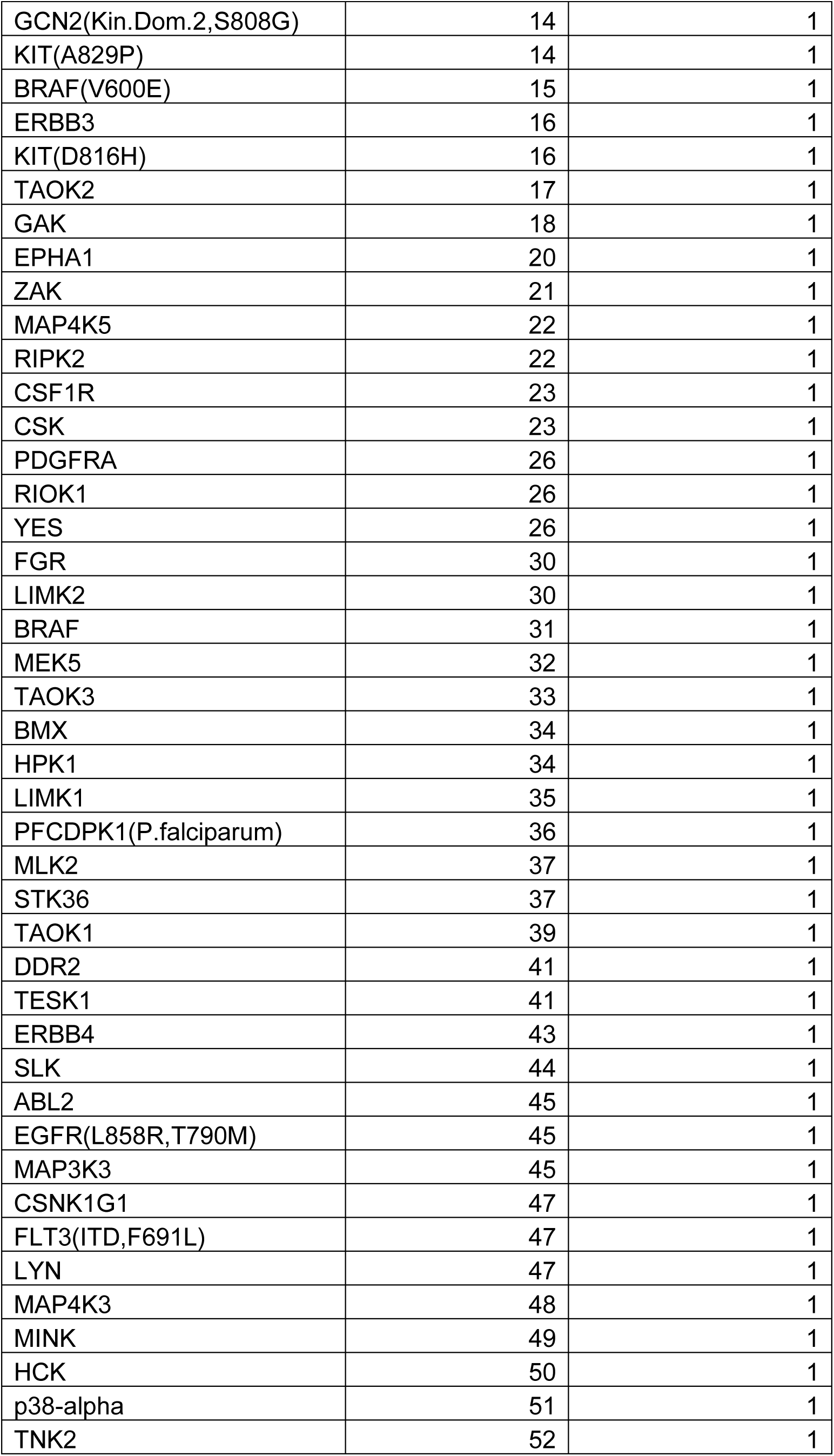
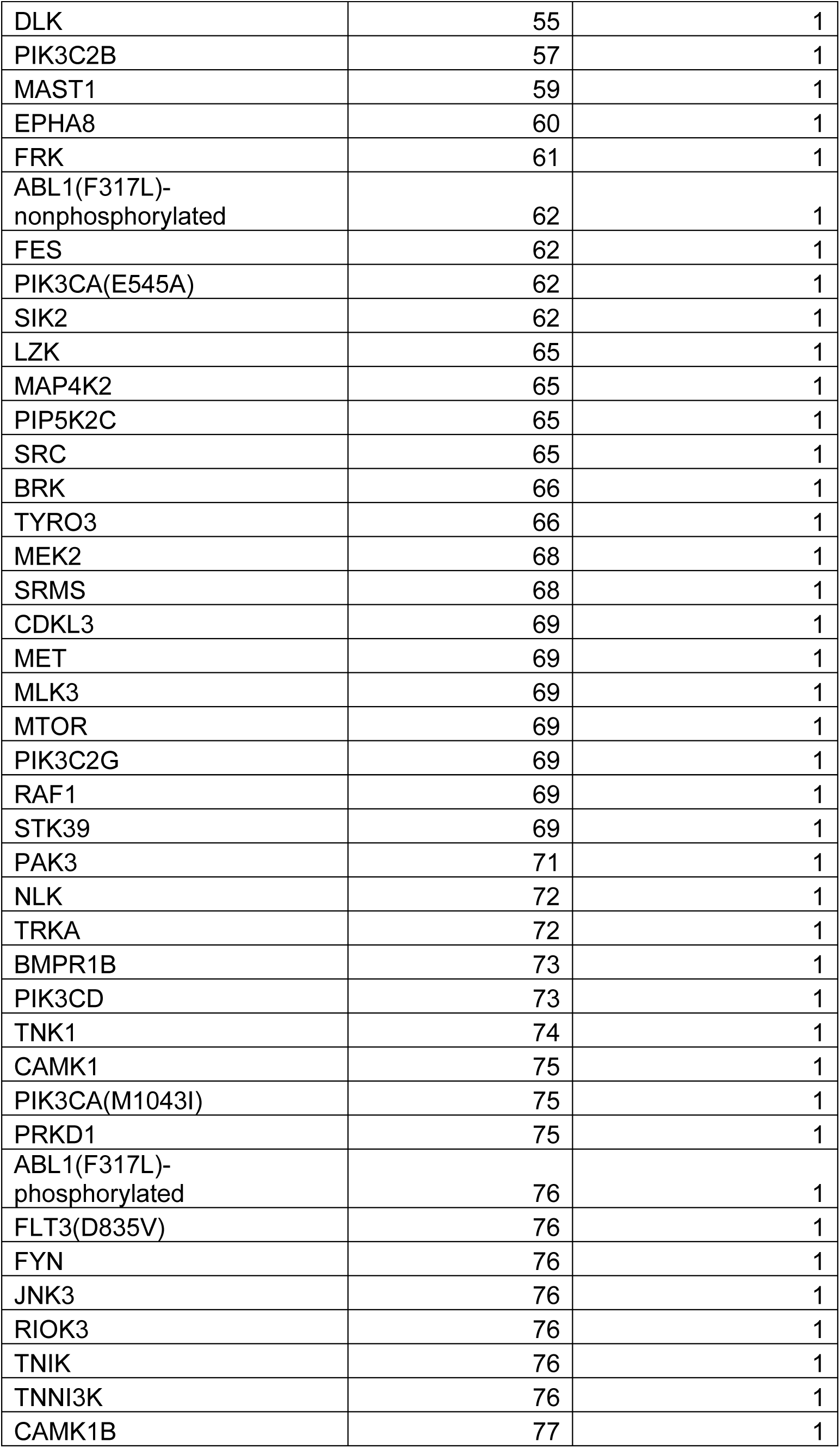
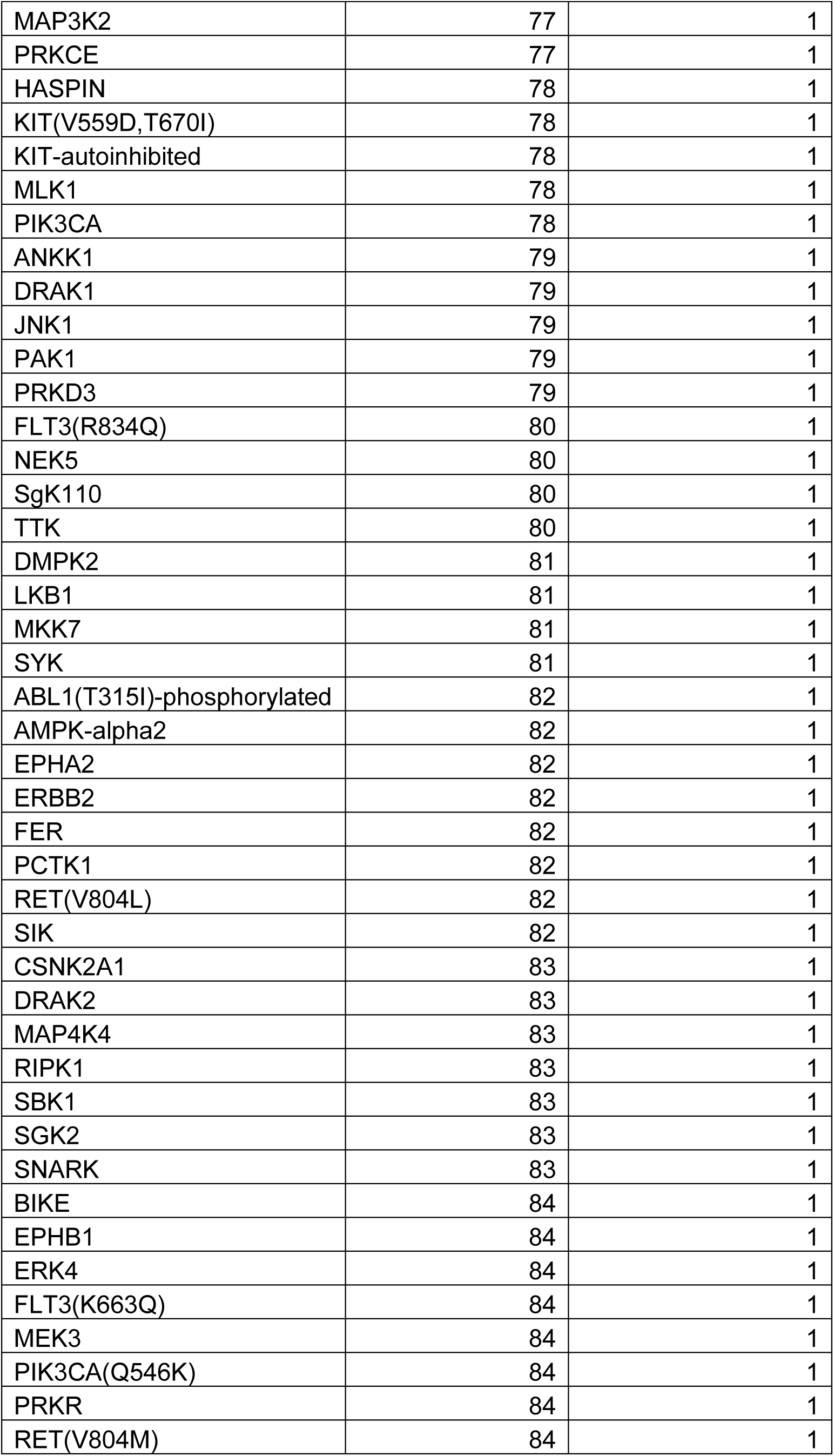
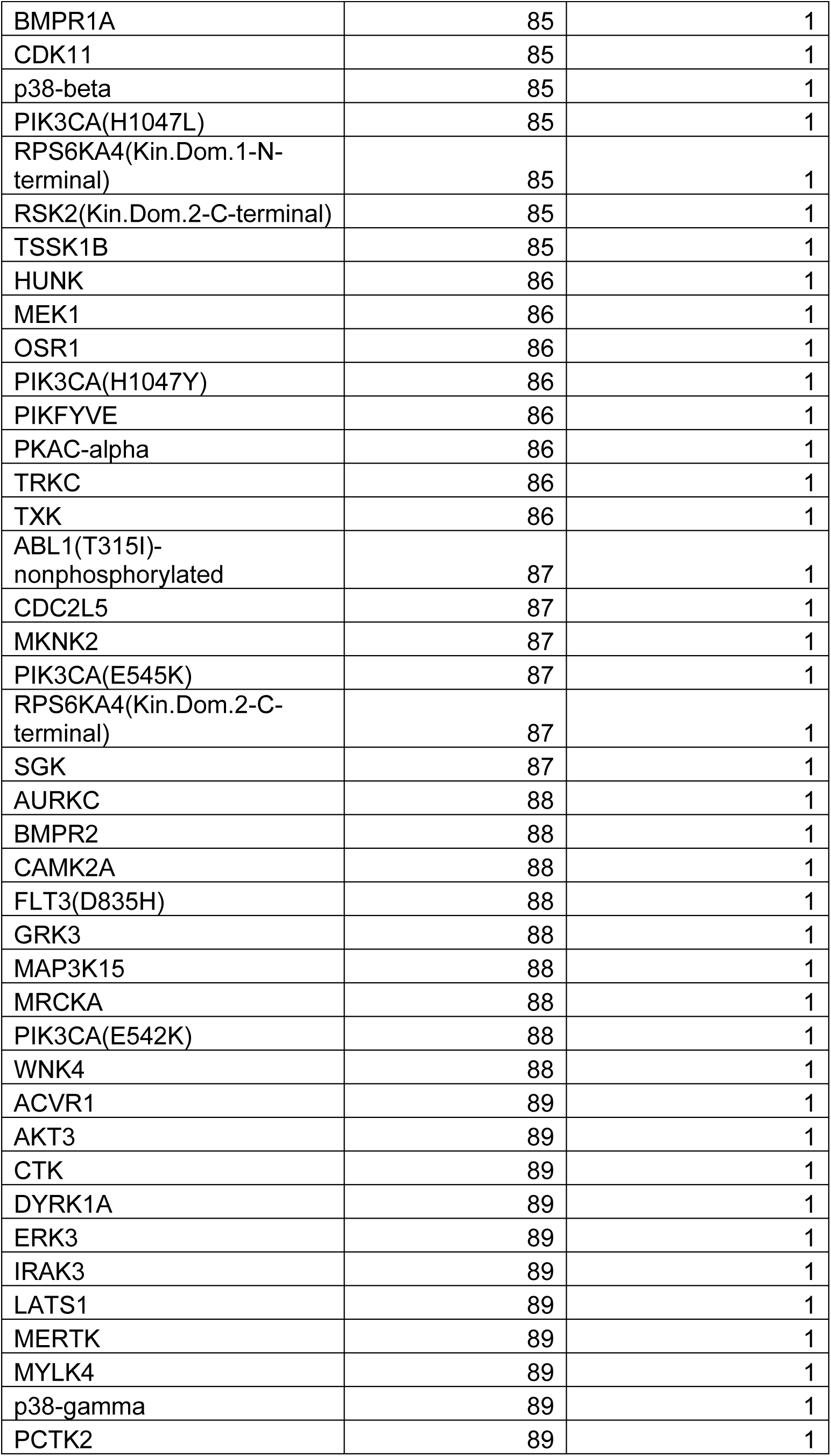
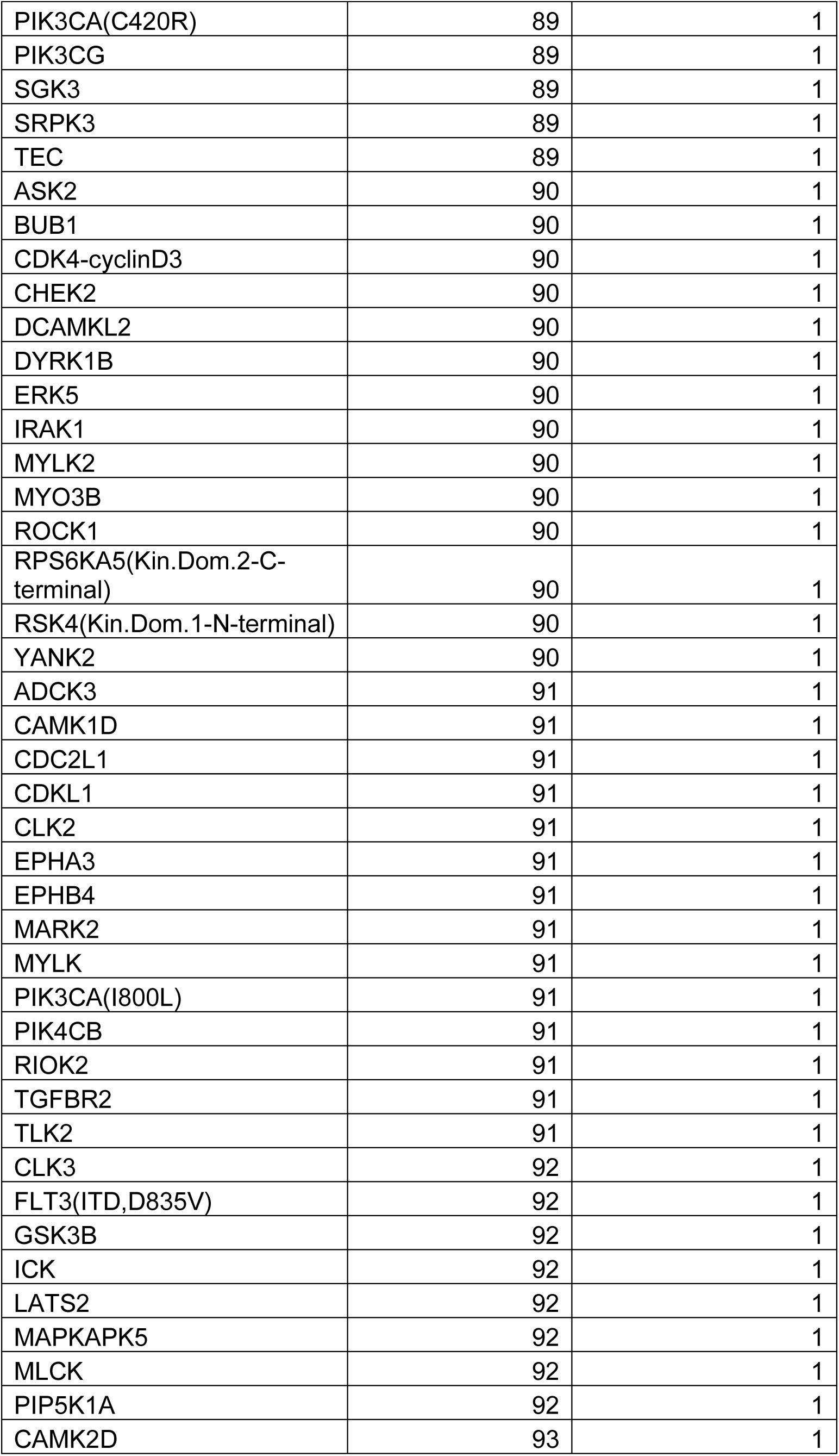
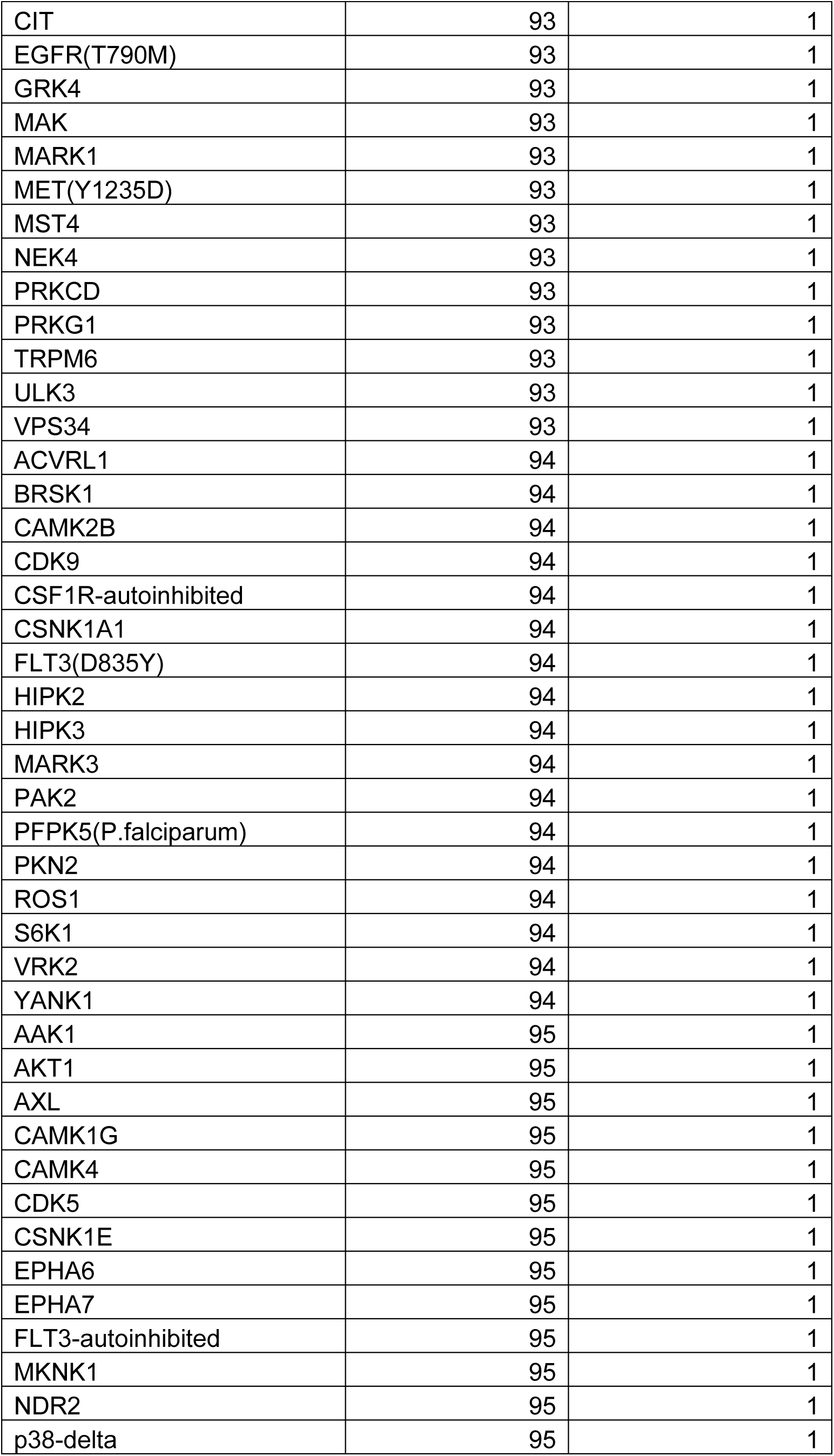
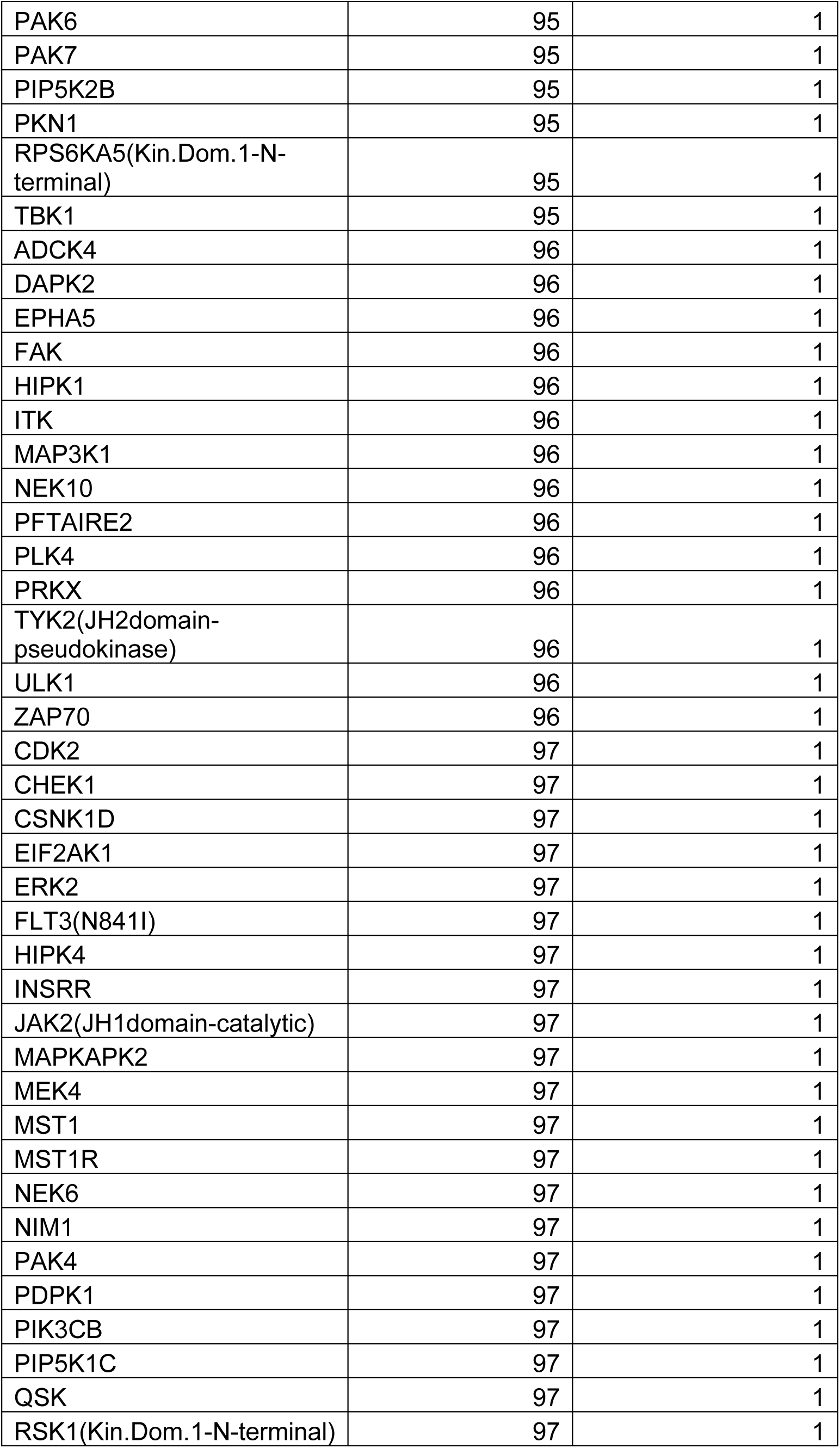
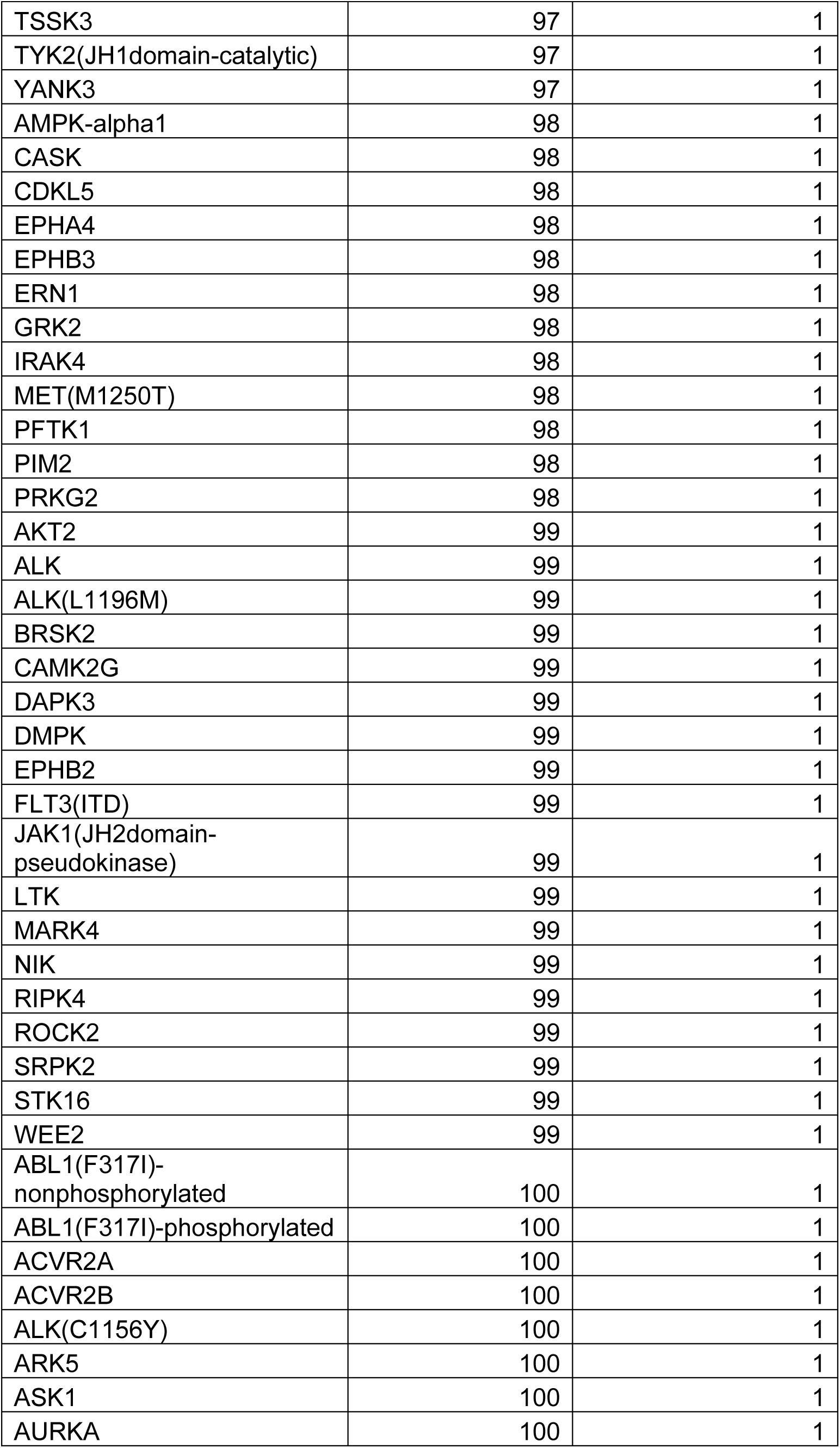
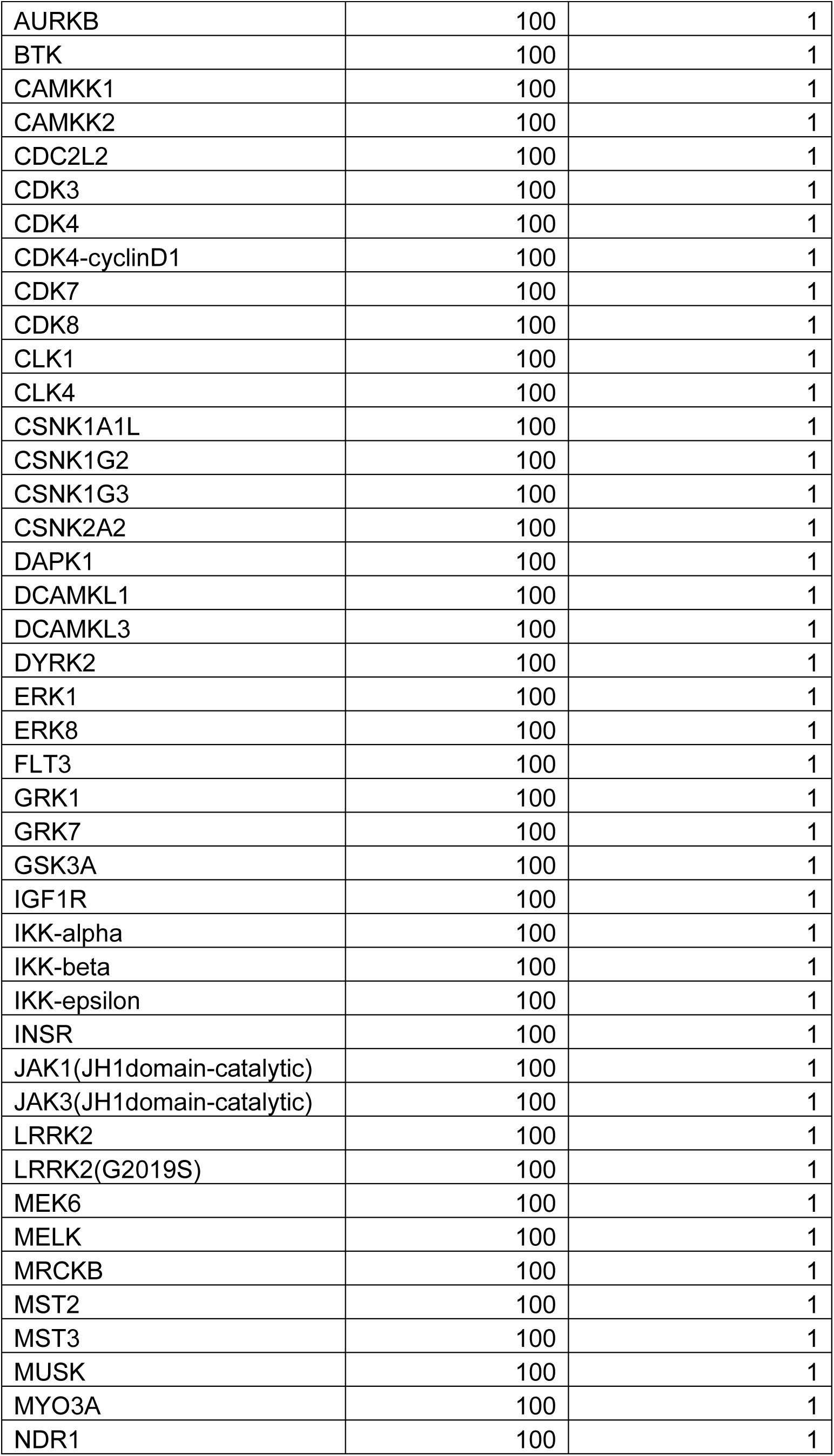
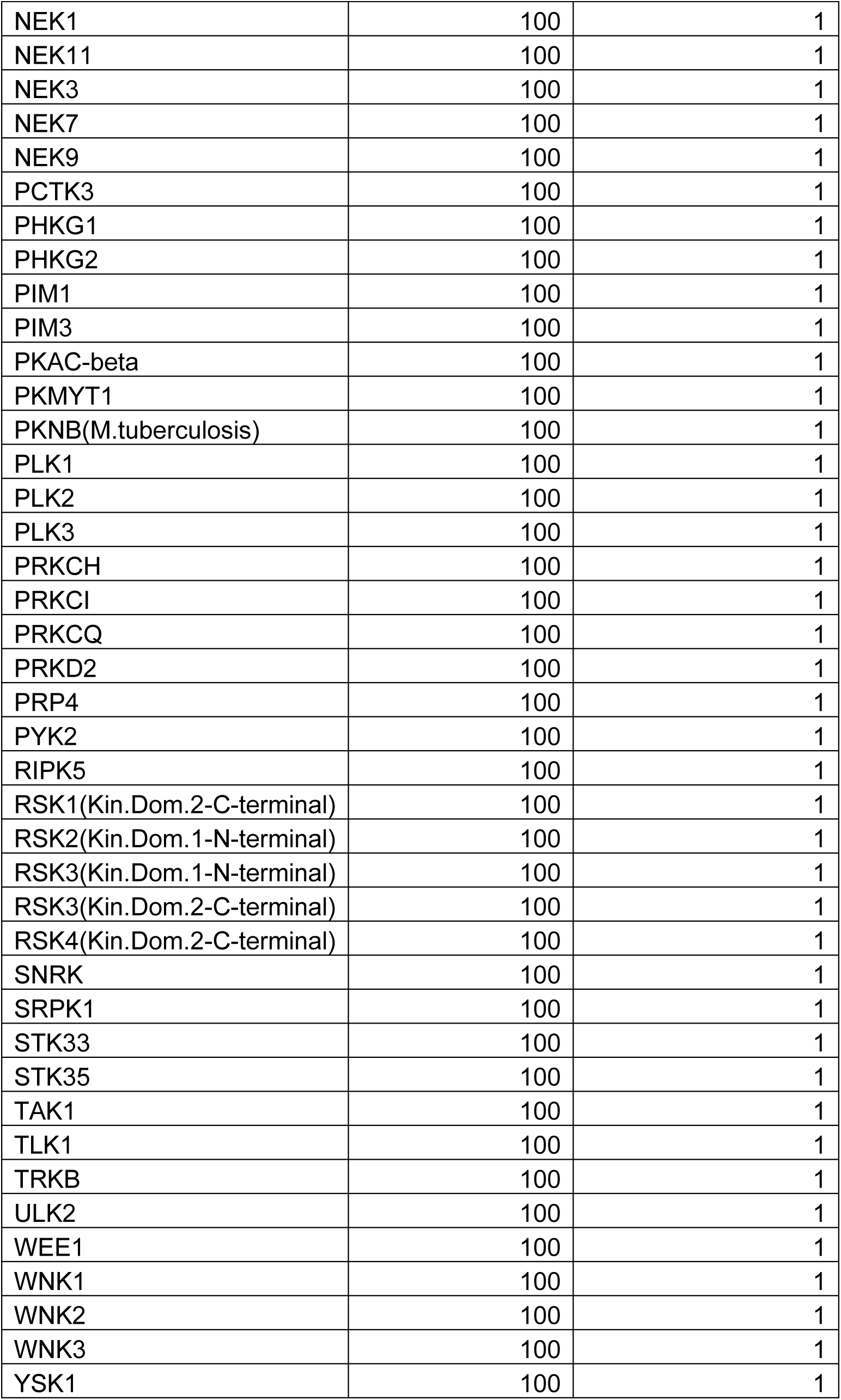
ZNL-8. DiscoverX KINOME*scan* at 1 µM

**Table S5.**
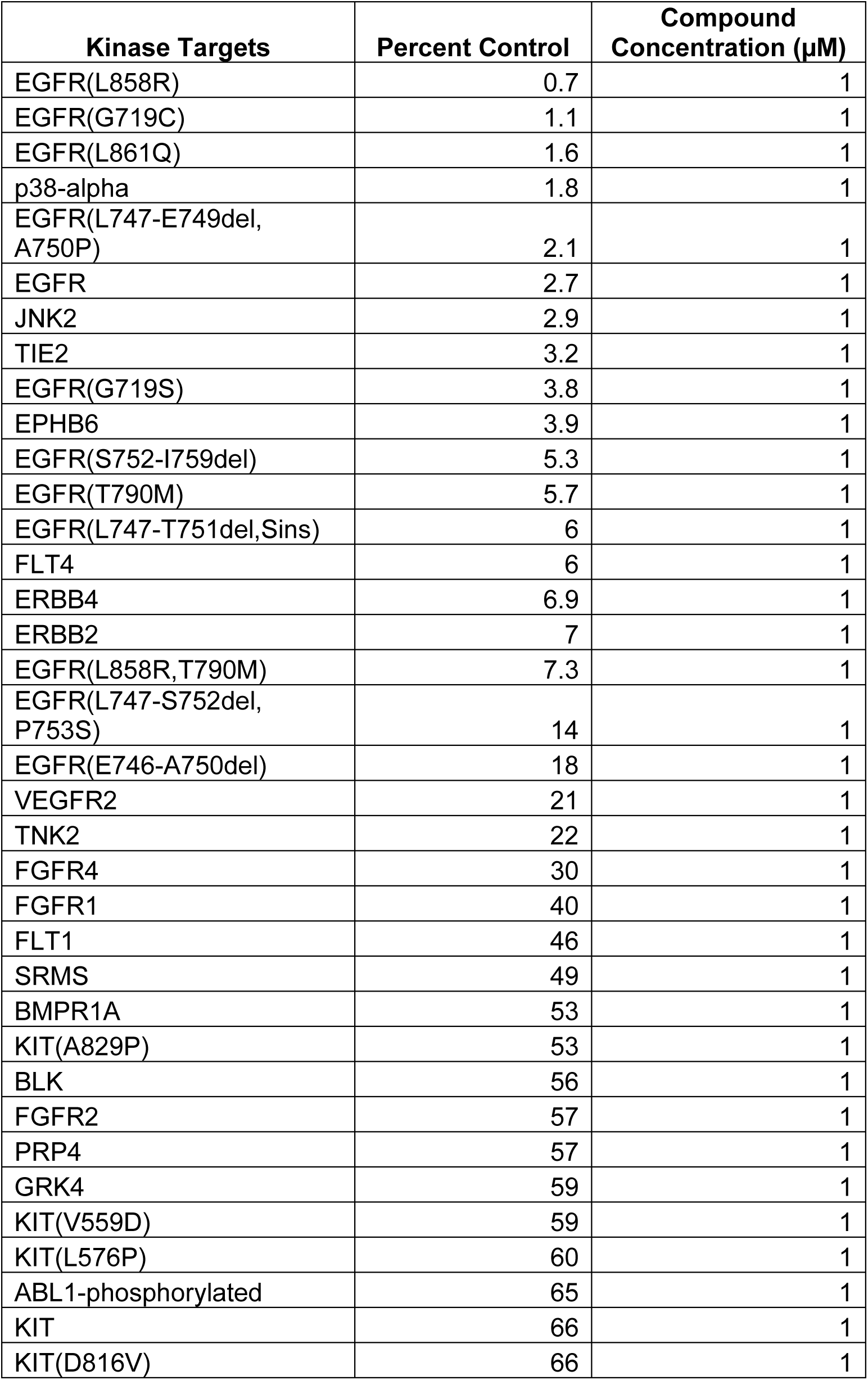
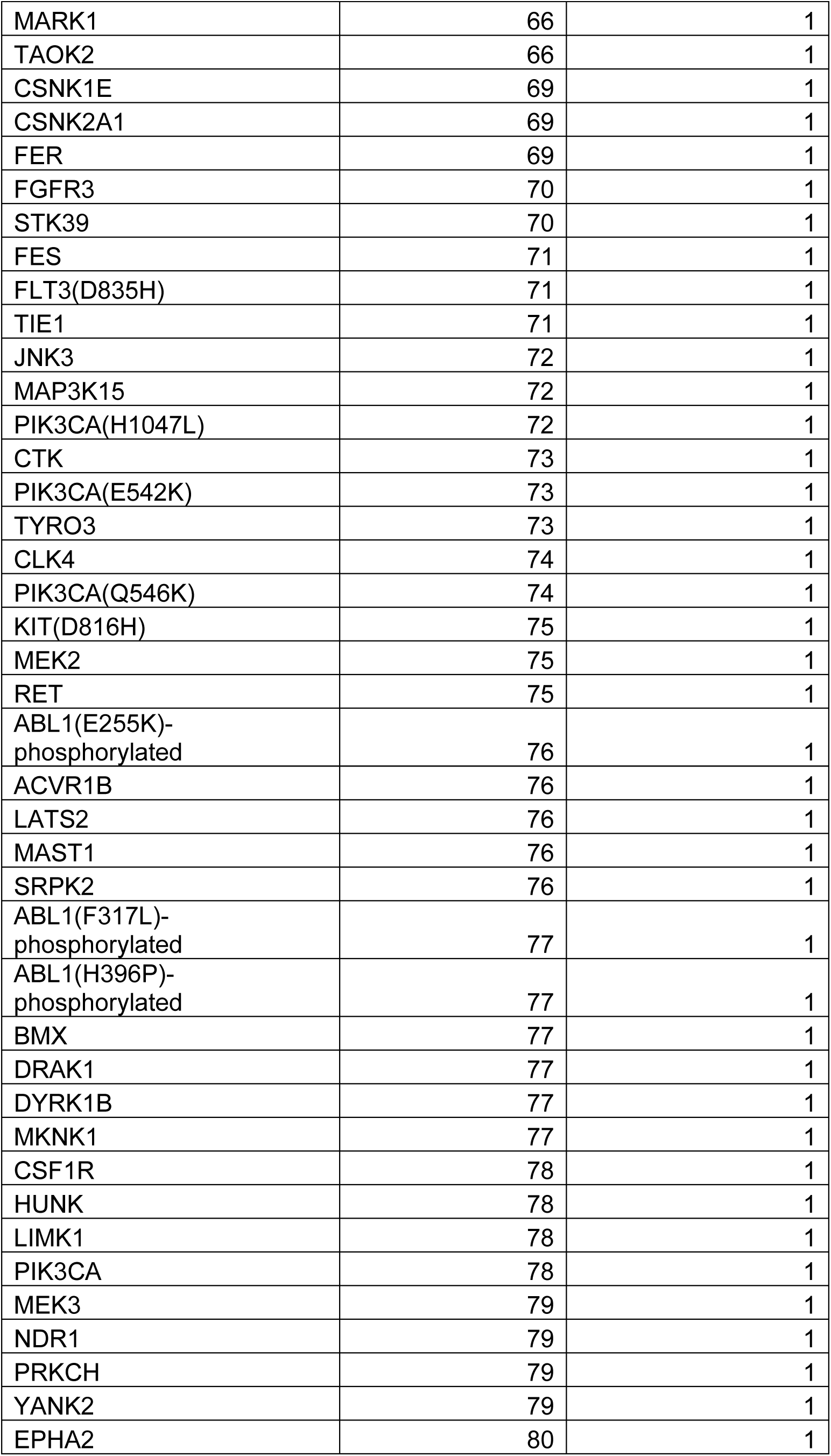
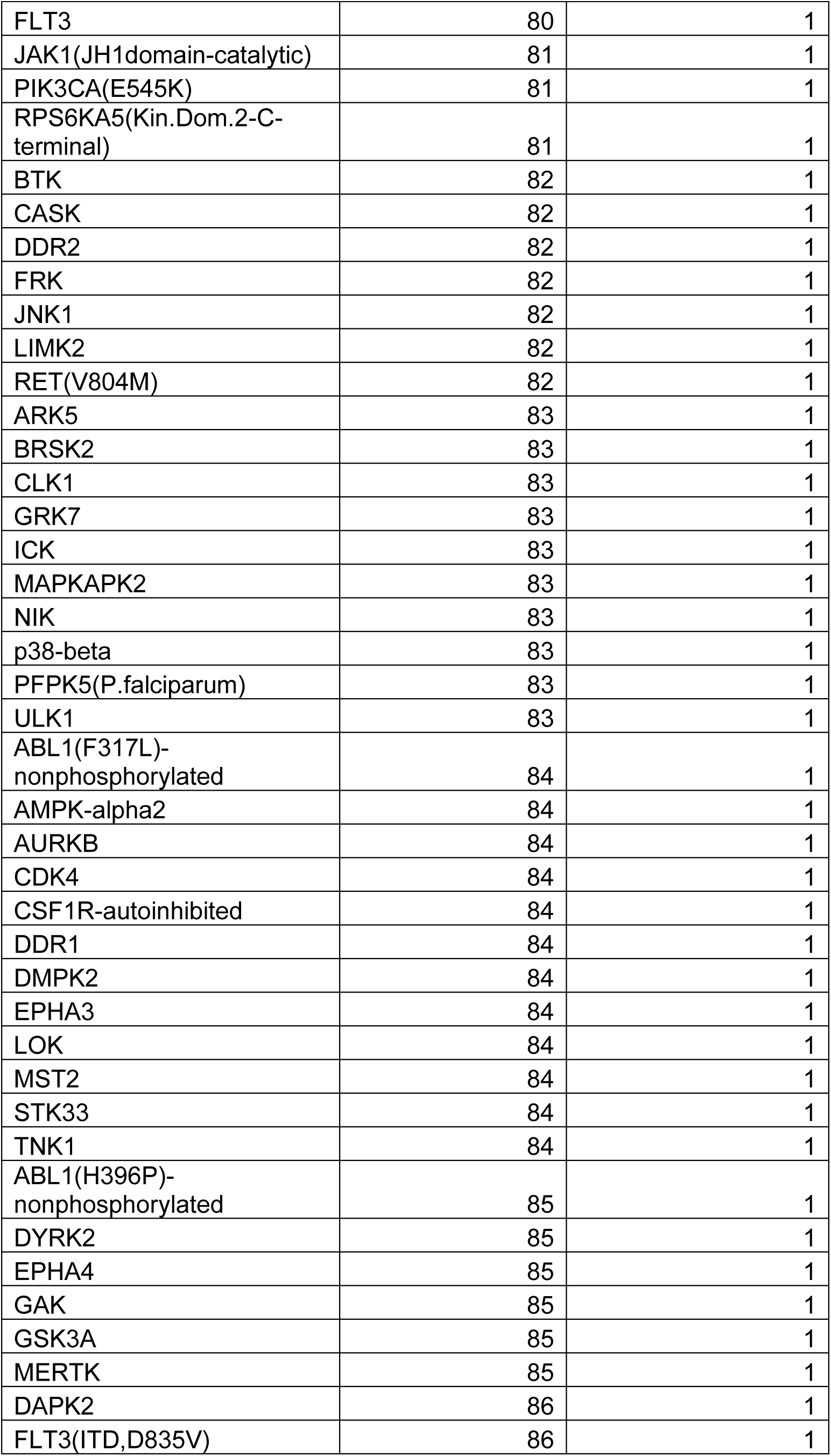
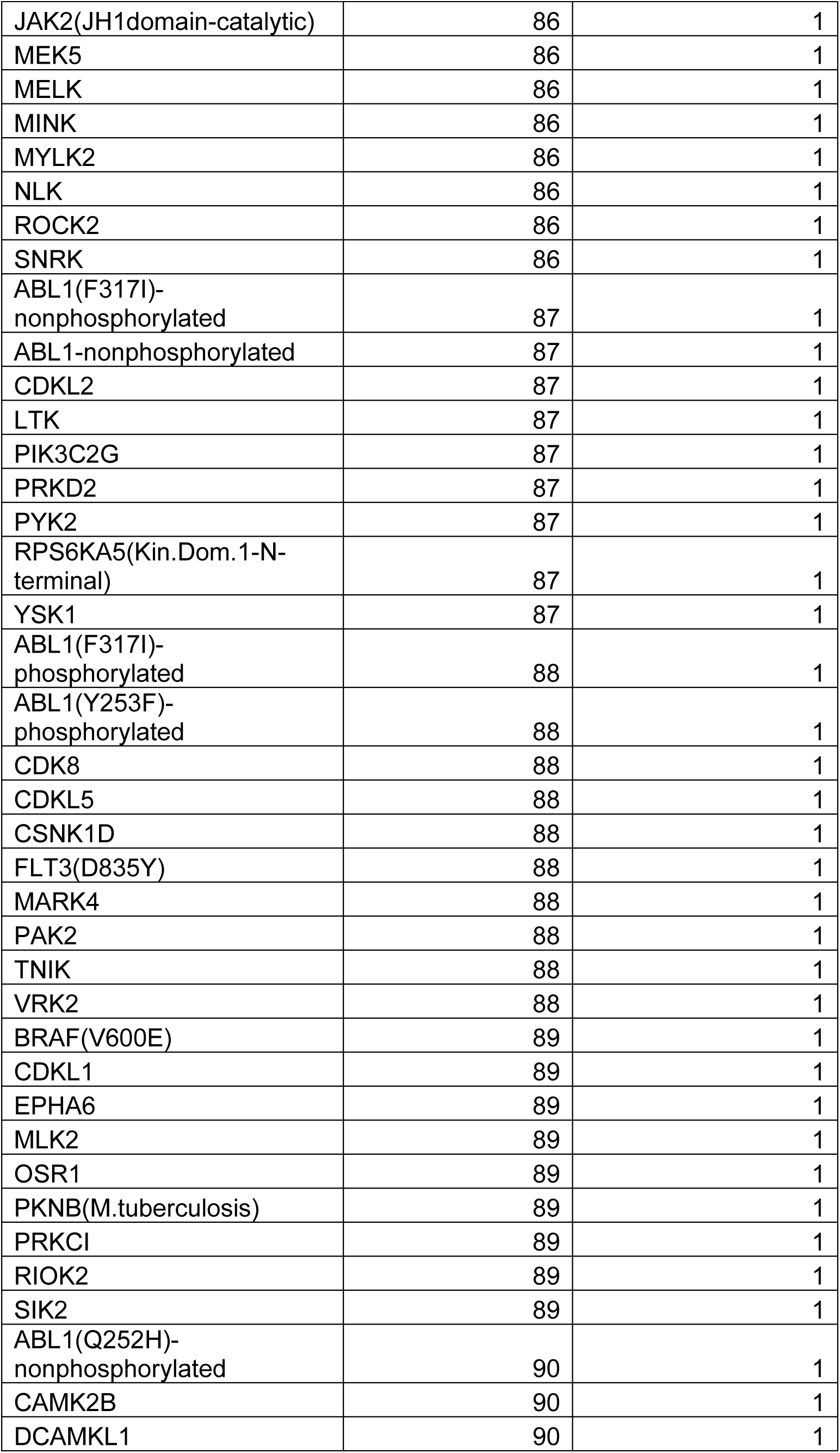
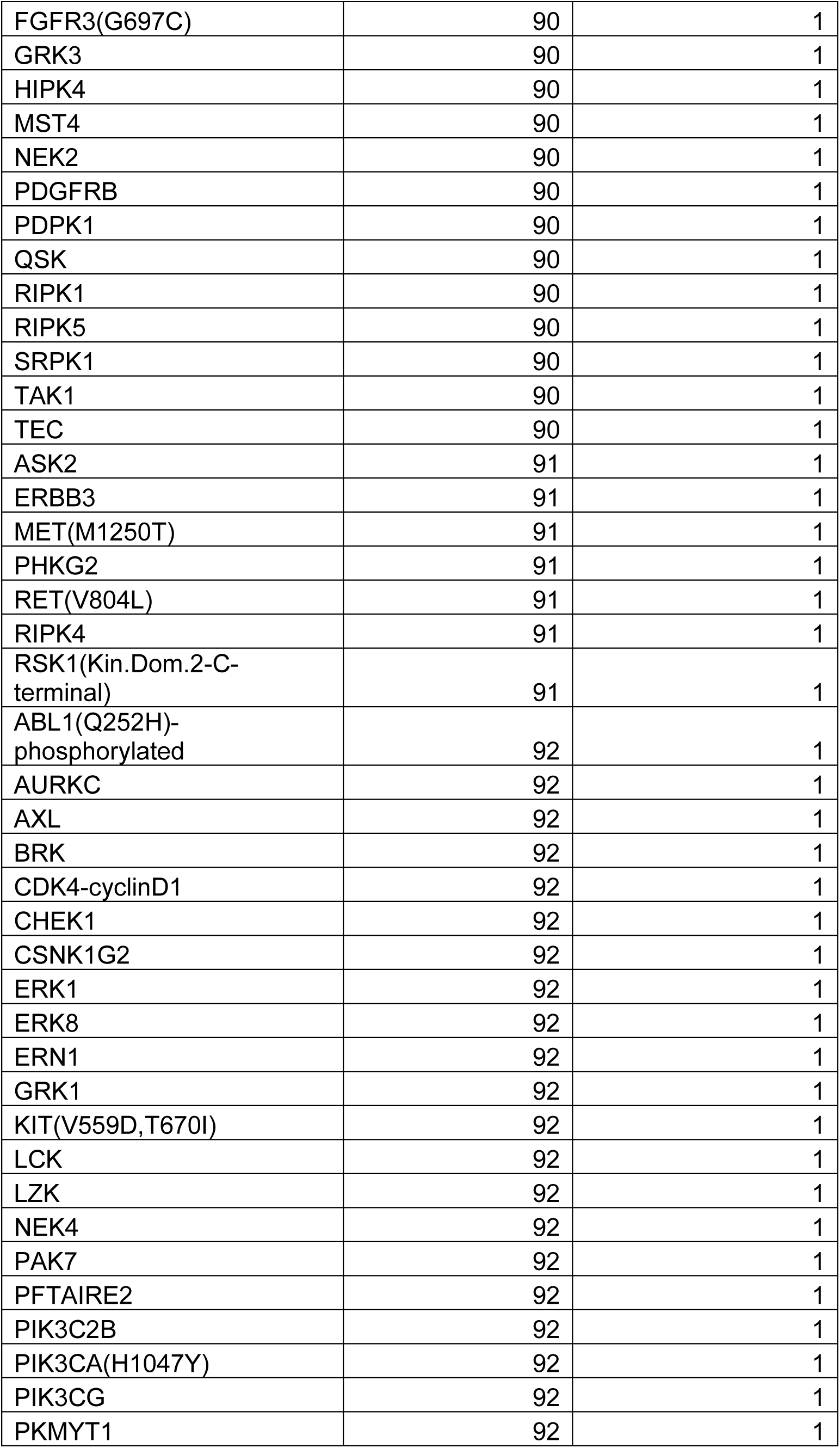
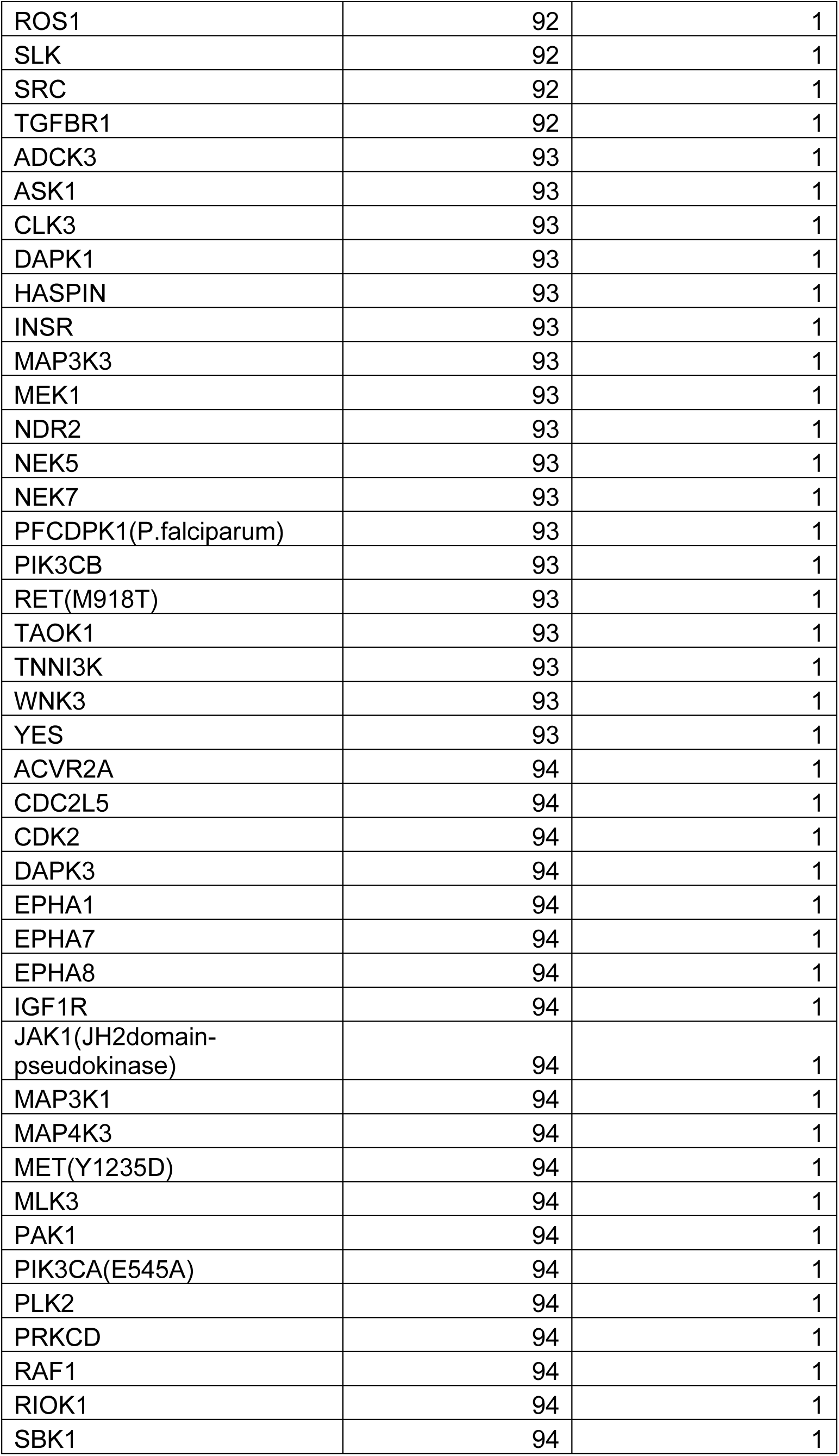
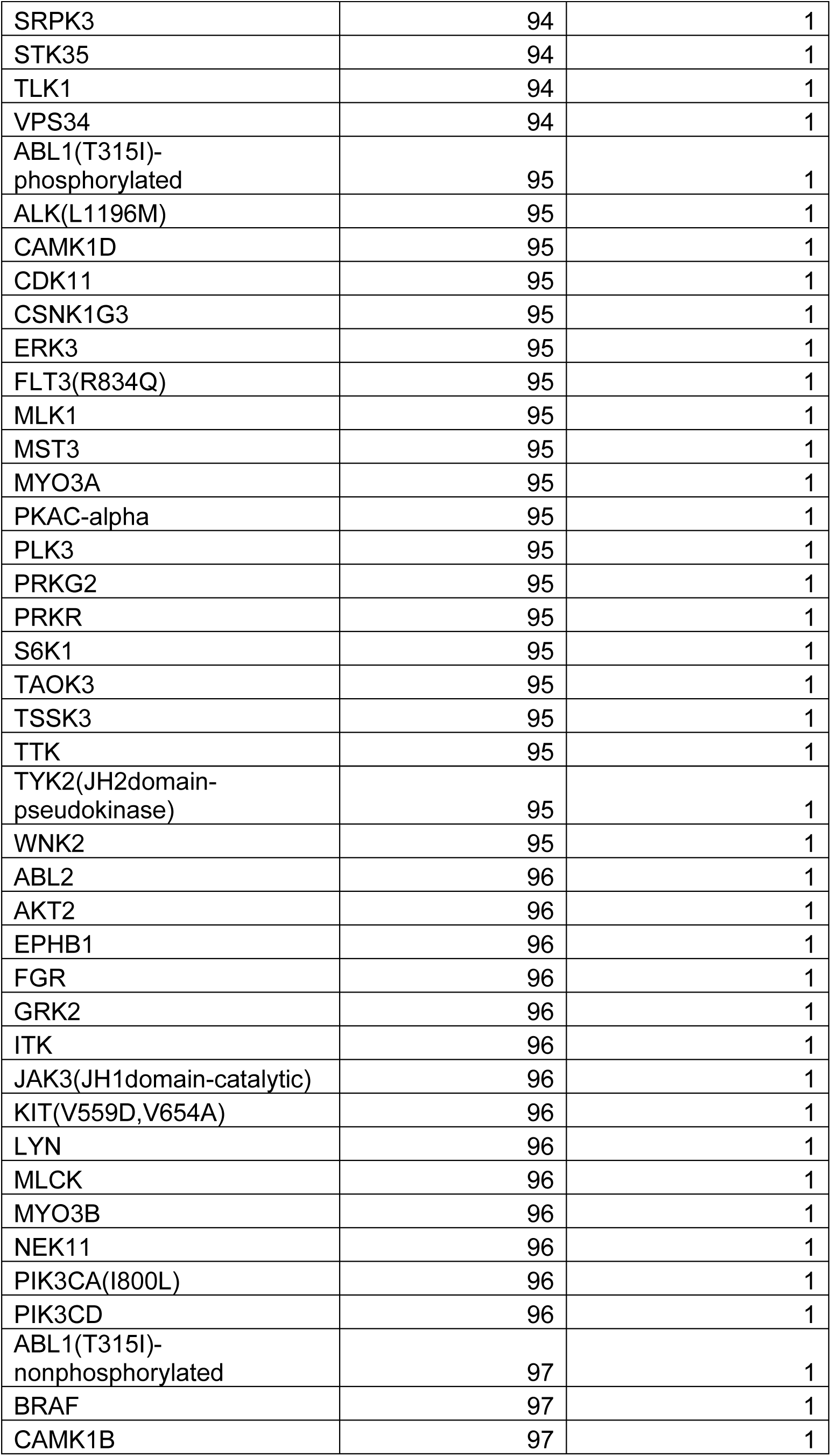
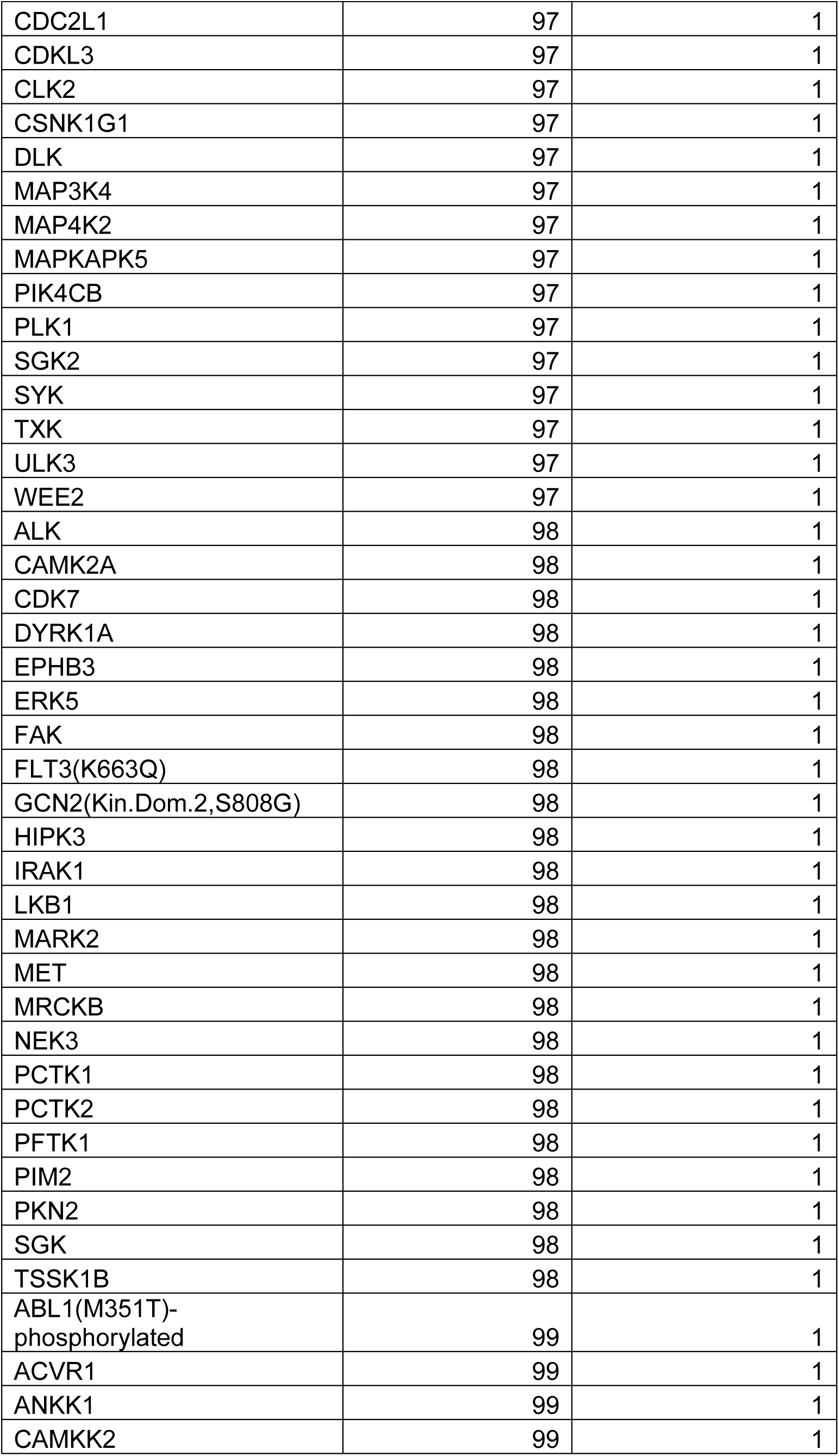
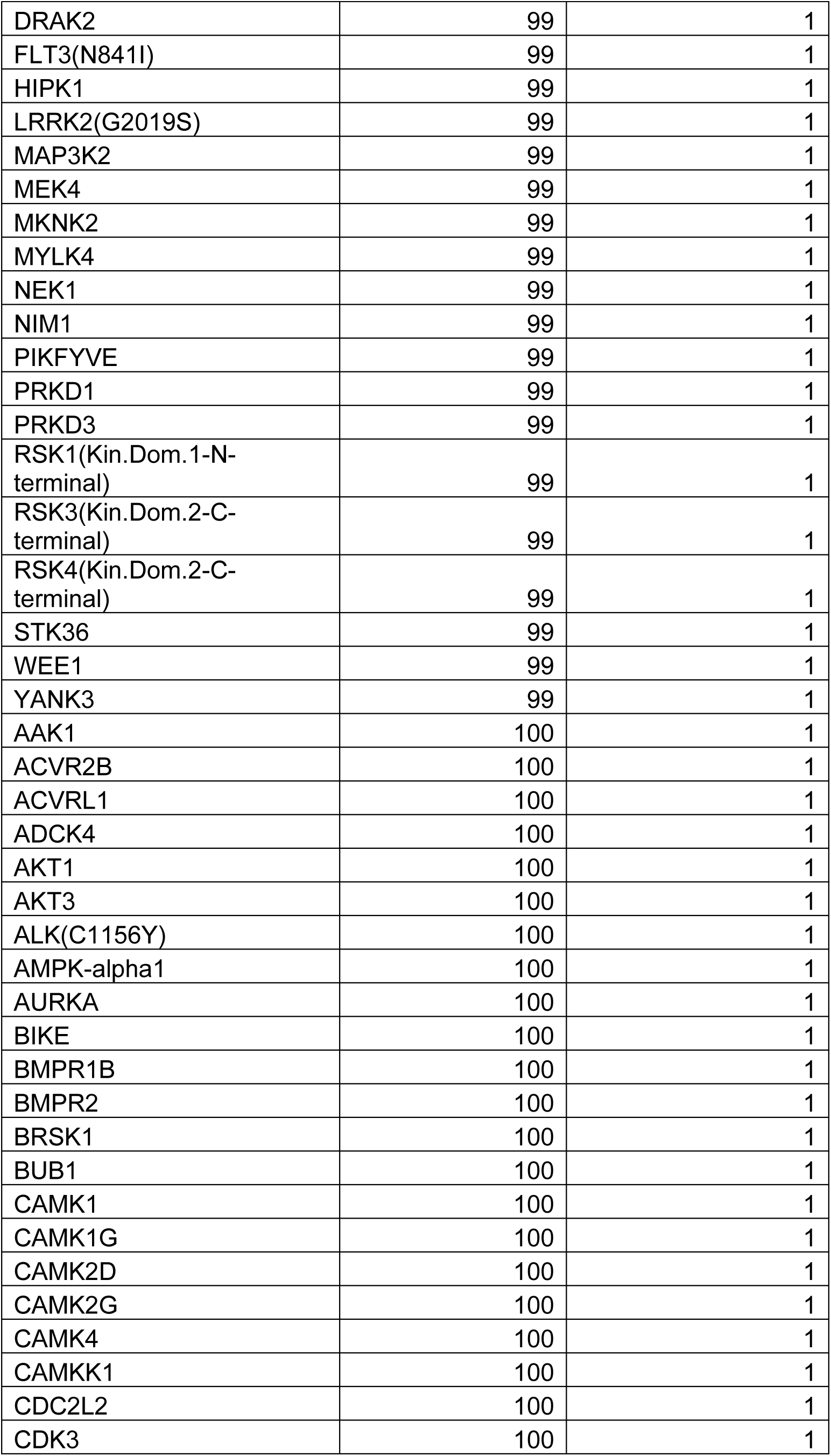
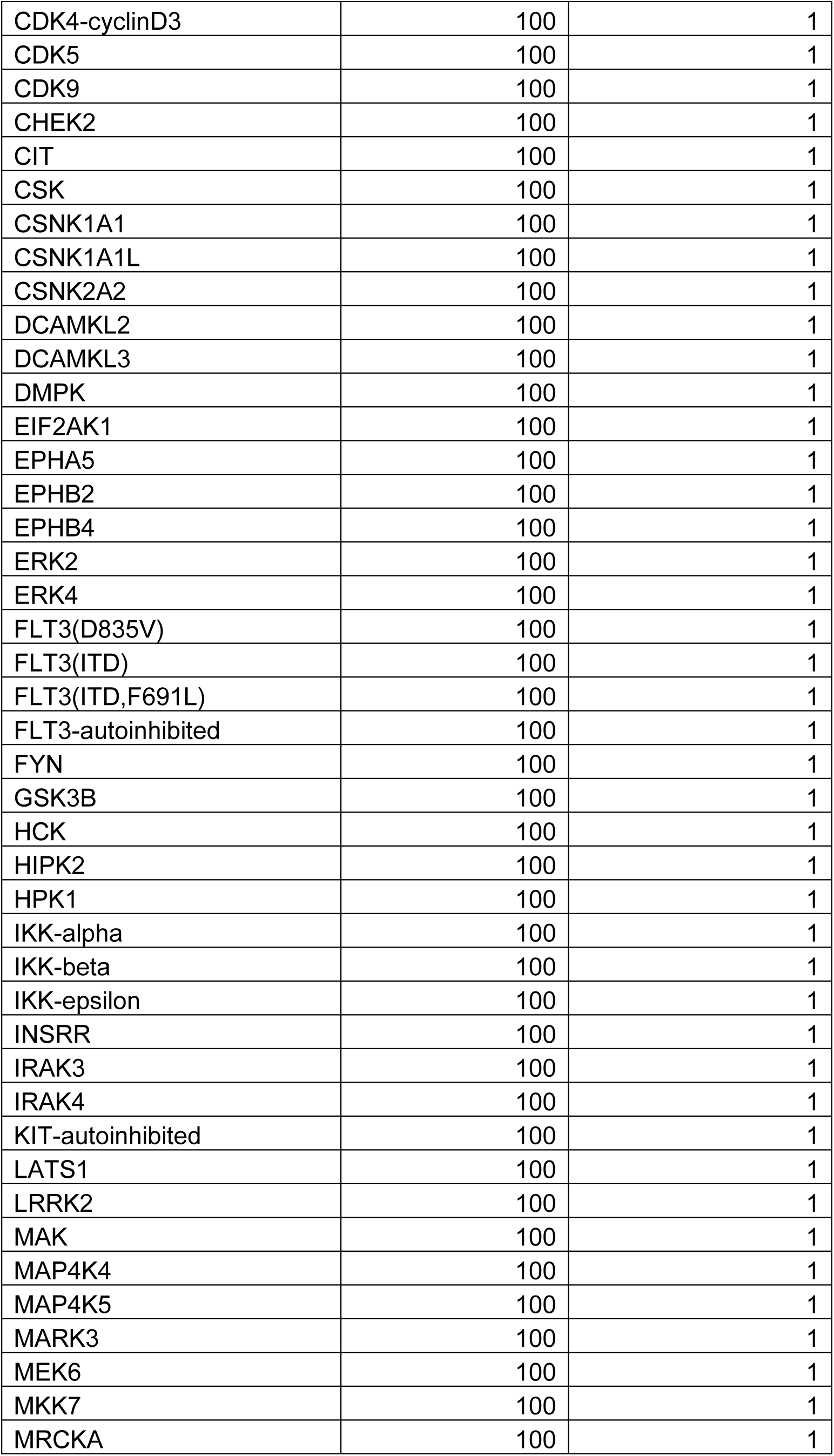
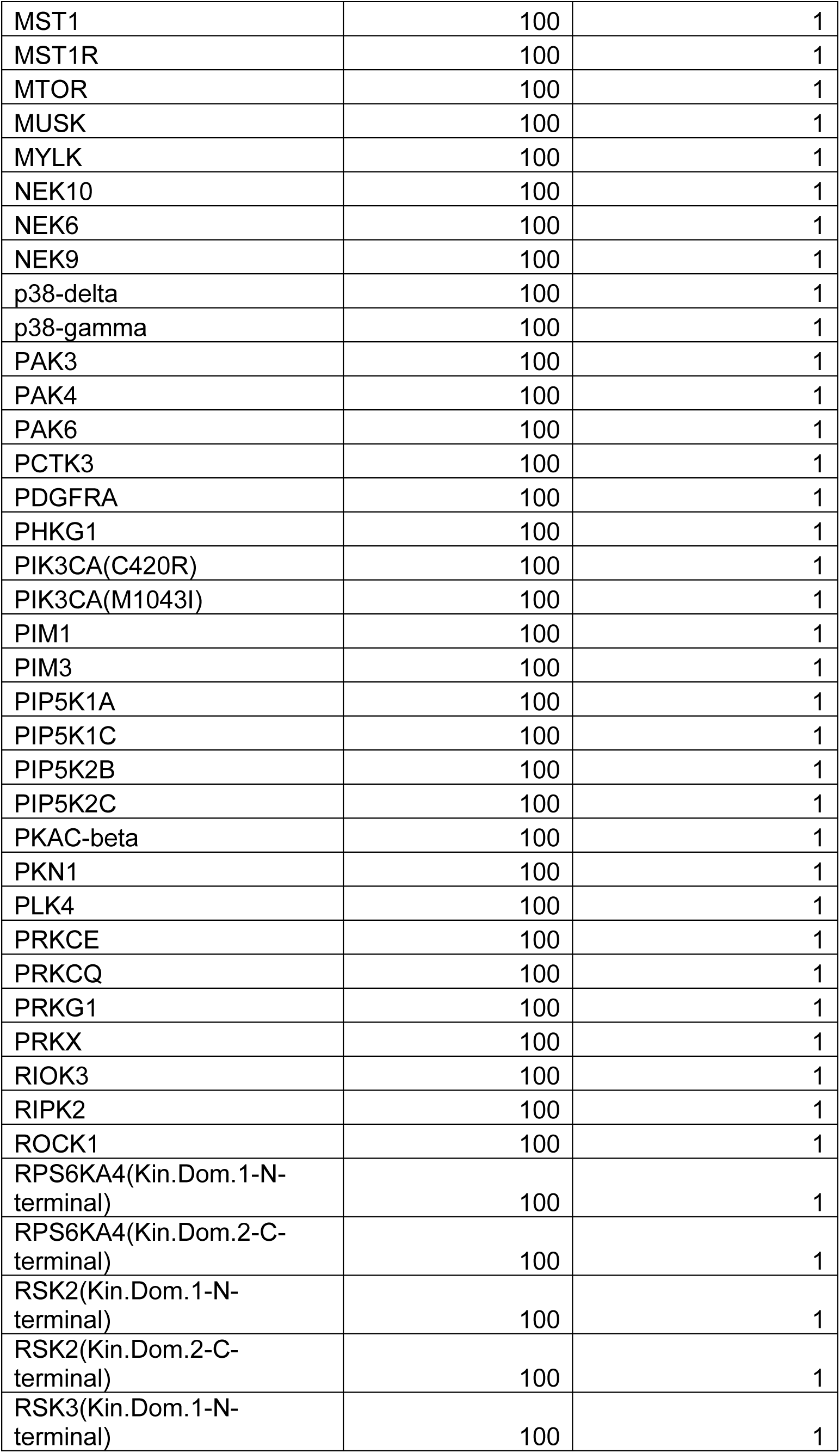
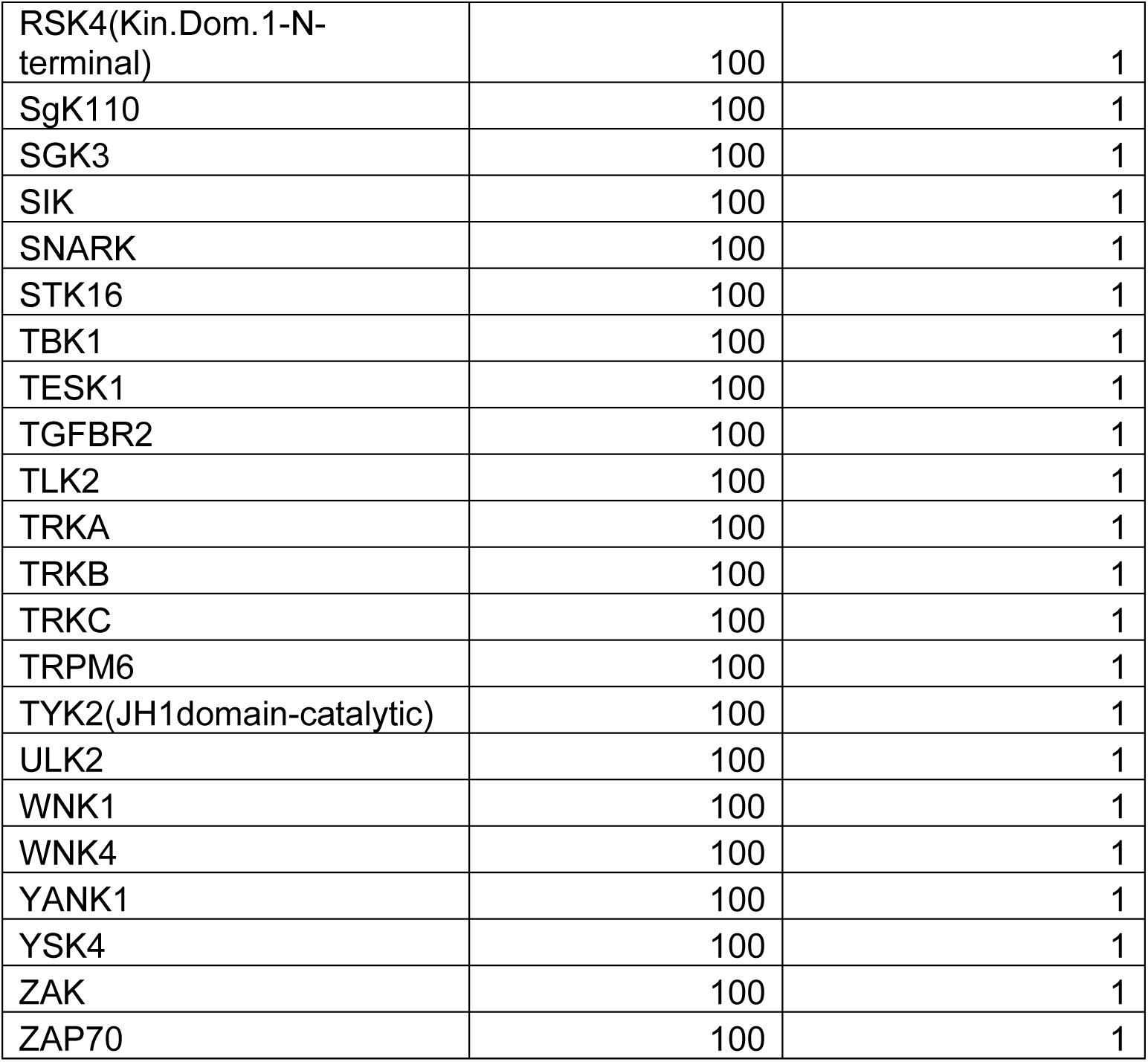
YNW-1. DiscoverX KINOME*scan* at 1 µM

**Table S6.** Biochemical activities against a panel of kinase targets

| YNW-1 |  |
| --- | --- |
| ERBB2 (HER2) | 126 |
| ERBB4 (HER4) | 73.6 |
| FLT4 (VEGFR3) | 108 |
| MAPK14 (p38 alpha) Direct | 732 |
| TEK (Tie2) | 250 |
| EPHA6 | >10000 |

<sup>a</sup>IC<sub>50</sub> (nM) against EGFR L858R, BTK, JAK3, ITK and BLK were obtained with Z'-LYTE
biochemical assay. <sup>b</sup> K<sub>d</sub> (nM) of EPHA6 was obtained from KdELECT assays (DiscoverX).

## Chemistry

Unless otherwise stated, all reagents and solvents were purchased from commercial suppliers and used without further purification. ¹H NMR spectra were recorded on a Bruker A400 or A500 spectrometer operating at 400 or 500 MHz. Chemical shifts (δ) are reported in parts per million (ppm, d) relative to tetramethyl silane (TMS) as an internal standard. Coupling constants (J) are given in hertz (Hz). Signal multiplicities are designated as follows: s (singlet), br (broad singlet), d (doublet), t (triplet), q (quartet), and m (multiplet). All compounds used in biological assays were confirmed to be >95% purities by analytical reverse-phase HPLC.

## Synthesis of ZNL-1

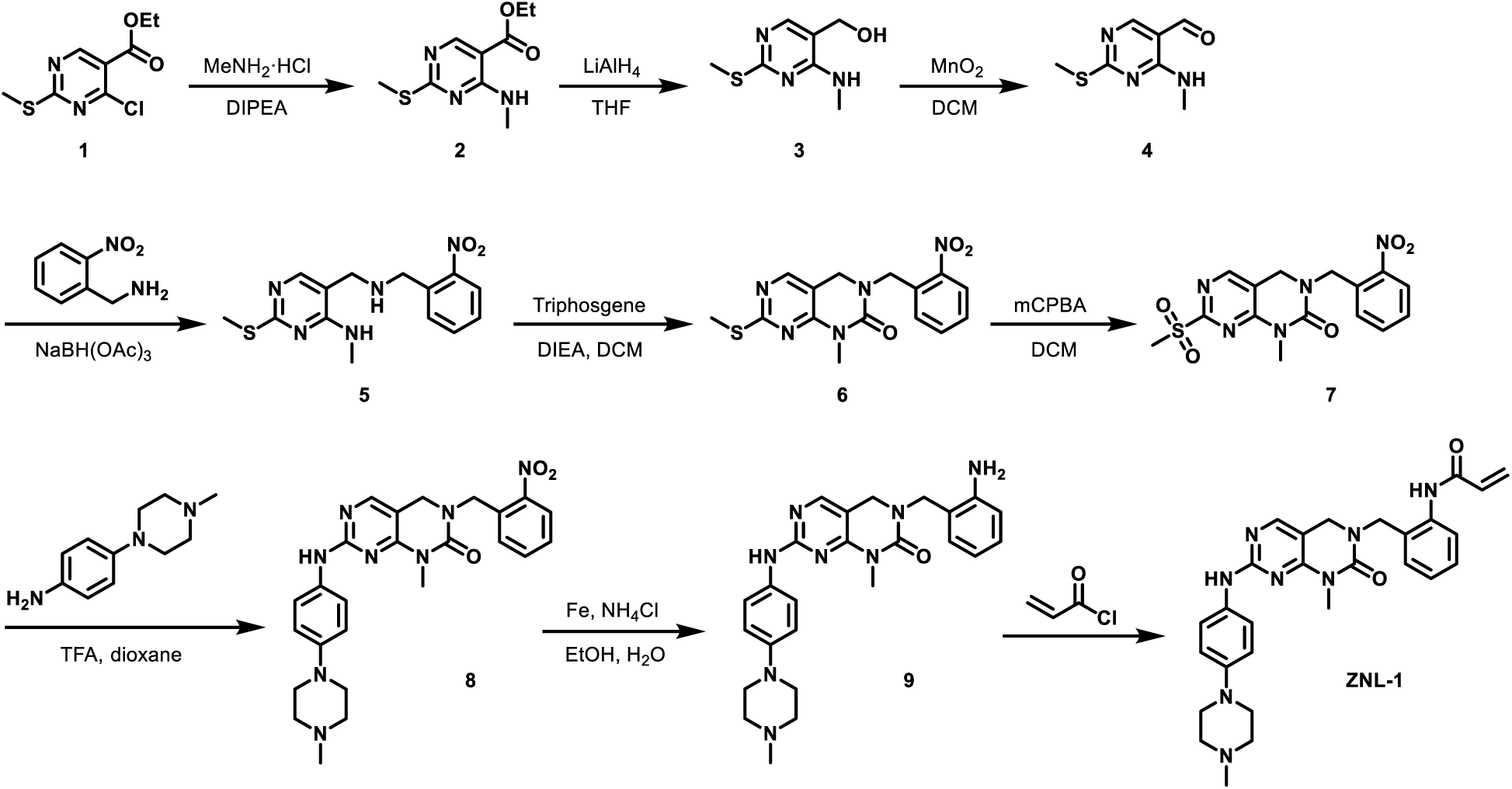

### Compound 2

To a solution of ethyl 4-chloro-2-methylsulfanyl-pyrimidine-5-carboxylate **1** (10 g, 42.98 mmol, 1 eq) in THF (200 mL) was added TEA (10.00 g, 98.85 mmol, 13.76 mL, 2.3 eq) and methenamine hydrochloride (6.67 g, 98.85 mmol, 2.3 eq) at 0°C.The mixture was stirred at 0°C for 1 hr and at 25°C for 12 hr. LCMS showed complete consumption of the reactant and formation of the desired product mass. The reaction mixture was concentrated under reduced pressure. The residue was diluted with H_2_O 100 mL and extracted with EtOAc (100 mL × 3). The combined organic layers were washed with brine (100 mL × 3), dried over Na_2_SO_4_, filtered and concentrated under reduced pressure to give **2** (9.4 g) as a white solid.

### Compound 3

The suspension of LAH (2.35 g, 62.04 mmol, 1.5 eq) in THF (40 mL) was cooled at 0°C and to this the solution of **2** (9.4 g, 41.36 mmol, 1 eq) in THF (20 mL) was added dropwise under argon and allowed to stir at 0 °C for 30 min. The reaction was quenched by addition of saturated MgSO_4_ solution (20 mL). The mixture and filtered and the filtrate concentrated under reduced pressure to give **3** (5.5 g, crude) as a yellow solid.

### Compound 4

To a solution of **3** (3.5 g, 18.89 mmol, 1 eq) in DCM (50 mL) was added MnO_2_ (9.86 g, 113.36 mmol, 6 eq). The mixture was stirred at 40°C for 12 hr. The reaction mixture was filtered and the filtrate was concentrated under reduced pressure to give a residue, the residue was purified by flash silica gel chromatography (ISCO®; 20 g SepaFlash® Silica Flash Column, Eluent of 5∼50% Ethyl acetate/Petroleum ether gradient @ 80mL/min) to give **4** (3 g, 16.37 mmol, 87% yield) as a yellow solid.

### Compound 5

To a mixture of (2-nitrophenyl)methanamine hydrochloride (2 g, 10.60 mmol, 1 eq) in DCE (30 mL) was added TEA to adjust pH∼=8, **4** (2.91 g, 15.91 mmol, 1.5 eq) and AcOH was added to adjust pH∼=5 and the mixture was stirred at 25°C for 2 hr. NaBH(OAc)_3_ (6.74 g, 31.81 mmol, 3 eq) was added to the reaction. The mixture was stirred at 25°C for 12 hr. The reaction mixture was partitioned between H_2_O 30 mL and DCM 30 mL. The water phase was separated, extracted with DCM (30 mL × 3), dried over Na_2_SO_4_, filtered and concentrated under reduced pressure to give a residue. The residue was purified by flash silica gel chromatography (ISCO®; 20 g SepaFlash® Silica Flash Column, Eluent of 0∼65% Ethyl acetate/Petroleum ether gradient @ 80mL/min) to give **5** (2.65 g, 8.30 mmol, 78.25% yield) as a yellow oil.

### Compound 6

To a solution of **5** (2.65 g, 8.30 mmol, 1 eq) in DCM (30 mL) was added DIEA (3.22 g, 24.89 mmol, 4.34 mL, 3 eq) and bis(trichloromethyl) carbonate (984.88 mg, 3.32 mmol, 0.4 eq) in DCM (10 mL) at 0°C. The mixture was stirred at 25°C for 2 hr. LCMS showed complete consumption of the reactant and formation of the desired product mass. The reaction mixture was partitioned between NaHCO_3_ 50 mL and DCM 50 mL. The water phase was separated, extracted with DCM (50 mL × 3), dried over Na_2_SO_4_, filtered and concentrated under reduced pressure to give a residue. The residue was purified by flash silica gel chromatography (ISCO®; 20 g SepaFlash® Silica Flash Column, Eluent of 0∼100% Ethyl acetate/Petroleum ether gradient @80 mL/min) to give **6** (700 mg, 2.03 mmol, 24.43% yield) as a yellow solid.

### Compound 7

A solution of m-CPBA (1.53 g, 7.53 mmol, 85% purity, 4 eq) in DCM (10 mL) was added to **6** (650 mg, 1.88 mmol, 1 eq) in DCM (10 mL) at 0°C. The mixture was stirred at 25°C for 12 hr. LCMS showed complete consumption of the reactant and formation of the desired product mass. The residue was diluted with Na_2_SO_3_ (10%) 14 mL and extracted with DCM (10 mL × 3). The combined organic layers were washed with NaHCO_3_ (10 mL × 3), dried over Na_2_SO_4_, filtered and concentrated under reduced pressure to give a residue. The crude material was suspended in ethanol and stirred at room temperature for 30 min. The solid was filtered, washed with ethanol and dried. The residue was purified by flash silica gel chromatography (ISCO®; 12 g SepaFlash® Silica Flash Column, Eluent of 0∼100% Ethyl acetate/Petroleum ether gradient @ 80 mL/min) to give **7** (450 mg, 1.19 mmol, 63.36% yield) as yellow gum.

### Compound 8

To a solution of **7** (160 mg, 423.98 umol, 1 eq) in dioxane (5 mL) was added TFA (72.52 mg, 635.97 umol, 47.09 uL, 1.5 eq) and 4-(4-methylpiperazin-1-yl) aniline (121.64 mg, 635.97 umol, 1.5 eq). The mixture was stirred at 120°C for 12 hr. The reaction mixture was concentrated under reduced pressure. The residue was diluted with NaHCO_3_ 10 mL and extracted with EtOAc (10 mL × 3). The combined organic layers were washed with brine (10 mL × 3), dried over Na_2_SO_4_, filtered and concentrated under reduced pressure to give a residue. The residue was purified by prep-TLC (SiO2, DCM: MeOH = 10:1) to give **8** (180 mg, 305.81 umol, 72.13% yield, 83% purity) as a yellow solid.

### Compound 9

To a solution of **8** (180 mg, 368.44 umol, 1 eq) in EtOH (6 mL) and H_2_O (2 mL) was added NH_4_Cl (39.42 mg, 736.89 umol, 2 eq) and stirred at 25°C for 5 min. Then the mixture was heated to 90°C. Fe (82.30 mg, 1.47 mmol, 4 eq) was added to the mixture. The mixture was stirred at 90°C for 2 hr. The reaction mixture was filtered and the filtrate dried in vacuum to give a residue. The residue was purified by prep-TLC (SiO_2_, 10% methol in DCM) to give **9** (110 mg, 239.9 umol, 65% yield) as a yellow solid.

### ZNL-1

To a solution of **9** (50 mg, 109.04 umol, 1 eq) and DIPEA (42.3 mg, 327.11 umol, 54.1 uL, 3 eq) in DMF (1 mL) was added and prop-2-enoyl chloride (9.87 mg, 109.04 umol, 8.9 uL, 1 eq) at 0°C. The mixture was stirred at 0°C for 90 min. The reaction mixture was concentrated under reduced pressure to give a residue. The residue was purified by prep-HPLC (FA condition, Phenomenex Luna C18 75*30mm*3um; [water (FA)-ACN]; B%:1%-30%, 8min) to give **ZNL-1** (11 mg, 18.54 umol, 17.01% yield, 94.166% purity, FA) as a white solid. MS (ESI): m/z = 513.2 [M+H] ^+^

^1^HNMR (400 MHz, DMSO-*d_6_*) *δ* ppm 9.93 (s, 1 H), 9.24 (s, 1 H), 8.18 (s, 1 H), 8.01 (s, 1 H), 7.77 (d, J=8.0 Hz, 1 H), 7.56 (d, J=8.0 Hz, 2 H), 7.35-7.22 (m, 2 H), 7.21-7.17 (m, 1 H), 6.88 (d, J=8.0 Hz, 2 H), 6.53-6.46 (m, 1 H), 6.30-6.25 (m, 1 H), 5.82-5.79 (m, 1 H), 4.61 (s, 2 H), 4.26 (s, 2 H), 3.31 (s, 3 H), 3.07-3.04 (m, 4 H), 2.51-2.48 (m, 4 H), 2.25 (s, 3 H).

## Synthesis of ZNL-2 (*S*)/(*R*)

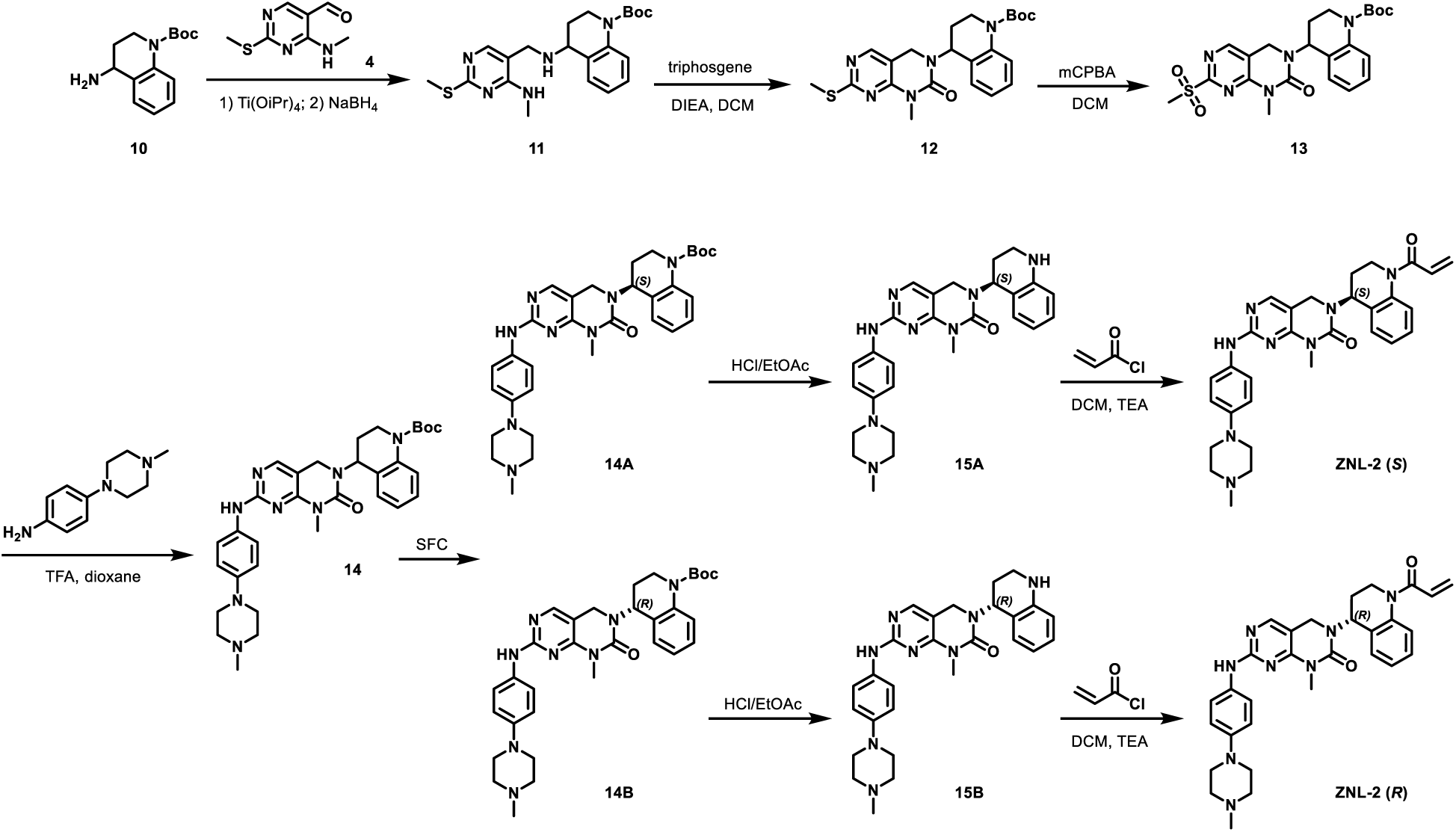

### Compound **11**

To a solution of **4** (3 g, 16.37 mmol, 1 eq) in MeOH (100 mL) was added **10** (4.07 g, 16.37 mmol, 1 eq) and Ti(O*^i^*Pr)_4_ (13.96 g, 49.12 mmol, 14.50 mL, 3 eq). The mixture was stirred at 30°C for 12 hr. Then NaBH_4_ (3.72 g, 98.24 mmol, 6 eq) was added to the mixture at 0°C. The mixture was stirred at 25°C for 3 hr. The reaction mixture was concentrated under reduced pressure to give a residue. The reaction mixture was quenched by addition NH_4_Cl (100 mL) at 0°C. The reaction mixture was extracted with acetate ethyl (100 mL × 3). The combined organic layers were washed with brine (30 mL × 2), dried over Na_2_SO_4_, filtered and concentrated under reduced pressure to give a residue. The residue was purified by flash silica gel chromatography (ISCO®; 40 g SepaFlash® Silica Flash Column, Eluent of 0∼50% Ethyl acetate/Petroleum ether gradient @ 120 mL/min) to **11** (6 g) as a light-yellow oil.

### Compound **12**

To a solution **11** (6 g, 14.44 mmol, 1 eq) and DIEA (9.33 g, 72.19 mmol, 12.57 mL, 5 eq) in DCM (100 mL) was added triphosgene (4.28 g, 14.44 mmol, 1 eq) at 0°C. The mixture was stirred at 25°C for 12 hr. The reaction mixture was partitioned between NaHCO_3_ (100 mL) and dDCM (100 mL × 2). The organic phase was separated, washed with brine (100 mL × 1), dried over Na_2_SO_4_, filtered and concentrated under reduced pressure to give a residue. The residue was purified by flash silica gel chromatography (ISCO®; 40 g SepaFlash® Silica Flash Column, Eluent of 0∼50% Ethyl acetate/Petroleum ether gradient @ 120 mL/min) to **12** (5.8 g, 11.71 mmol, 81.13% yield, 89.181% purity) as a light-yellow oil.

### Compound **13**

To a solution of **12** (3 g, 6.79 mmol, 1 eq) in DCM (50 mL) was added m-CPBA (2.21 g, 10.87 mmol, 85% purity, 1.6 eq) at 0°C. The mixture was stirred at 25°C for 12 hr. The residue was diluted with Na_2_SO_3_ (50 mL) and extracted with dichloromethane (50 mL × 3). The combined organic layers were washed with NaHCO_3_ (50 mL × 3), dried over Na_2_SO_4_, filtered and concentrated under reduced pressure to **13** (3.2 g, crude) as a white solid.

### Compound 14, 14A, 14B

To a solution of **13** (2 g, 4.22 mmol, 1 eq) in dioxane (20 mL) was added TFA (722.34 mg, 6.34 mmol, 469.05 uL, 1.5 eq) and 4-(4-methylpiperazin-1-yl) aniline (969.40 mg, 5.07 mmol, 1.2 eq). The mixture was stirred at 120°C for 12 hr. The residue was purified by prep-HPLC (FA condition, column: Phenomenex luna C18 (250×70mm, 15 um); mobile phase: [water (FA)-ACN]; B%: 30%-60%, 20min) to give product **14**. Then **14** was separated by SFC column: DAICEL CHIRALPAK AD (250mm×50mm,10um); mobile phase: A: CO_2_ B: EtOH (0.1%IPAm, v/v); B%: 60%-60%, 7min to give **14A** (Rt = 1.709 min, 500 mg, 855.12 umol, 20.25% yield) and **14B** (Rt =1.993 min, 530 mg, 906.43 umol, 21.46% yield) as a brown solid.

### Compound **15A, 15B**

To a solution of **14A** or **14B** (350 mg, 598.6 umol, 1 eq) in HCl/EtOAc (4 M, 3.5 mL, 23.3 eq) was stirred at 25°C for 1 hr. The reaction mixture was concentrated under reduced pressure to give product **15A** or **15B** as yellow solid used into the next step without further purification.

### ZNL-2 (*S*)

To a solution of **15A** (125 mg, 239.9 umol, 1 eq, HCl) and DIEA (93.0 mg, 719.7 umol, 125.3 uL, 3 eq) in DMF (2 mL) was added a DCM solution of prop-2-enoyl chloride (26.1 mg, 287.9 umol, 23.5 uL, 1.2 eq) at 0°C. The reaction was stirred at 0°C for 1 hr. The reaction mixture was filtered. The residue was purified by prep-HPLC (FA condition; column:Phenomenex luna C18 100×40mm×3 um; mobile phase: [water(FA)-ACN]; B%: 1%-40%,8min), and then the residue was purified by prep-HPLC (neutral condition; column: Waters Xbridge Prep OBD C18 150×40mm×10um; mobile phase: [water (NH_4_HCO_3_)-ACN]; B%: 20%-50%, 8min) to yield **ZNL-2 (*S*)** (100% purity, 69 mg) as a white solid. MS (ESI): m/z = 539.2 [M+H] ^+^

VT (Variable Temperature) -^1^H NMR (400 MHz, DMSO-*d_6_*) *δ* = 8.96 - 8.80 (m, 1H), 7.99 - 7.84 (m, 1H), 7.59 - 7.43 (m, 2H), 7.30 - 7.15 (m, 3H), 6.86 (d, J = 9.0 Hz, 2H), 6.72 - 6.51 (m, 1H), 6.29 - 6.12 (m, 1H), 5.78 - 5.68 (m, 1H), 5.68 - 5.50 (m, 1H), 4.25 - 4.05 (m, 2H), 3.96 - 3.82 (m, 1H), 3.80 - 3.69 (m, 1H), 3.62 - 3.49 (m, 1H), 3.34 (s, 3H), 3.10 - 3.04 (m, 4H), 2.46 (br d, J = 4.9 Hz, 4H), 2.28 - 2.17 (m, 4H), 2.15 - 1.96 (m, 1H) **ZNL-2 (*S*)** was also made starting with **(*S*)-10**, and the absolutely configuration was confirmed by SFC comparing.

### ZNL-2 (*R*)

To a solution of **15B** (300 mg, 575.75 umol, 1 eq, HCl) and TEA (174.78 mg, 1.73 mmol, 240.41 uL, 3 eq) in DCM (5 mL) was added a DCM solution of prop-2-enoyl chloride (52.11 mg, 575.75 umol, 46.95 uL, 1 eq). The mixture was stirred at 0°C for 1 hr. The reaction mixture was concentrated under reduced pressure to give a residue. The residue was purified by prep-HPLC (FA condition, column: Phenomenex Luna C18 75×30mm×3um ;mobile phase: [water(FA)-ACN]; B%:1%-35%,8min) and (NH4HCO3 condition, column: Phenomenex C18 75×30mm×3um;mobile phase: [water (NH4HCO3)-ACN]; B%: 20%-55%,8min) to give **ZNL-2 (*R*)** (76.10 mg, 139.89 umol, 24.30% yield, 99.016% purity) as a yellow solid. MS (ESI): m/z = 539.2 [M+H] ^+^

**ZNL-2 (*R*)** was also made starting with **(*R*)-10**, and the absolutely configuration was confirmed by SFC comparing.

## Synthesis of ZNL-3

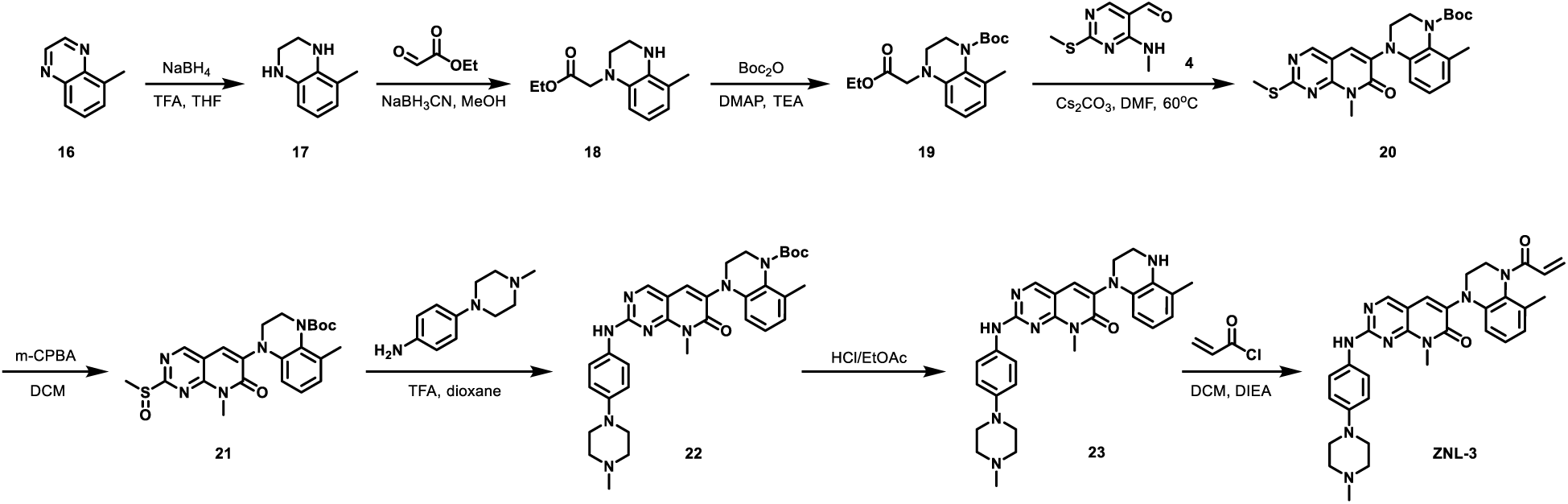

### Compound 17

To a solution of 5-methylquinoxaline **16** (5.0 g, 34.7 mmol, 1 *eq*) in THF (50 mL) was added NaBH_4_ (4.6 g, 121.4 mmol, 3.5 *eq*). The mixture was stirred at 0°C for 0.5 hr. TFA (7.1 g, 62.4 mmol, 4.6 mL, 1.8 *eq*) was added to the mixture at 0°C. The mixture was stirred at 0°C for 0.5 hr. Then the mixture was stirred at 20°C for 12 hr. The reaction mixture was added water (100 mL) at 0°C. The mixture was extracted with ethyl acetate (20 mL × 3). The combined organic layers were washed with brine (15 mL × 2), dried over Na_2_SO_4_, filtered and concentrated under reduced pressure to give a residue. The residue was purified by column chromatography (SiO_2_, Petroleum ether/Ethyl acetate=5/1 to 3/1) to give 5-methyl-1,2,3,4-tetrahydroquinoxaline **17** (4.7 g, 31.7 mmol, 91.4% yield) as red oil.

### Compound 18

To a solution of **17** (4.3 g, 29.0 mmol, 1 *eq*) and ethyl 2-oxoacetate (6.5 g, 31.9 mmol, 50% purity, 1.1 *eq*) in MeOH (100 mL) was adjust pH to 4-5 by AcOH, the mixture was stirred at 25°C for 3 hr, then NaBH_3_CN (5.5 g, 87.0 mmol, 3 *eq*) was added, the mixture was stirred at 25°C for 12 hr. The mixture was concentrated to get a residue, the residue was quenched by Sat. NaHCO_3_ (20 mL) and then extracted by ethyl acetate (30 mL × 3), the combined organic layers was dried by Na_2_SO_4_, filtered and the filtrate was concentrated to get a residue. The residue was purified by column chromatography (SiO_2_, Petroleum ether /Ethyl acetate=10/1 to 3/1) to give ethyl 2-(5-methyl-3,4-dihydro-2H-quinoxalin-1-yl) acetate **18** (3 g, 12.8 mmol, 44.1% yield) as brown oil.

### Compound 19

To a solution of **18** (2.5 g, 10.7 mmol, 1 *eq*) in Boc_2_O (15 mL) was added DMAP (130.4 mg, 1.1 mmol, 0.1 *eq*) and TEA (5.4 g, 53.4 mmol, 7.4 mL, 5 *eq*). The mixture was stirred at 80°C for 48 hr. The mixture was concentrated to get a residue. The residue was purified by column chromatography (SiO_2_, Petroleum ether / Ethyl acetate=10/1 to 5/1) to give **19** (2.45 g, 7.3 mmol, 68.7% yield) as red oil.

### Compound 20

To a solution of **19** (2.4 g, 7.3 mmol, 1 *eq*) in DMF (20 mL) was added Cs_2_CO_3_ (7.2 g, 22.0 mmol, 3 *eq*) and 4-(methylamino)-2-methylsulfanyl-pyrimidine-5-carbaldehyde **4** (1.5 g, 8.1 mmol, 1.1 *eq*). The mixture was stirred at 80°C for 12 hr. The mixture was poured into water (100 mL) and then extracted by ethyl acetate (30 mL × 3), the organic layers was washed by brine (20 mL × 2), dried by Na_2_SO_4_, then filtered and the filtrate was concentrated to get a residue. The residue was purified by flash silica gel chromatography (25 g Silica Flash Column, Eluent of 0∼15% Ethyl acetate/Petroleum ether gradient @ 120 mL /min) to give **20** (2.1 g, 4.6 mmol, 63.2% yield) as a yellow solid.

### Compound 21

To a solution of **20** (2.1 g, 4.63 mmol, 1 *eq*) in DCM (50 mL) was added m-CPBA (1.7 g, 8.3 mmol, 85% purity, 1.8 eq). The mixture was stirred at 25°C for 12 hr. The reaction mixture was partitioned between Na_2_SO_3_ (30 mL) and dichloromethane (30 mL × 3). The organic phase was separated, washed with NaHCO_3_ (20 mL × 3), dried over Na_2_SO_4_, filtered and concentrated under reduced pressure to give a residue. The residue was purified by column chromatography (SiO_2_, Petroleum ether/Ethyl acetate=10/1 to 1/1) to give compound **21** (1 g, 2.1 mmol, 46.0% yield) as a red solid.

### Compound 22

To a solution of **21** (400 mg, 857.8 umol, 1 *eq*) in dioxane (4 mL) was added TFA (116.6 mg, 1022.2 umol, 75.9 uL, 1.2 *eq*) and 4-(4-methylpiperazin-1-yl) aniline (163.7 mg, 851.9 umol, 1 *eq*). The mixture was stirred at 80°C for 12 hr. The mixture was concentrated to get a residue. The residue was purified by prep-HPLC (FA condition; column: Phenomenex luna C18 100 × 40mm × 3 um; mobile phase: [water (TFA)-ACN]; B%: 25%-70%,8min) to give **22** (300 mg, 502.8 umol, 59.0% yield) as a red solid.

### Compound 23

To a solution of **22** (300 mg, 502.8 umol, 1 *eq*) in EtOAc (5 mL) was added HCl/EtOAc (10 mL, 4M). The mixture was stirred at 25°C for 1 hr. The mixture was concentrated to give **23** (250 mg, crude) as a yellow solid.

### ZNL-3

To a solution of **23** (100 mg, 187.6 umol, 1 eq, HCl) in DCM (1 mL) was added DIEA (97.0 mg, 750.4 umol, 130.7 uL, 4 eq) and prop-2-enoyl chloride (17.0 mg, 187.6 umol, 15.3 uL, 1 eq). The mixture was stirred at 0°C for 1 hr. The mixture was concentrated to get a residue. The residue was purified by prep-HPLC (FA condition; column: Phenomenex Luna C18 200 × 40mm × 10um; mobile phase: [water (FA)-ACN]; B%:5%-40%, 8min) to give compound **ZNL-3** (28.1 mg, 51.0 umol, 27.2% yield, 100% purity) as a white solid. MS (ESI): m/z = 551.3 [M+H] ^+^

^1^H NMR (400 MHz, DMSO-*d_6_*) *δ* = 9.89 (br d, *J* = 1.0 Hz, 1H), 8.71 (s, 1H), 7.90 (s, 1H), 7.66 (br d, *J* = 8.1 Hz, 2H), 6.99 - 6.86 (m, 3H), 6.64 (d, *J* = 8.0 Hz, 1H), 6.35 (d, *J* = 8.0 Hz, 1H), 6.31 - 6.23 (m, 2H), 5.73 (m, 1H), 4.88 (m, 1H), 3.70 (m, 1H), 3.63 - 3.55 (m, 3H), 3.41 (m, 1H), 3.13 - 3.06 (m, 4H), 3.01 (m, 1H), 2.48 - 2.43 (m, 4H), 2.22 (s, 3H), 2.15 - 2.01 (m, 3H)

## Synthesis of ZNL-3r

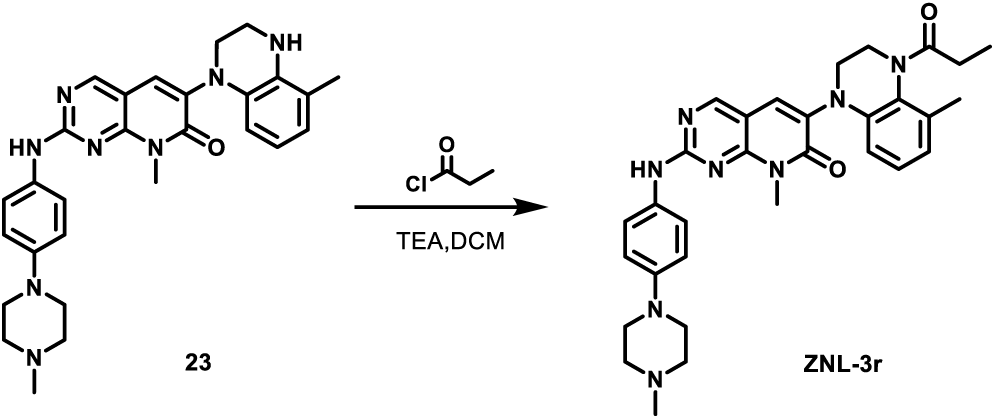

To a solution of **23** (60 mg, 112.6 umol, 1 *eq*, HCl) in DCM (3 mL) was added TEA (34.2 mg, 337.7 umol, 47.0 uL, 3 *eq*) and propanoyl chloride (20.8 mg, 225.1 umol, 20.8 uL, 2 *eq*) at 0°C. The mixture was stirred at 25°C for 12 hr. The reaction mixture was concentrated under reduced pressure to remove **solvent**. The residue was purified by prep-HPLC (**FA condition** column: Phenomenex C 18 75 × 30 mm × 3um; mobile phase: [water (FA)-ACN]; B%: 10%-45%, 8 min) to compound **ZNL-3r** (27.8 mg, 50.1 umol, 44.5% yield, 99.8% purity) as a yellow solid. MS (ESI): m/z = 553.5 [M+H] ^+^

^1^H NMR (400 MHz, DMSO-*d_6_*) *δ* = 9.66 (s, 1H), 8.70 (s, 1H), 7.82 (s, 1H), 7.75 - 7.66 (m, 2H), 7.09 - 6.97 (m, 2H), 6.92 - 6.81 (m, 1H), 6.70 - 6.58 (m, 1H), 6.33 (br d, *J* = 8.0 Hz, 1H), 5.06 - 4.45 (m, 1H), 3.62 (m, 5H), 3.40 - 3.25 (m, 5H), 2.88 (s, 3H), 2.50 - 2.45 (m, 4H), 2.45 - 1.89 (m, 5H), 1.04 (br s, 3H)

## Synthesis of ZNL-4

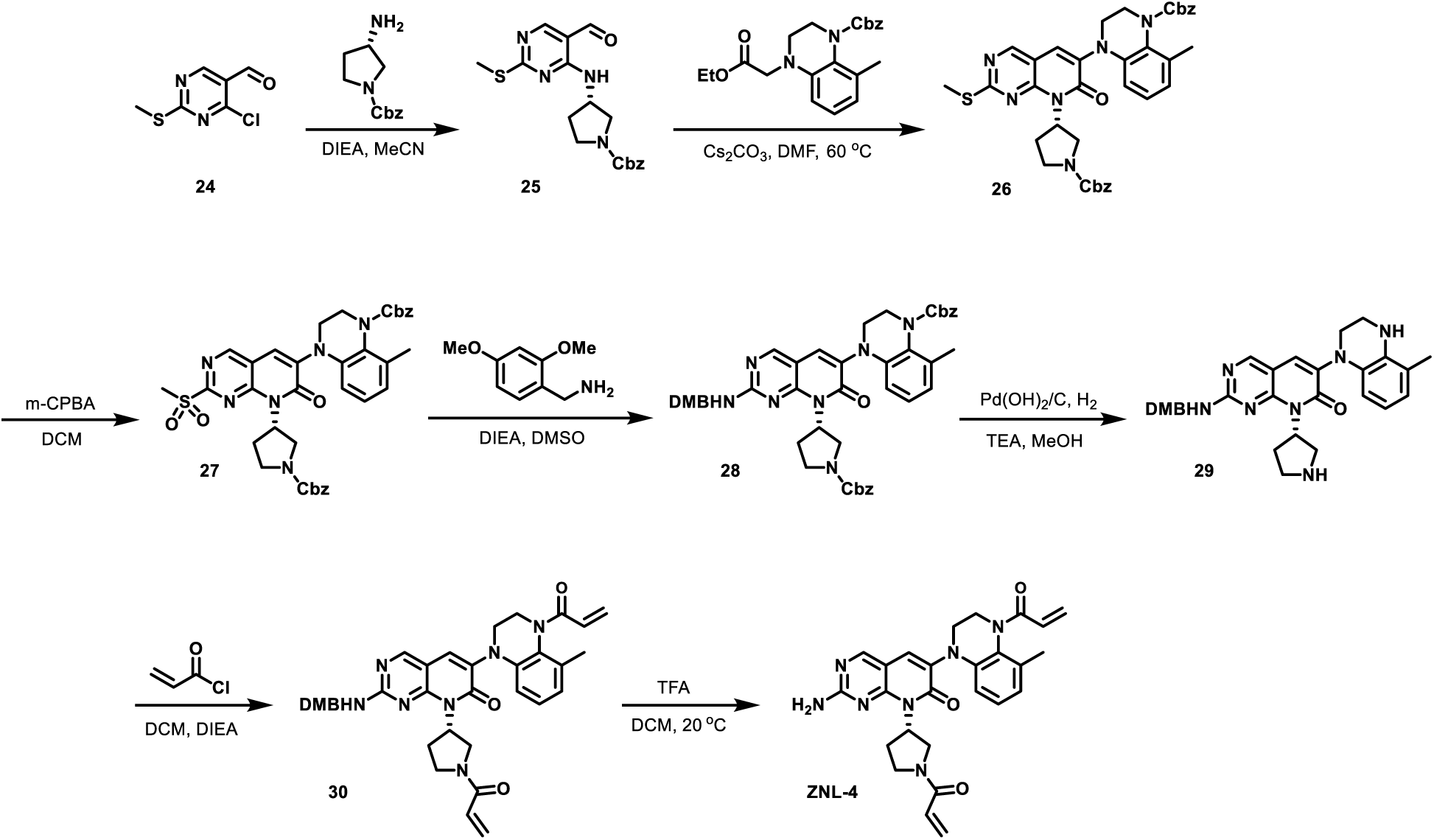

### Compound 25

To a solution of 4-chloro-2-methylsulfanyl-pyrimidine-5-carbaldehyde **24** (1.56 g, 8.27 mmol, 1.1 *eq*) in ACN (100 mL) was added DIEA (2.92 g, 22.55 mmol, 3.93 mL, 3 *eq*) and benzyl (3*S*)-3-aminopyrrolidine-1-carboxylate (1.66 g, 7.52 mmol, 1 *eq*). The mixture was stirred at 20°C for 1 hr. The reaction mixture was diluted with H_2_O 100 ml and then extracted with DCM (50 mL * 4). The combined organic layers were dried over Na_2_SO_4_, filtered and concentrated under reduced pressure to give a residue. The residue was purified by flash silica gel chromatography (ISCO®; 20 g SepaFlash® Silica Flash Column, Eluent of 0 ∼ 20% Ethyl acetate/ Petroleum ether gradient @ 80 mL/ min). Compound **25** (2 g, 5.26 mmol, 70.00% yield, 98.00% purity) was obtained as a yellow solid.

### Compound 26

To a solution of **25** (1 g, 2.68 mmol, 1 *eq*) in DMF (20 mL) was added Cs_2_CO_3_ (2.62 g, 8.05 mmol, 3 *eq*) and benzyl 4-(2-ethoxy-2-oxo-ethyl)-8-methyl-2,3-dihydroquinoxaline-1-carboxylate (989.22 mg, 2.68 mmol, 1 *eq*). The mixture was stirred at 60°C for 2 hr. LCMS showed benzyl (S)-3-((5-formyl-2-(methylthio) pyrimidin-4-yl) amino) pyrrolidine-1-carboxylate (1 g, 2.68 mmol, 1 *eq*) was consumed completely and 14% of desired compound was detected. The reaction mixture was diluted with H_2_O 100 ml and then extracted with DCM (50 mL * 4). The combined organic layers were dried over Na_2_SO_4_, filtered and concentrated under reduced pressure to give a residue. The residue was purified by flash silica gel chromatography (ISCO®; 40g SepaFlash® Silica Flash Column, Eluent of 0 ∼ 30% Ethyl acetate/ Petroleum ether gradient @ 80 mL/ min). Compound **26** (1.3 g, 1.61 mmol, 60.09% yield, 84% purity) was obtained as a yellow solid.

### Compound 27

To a solution of **26** (1.2 g, 1.77 mmol, 1 *eq*) in DCM (3 mL) was added *m*-CPBA (539.97 mg, 2.66 mmol, 85% purity, 1.5 *eq*). The mixture was stirred at 20°C for 12 hr. The reaction mixture was diluted with Na_2_SO_3_ saturated 100 ml and then extracted with DCM (50 mL *4). The combined organic layers were dried over Na_2_SO_4_, filtered and concentrated under reduced pressure to give a residue. Compound **27** (1.3 g, crude) was obtained as a yellow oil.

### Compound 28

To a solution of **27** (650 mg, 917.07 μmol, 1 *eq*) in Tol. (0.5 mL) was added DIEA (474.10 mg, 3.67 mmol, 638.95 μL, 4 *eq*) and (2,4-dimethoxyphenyl) methanamine (168.67 mg, 1.01 mmol, 151.55 μL, 1.1 *eq*). The mixture was stirred at 20°C for 1 hr. The reaction mixture was diluted with H_2_O 100ml and then extracted with DCM (50 mL * 4). The combined organic layers were dried over Na_2_SO_4_, filtered and concentrated under reduced pressure to give a residue. The residue was purified by flash silica gel chromatography (ISCO®; 20 g SepaFlash® Silica Flash Column, Eluent of 0 ∼ 20% Petroleum ether: Ethyl acetate @ 80 mL/ min). Compound **28** (550 mg, 674.47 μmol, 73.55% yield, 97.6% purity) was obtained as a yellow solid.

### Compound 29

To a solution of **28** (470 mg, 590.54 μmol, 1 *eq*) in MeOH (25 mL) and THF (2 mL) was added Pd(OH)_2_ (0.400 g, 590.54 μmol, 10% purity, 1.00 *eq*) and TEA (59.76 mg, 590.54 μmol, 82.20 μL, 1 *eq*). The mixture was stirred at 20 °C for 1 hr under H_2_. The mixture was filtered through a Celite pad, and the filtrate was concentrated to give a residue. Compound **29** (330 mg, crude) was obtained as a yellow solid.

### Compound 30

To a solution of **29** (150 mg, 284.30 μmol, 1 *eq*) in DCM (5 mL) was added TEA (86.30 mg, 852.89 μmol, 118.71 μL, 3 *eq*) and prop-2-enoyl chloride (46.32 mg, 511.74 μmol, 41.58 μL, 1.8 *eq*) at 0°C. The mixture was stirred at 20°C for 1 hr. The reaction mixture was concentrated under reduced pressure to give a residue. Compound **30** (150 mg, crude) was obtained as a yellow solid.

### ZNL-4

To a solution of **30** (150 mg, 235.96 μmol, 1 *eq*) in DCM (5 mL) was added TFA (807.11 mg, 7.08 mmol, 525.81 μL, 30 *eq*). The mixture was stirred at 12°C for 12 hr. The mixture was filtered through a Celite pad, and the filtrate was concentrated to give a residue. The residue was purified by prep-HPLC (column: Phenomenex Luna C18 75 * 30 mm * 3 um; mobile phase: [H_2_O (0.1% TFA)-ACN]; gradient: 15% - 40% B over 8.0 min). **ZNL-4** (62.00 mg, 98.68 μmol, 41.82% yield, 95.43% purity, TFA) was obtained as a yellow solid. MS (ESI): *m*/*z* = 486.4 [M+H] ^+^

^1^H NMR (400 MHz, DMSO-*d_6_*) *δ* = 8.59 (s, 1H), 7.85 (br s, 1H), 7.28 (br s, 2H), 7.05 - 6.76 (m, 1H), 6.70 - 6.46 (m, 2H), 6.34 (br d, *J* = 8.0 Hz, 1H), 6.28 (br s, 1H), 6.26 (br d, *J* = 2.5 Hz, 1H), 6.17 - 6.10 (m, 1H), 5.88 - 5.55 (m, 2H), 4.86 (br dd, *J* = 3.8, 12.3 Hz, 1H), 4.13 - 3.91 (m, 1H), 3.89 - 3.59 (m, 4H), 3.49 (br s, 1H), 3.05 - 2.90 (m, 1H), 2.80 - 2.62 (m, 1H), 2.27 - 1.90 (m, 5H)

## Synthesis of ZNL-5

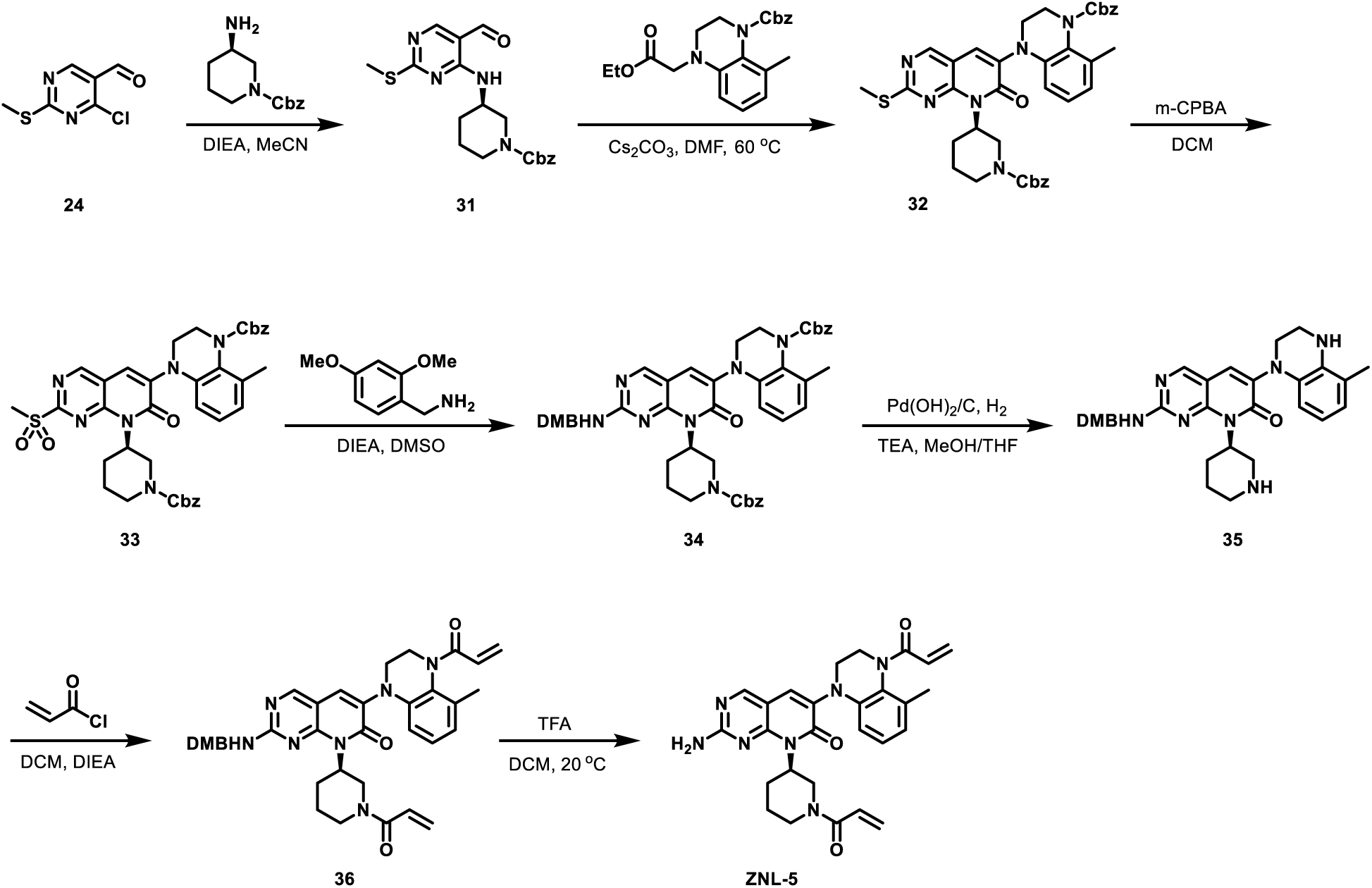

### Compound 31

To a solution of 4-chloro-2-methylsulfanyl-pyrimidine-5-carbaldehyde **24** (5.40 g, 28.64 mmol, 1 *eq*) in MeCN (75 mL) was added DIEA (11.10 g, 85.91 mmol, 14.96 mL, 3 *eq*) and benzyl (3*R*)-3-aminopiperidine-1-carboxylate (6.37 g, 27.20 mmol, 0.95 *eq*) at 20°C. The mixture was stirred at 20°C for 4 hr. The residue was purified by column chromatography (SiO_2_, Petroleum ether/ Ethyl acetate = 1/0 to 1/2, TLC (Petroleum ether: Ethyl acetate = 1/2, Rf = 0.55). Compound **31** (5 g, 12.94 mmol, 45.18% yield) was obtained as a yellow oil.

### Compound 32

To a solution of **31** (1 g, 2.59 mmol, 1 *eq*) in DMF (25 mL) was added Cs_2_CO_3_ (2.53 g, 7.76 mmol, 3 *eq*) and benzyl 4-(2-ethoxy-2-oxo-ethyl)-8-methyl-2,3-dihydroquinoxaline-1-carboxylate (953.32 mg, 2.59 mmol, 1 *eq*). The mixture was stirred at 60°C for 2 hr. The reaction mixture was filtered and the filtrate concentrated under reduced pressure to give a residue. The residue was purified by column chromatography (SiO_2_, Petroleum ether/ Ethyl acetate = 1/0 to 1/1, TLC: Petroleum ether/Ethyl acetate = 1/1, Rf = 0.42). Compound **32** (1.1 g, 1.59 mmol, 61.54% yield) was obtained as a yellow solid.

### Compound 33

To a solution of **32** (1.1 g, 1.59 mmol, 1 *eq*) in DCM (15 mL) was added *m*-CPBA (858.70 mg, 3.98 mmol, 80% purity, 2.5 *eq*) at 0°C. The mixture was stirred at 25°C for 2 hr. The reaction mixture was quenched by addition *sat.aq.* Na_2_SO_3_ 10 mL at 0°C and then extracted with DCM 20 mL (10 mL * 2). The combined organic layers were washed with brine 20 mL (5 mL * 4), dried over Na_2_SO_4_, filtered and concentrated under reduced pressure to give a yellow solid. Compound **33** (1.0 g, 1.38 mmol, 86.88% yield) was obtained as a yellow solid.

### Compound 34

To a solution of **33** (500 mg, 691.75 μmol, 1 *eq*) in DMSO (5 mL) was added DIEA (268.20 mg, 2.08 mmol, 361.46 μL, 3 *eq*) and (2,4-dimethoxyphenyl)methaneamine (231.33 mg, 1.38 mmol, 207.84 μL, 2 *eq*). The reaction mixture was concentrated under reduced pressure to give a residue. The residue was purified by column chromatography (SiO_2_, Petroleum ether/ Ethyl acetate = 1/0 to 1/1). Compound **34** (350 mg, 411.41 μmol, 59.47% yield, 95.2% purity) was obtained as a yellow solid.

### Compound 35

To a solution of **34** (350 mg, 432.15 μmol, 1 *eq*) in MeOH (3 mL) and THF (0.5 mL) was added TEA (43.73 mg, 432.15 μmol, 60.15 μL, 1 *eq*) and Pd(OH)_2_/C (100 mg, 432.15 μmol, 1.00 *eq*) under N_2_ atmosphere. The suspension was degassed and purged with H_2_ for 3 times. The mixture was stirred under H2 (15 Psi) at 25°C for 2 hr. The reaction mixture was filtered, and the filtrate was concentrated under reduced pressure to give a yellow solid. Compound **35** (180 mg, 332.32 μmol, 76.90% yield) was obtained as a yellow solid.

### Compound 36

To a solution **36** (130.00 mg, 240.01 μmol, 1 *eq*) in DCM (2 mL) was added TEA (72.86 mg, 720.03 μmol, 100.22 μL, 3 *eq*) and prop-2-enoyl chloride (43.45 mg, 480.02 μmol, 39.00 μL, 2 *eq*) at 0°C. The mixture was stirred at 25 °C for 2 hr. Compound **36** (250 mg, crude) was obtained as a yellow solid.

### ZNL-5

To a solution of **36** (250 mg, 404.70 μmol, 1 *eq*) in DCM (4 mL) was added TFA (1.23 g, 10.77 mmol, 0.8 mL, 26.61 *eq*). The mixture was stirred at 25 °C for 2 hr. The mixture was concentrated by vacuum under reduced pressure. The residue was purified by prep-HPLC (column: Phenomenex Luna C18 100 * 30 mm*5 um; mobile phase: [H_2_O (0.2% FA) - ACN]; gradient: 1%-55% B over 8.0 min). Compound **ZNL-5** (13.46 mg, 24.63 μmol, 40.85% yield, 99.85% purity) was obtained as a yellow solid. MS (ESI): *m*/*z* = 500.3 [M+H] ^+^

^1^H NMR (400 MHz, DMSO-*d_6_*) *δ* = 8.56 (s, 1H), 7.81 (s, 1H), 7.29 (br s, 2H), 7.02 - 6.71 (m, 2H), 6.66 - 6.49 (m, 1H), 6.40 - 6.18 (m, 3H), 6.11 (br d, J=16.63 Hz, 1H), 5.86 - 5.58 (m, 2H), 4.86 (m, 1H), 4.55 - 4.32 (m, 1H), 4.15 - 3.98 (m, 1H), 3.65 (br d, *J* = 6.25 Hz, 1H), 3.49 - 3.40 (m, 2H), 3.31 - 3.21 (m, 2H), 3.04 - 2.92 (m, 1H), 2.17 - 1.96 (m, 3H), 1.90 - 1.66 (m, 2H), 1.55 - 1.19 (m, 2H)

## Synthesis of ZNL-6

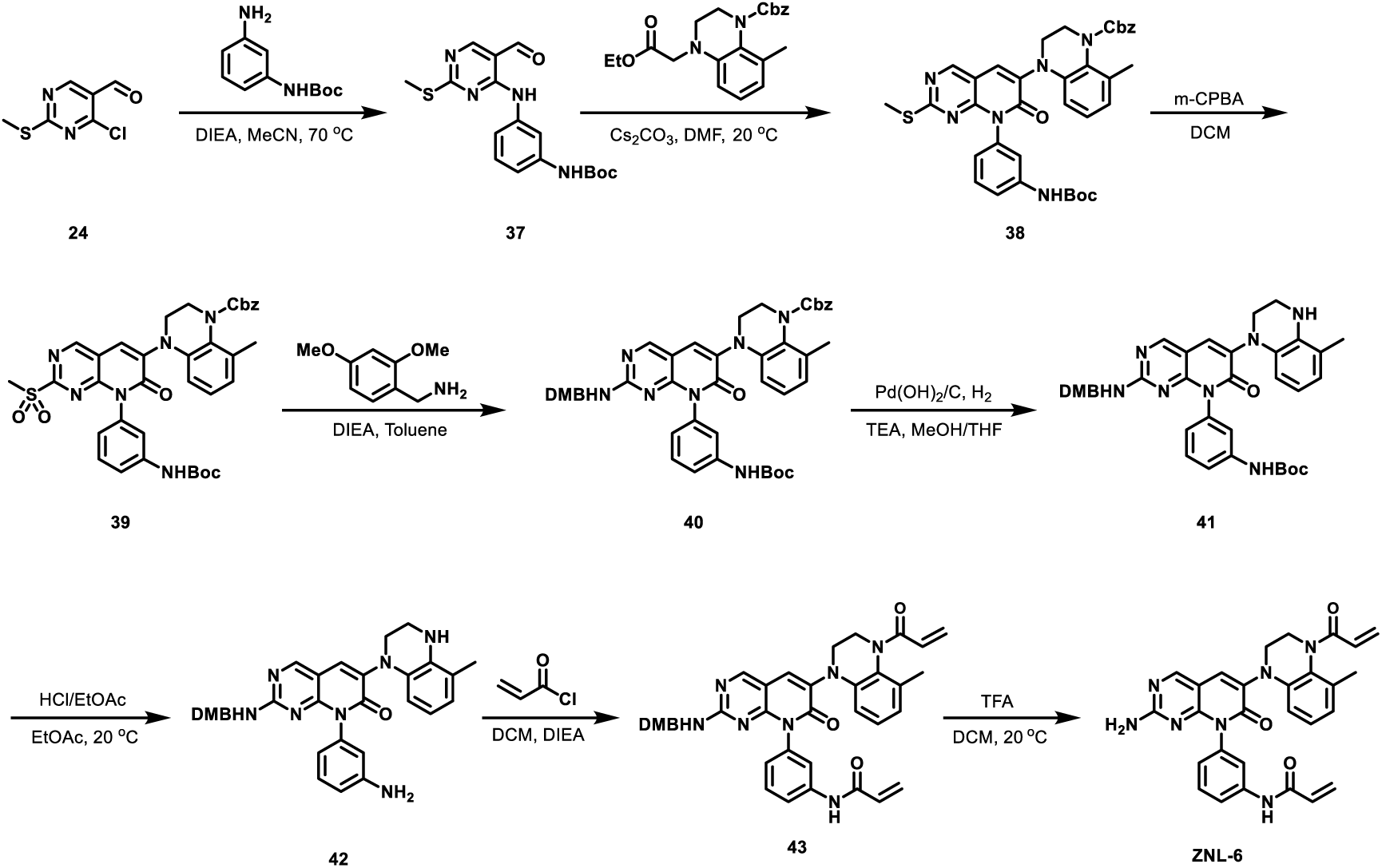

### Compound 37

To a solution of 4-chloro-2-(methylthio) pyrimidine-5-carbaldehyde **24** (3 g, 15.90 mmol, 1 *eq*) and tert-butyl N-(3-aminophenyl) carbamate (3.31 g, 15.90 mmol, 1 *eq*) in MeCN (30 mL) was added DIEA (4.11 g, 31.81 mmol, 5.54 mL, 2 *eq*). The mixture was stirred at 70°C for 12 hr. The reaction mixture was concentrated under reduced pressure to remove solvent. The residue was diluted with H_2_O 30 mL and extracted with EtOAc (30 mL * 3). The combined organic layers were dried over Na_2_SO_4_, filtered and concentrated under reduced pressure to give a residue. The residue was purified by flash silica gel chromatography (ISCO®; 12 g SepaFlash® Silica Flash Column, Eluent of 0 ∼ 30% Ethyl acetate/Petroleum ethergradient @ 80 mL/min). Compound **37** (4.3 g, 11.93 mmol, 75.01% yield) was obtained as a yellow oil.

### Compound 38

To a solution of 37 (500 mg, 1.39 mmol, 1 eq) in DMF (20 mL) was added Cs_2_CO_3_ (1.36 g, 4.16 mmol, 3 eq) and benzyl 4-(2-ethoxy-2-oxoethyl)-8-methyl-3,4-dihydroquinoxaline-1(2H)-carboxylate (511.09 mg, 1.39 mmol, 1 eq). The mixture was stirred at 20 °C for 2 hr. The reaction mixture was diluted with H_2_O 100 ml and then extracted with DCM (50 mL * 4). The combined organic layers were dried over Na_2_SO_4_, filtered and concentrated under reduced pressure to give a residue. The residue was purified by flash silica gel chromatography (ISCO®; 40g SepaFlash® Silica Flash Column, Eluent of 0∼20% Ethyl acetate/Petroleum ethergradient @ 80 mL/min). Compound **38** (370 mg, 500.92 μmol, 36.11% yield, 90% purity) was obtained as a white solid.

### Compound 39

To a solution of **38** (370 mg, 556.58 μmol, 1 *eq*) in DCM (10 mL) was added *m*-CPBA (169.50 mg, 834.87 μmol, 85% purity, 1.5 *eq*). The mixture was stirred at 20 °C for 12 hr. The reaction mixture was diluted with Na_2_SO_3_ saturated 20 ml and then extracted with DCM (10 mL *4). The combined organic layers were dried over Na_2_SO_4_, filtered and concentrated under reduced pressure to give a residue. Compound 39 (380 mg, crude) was obtained as a yellow solid.

### Compound 40

To a solution of **39** (360 mg, 516.67 μmol, 1 eq) in Tol. (3 mL) was added DIEA (267.10 mg, 2.07 mmol, 359.97 μL, 4.0 *eq*), (2,4-dimethoxyphenyl) methanamine (95.03 mg, 568.34 μmol, 85.38 μL, 1.1 *eq*). The mixture was stirred at 20°C for 1 hr. The reaction mixture was diluted with H_2_O 100 ml and then extracted with DCM (50 mL * 4). The combined organic layers were dried over Na_2_SO_4_, filtered and concentrated under reduced pressure to give a residue. The residue was purified by flash silica gel chromatography (ISCO®; 20 g SepaFlash® Silica Flash Column, Eluent of 0∼20% Petroleum ether:Ethyl acetate @ 80 mL/min). Compound **40** (400 mg, 474.57 μmol, 91.85% yield, 93% purity) was obtained as a white solid.

### Compound 41

To a solution of **40** (400 mg, 510.29 μmol, 1 *eq*) in MeOH (2.5 mL) and THF (0.4 mL) was added Pd(OH)_2_/C (0.400 g, 510.29 μmol, 10% purity, 1.00 eq) and TEA (51.64 mg, 510.29 μmol, 71.03 μL, 1 eq). The mixture was stirred at 20°C for 1 hr under H_2_ (15 Psi). The mixture was filtered through a Celite pad, and the filtrate was concentrated to give a residue. Compound **41** (330 mg, crude) was obtained as a yellow solid.

### Compound 42

To a solution of **41** (150 mg, 230.86 μmol, 1 eq) in EtOAc (2 mL) was added HCl/EtOAc (4 M, 2 mL, 34.65 eq). The mixture was stirred at 20°C for 2 hr. The reaction mixture was concentrated under reduced pressure to give a residue. Compound **41** (130 mg, crude, HCl) was obtained as a yellow solid.

### Compound 43

To a solution of **42** (130 mg, 236.53 μmol, 1 eq) in DCM (5 mL) was added TEA (119.67 mg, 1.18 mmol, 164.61 μL, 5 eq) and prop-2-enoyl chloride (38.53 mg, 425.75 μmol, 34.59 μL, 1.8 eq) at 0°C. The mixture was stirred at 20°C for 1 hr. The mixture was concentrated by vacuum under reduced pressure. Compound **43** (155.57 mg, crude) was obtained as a yellow solid.

### ZNL-6

To a solution of **43** (130 mg, 197.65 μmol, 1 eq) in DCM (2.5 mL) was added TFA (901.46 mg, 7.91 mmol, 587.27 μL, 40 *eq*). The mixture was stirred at 20 °C for 12 hr. The mixture was concentrated by vacuum under reduced pressure. The mixture was concentrated by vacuum under reduced pressure. The residue was purified by prep-HPLC (column: Phenomenex Luna C18 100*30 mm*5 um;mobile phase: [H_2_O (0.2% FA) - ACN]; gradient:1%-55% B over 8.0 min). Compound **ZNL-6** (8.60 mg, 15.47 μmol, 22.89% yield, 99.55% purity) was obtained as a yellow solid. MS (ESI): *m*/*z* = 508.3 [M+H] ^+^

^1^H NMR (400 MHz, DMSO-*d*_6_) *δ* = 10.30 (br s, 1 H), 8.62 (br s, 1H), 7.95 (br s, 1H), 7.76 - 7.62 (m, 1H), 7.54 (br s, 1H), 7.44 (br d, *J* = 6.8 Hz, 1H), 7.21 - 6.83 (m, 4H), 6.63 - 6.35 (m, 3H), 6.33 - 6.12 (m, 3H), 5.85 - 5.60 (m, 2H), 5.00 - 4.72 (m, 1H), 3.77 - 3.62 (m, 1H), 3.19 - 3.16 (m, 1H), 2.98-2.95 (m, 1H), 2.14 - 1.98 (m, 3H).

## Synthesis of ZNL-7

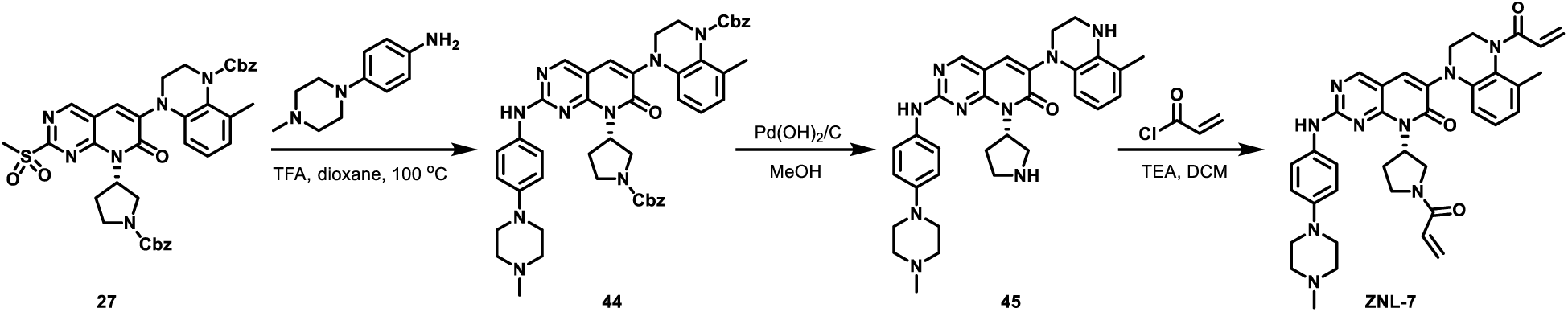

### Compound 44

To a solution of 27 (650 mg, 917.07 μmol, 1 eq) in dioxane (0.5 mL) was added TFA (156.85 mg, 1.38 mmol, 102.18 μL, 1.5 eq) 4-(4-methylpiperazin-1-yl) aniline (192.95 mg, 1.01 mmol, 1.1 eq). The mixture was stirred at 120°C for 1 hr. The reaction mixture was concentrated under reduced pressure to give a residue. The residue was purified by prep-HPLC (column: Phenomenex luna C18 100 * 40 mm *5 um; mobile phase: [H_2_O (0.1% TFA)-ACN]; gradient: 40% - 70% B over 8.0 min). Compound 44 (400 mg, 427.04 μmol, 46.57% yield, 99.71% purity, TFA) was obtained as a yellow solid.

### Compound 45

To a solution of **44** (200 mg, 243.92 μmol, 1 *eq*) in MeOH (25 mL) and THF (2 mL) was added Pd(OH)_2_ (0.200 g, 10% purity) and TEA (24.68 mg, 243.92 μmol, 33.95 μL, 1 *eq*). The mixture was stirred at 20°C for 1 hr under H_2_. The mixture was filtered through a Celite pad, and the filtrate was concentrated to give a residue. Compound **45** (130 mg, crude) was obtained as a yellow solid.

### ZNL-7

To a solution of **45** (120 mg, 217.52 μmol, 1 *eq*) in DCM (5 mL) was added TEA (66.03 mg, 652.55 μmol, 90.83 μL, 3 *eq*) and prop-2-enoyl chloride (35.44 mg, 391.53 μmol, 31.81 μL, 1.8 eq) at 0°C. The mixture was stirred at 20°C for 1 hr. The mixture was filtered through a Celite pad, and the filtrate was concentrated to give a residue. The residue was purified by prep-HPLC (column: Phenomenex Luna C18 75 * 30 mm* 3 um; mobile phase: [H_2_O (0.1% TFA) - ACN]; gradient:15% - 40% B over 8.0 min). Compound **ZNL-7** (88.11 mg, 113.10 μmol, 52.00% yield, 99.33% purity, TFA) was obtained as a white solid. MS (ESI): *m*/*z* = 660.5 [M+H] ^+^

^1^H NMR (400 MHz, DMSO-D*_6_*) *δ* = 10.00 - 9.77 (m, 1H), 8.72 (d, J = 2.1 Hz, 1H), 7.90 (br d, J = 3.5 Hz, 1H), 7.50 (br t, J = 8.5 Hz, 2H), 6.96 - 6.89 (m, 1H), 6.88 - 6.81 (m, 2H), 6.72 - 6.62 (m, 1H), 6.59 - 6.44 (m, 1H), 6.42 - 6.31 (m, 1H), 6.29 - 6.26 (m, 1H), 6.21 - 6.07 (m, 2H), 5.79 - 5.57 (m, 2H), 4.98 - 4.78 (m, 1H), 4.32 - 3.93 (m, 1H), 3.90 - 3.79 (m, 1H), 3.76 - 3.64 (m, 2H), 3.13 - 2.75 (m, 8H), 2.45 (br s, 4H), 2.23 (s, 3H), 2.18 - 1.98 (m, 5H)

## Synthesis of ZNL-8

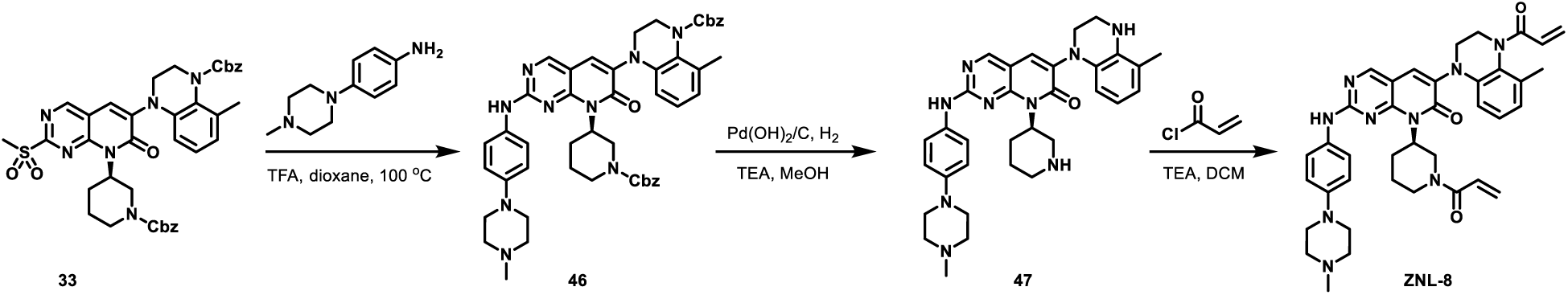

### Compound 46

To a solution of **33** (500 mg, 691.75 μmol, 1 *eq*) and 4-(4-methylpiperazin-1-yl) aniline (132.31 mg, 691.75 μmol, 1 *eq*) in dioxane (5 mL) was added TFA (230.25 mg, 2.02 mmol, 0.15 mL, 2.92 *eq*). The mixture was stirred at 120°C for 12 hr. The mixture was concentrated by vacuum under reduced pressure. The residue was purified by prep-HPLC (column: Phenomenex Luna C18 100 * 30 mm * 5 um; mobile phase: [H_2_O (0.2% FA) - ACN]; gradient: 40% - 70% B over 8.0 min). Compound **46** (400 mg, 479.63 μmol, 69.34% yield) was obtained as a yellow solid.

### Compound 47

A mixture of Pd(OH)_2_ (200 mg, 1.42 mmol, 3.96 *eq*) in MeOH (1.5 mL) and THF (1.5 mL) was added to **46** (300 mg, 359.72 μmol, 1 *eq*) and TEA (218.10 mg, 2.16 mmol, 300.00 μL, 5.99 *eq*) was degassed and purged with H_2_ for 3 times, then the mixture was stirred at 25°C for 2 hr under H_2_ (15 Psi) atmosphere. The reaction mixture was filtered, and the filtrate was concentrated to give a residue. Compound **47** (150 mg, crude) was obtained as a yellow oil.

### ZNL-8

To a solution of **47** (150 mg, 265.15 μmol, 1 *eq*) in DCM (2 mL) was added TEA (80.49 mg, 795.46 μmol, 110.72 μL, 3 *eq*) and prop-2-enoyl chloride (48.00 mg, 530.31 μmol, 43.09 μL, 2 *eq*) at 0°C. The mixture was stirred at 25°C for 2 hr. The mixture was concentrated by vacuum under reduced pressure. The residue was purified by prep-HPLC (column: Phenomenex Luna C18 100 * 30 mm* 5um; mobile phase: [H_2_O (0.2% FA)-ACN]; gradient: 1% - 55% B over 8.0 min). Compound **ZNL-**(39.57 mg, 54.52 μmol, 20.56% yield, 99.17% purity) was obtained as a yellow solid. MS (ESI): *m*/*z* = 674.5 [M+H] ^+^

^1^H NMR (400 MHz, DMSO-*d_6_) δ* = 10.01 - 9.49 (m, 1H), 8.71 (s, 1H), 7.88 (s, 1H), 7.63 - 7.22 (m, 2H), 7.03 - 6.72 (m, 4H), 6.65 (br d, *J* = 7.1 Hz, 1H), 6.41 - 6.03 (m, 4H), 5.84 - 5.71 (m, 1H), 5.64 - 5.31 (m, 1H), 5.00 - 4.74 (m, 1H), 4.67 - 4.34 (m, 1H), 4.23 - 4.03 (m, 1H), 3.88 - 3.62 (m, 2H), 3.17 - 2.97 (m, 6H), 2.79 - 2.53 (m, 2H), 2.46 (br d, *J* = 3.4 Hz, 4H), 2.23 (s, 3H), 2.17 - 2.02 (m, 3H), 1.93 - 1.62 (m, 2H), 1.49 - 1.10 (m, 2H)

## Synthesis of YNW-1

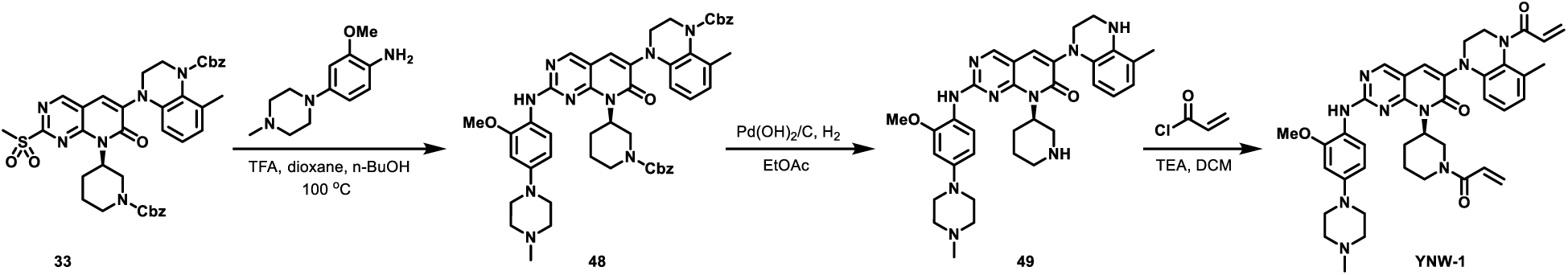

### Compound 48

To a solution of benzyl 33 (2 g, 2.8 mmol, 1 eq) in dioxane (16 mL) and n-BuOH (32 mL) was added 2-methoxy-4-(4-methylpiperazin-1-yl) aniline (1.2 g, 5.5 mmol, 2 eq) and TFA (1.9 g, 16.6 mmol, 1.2 mL, 6 eq). The mixture was stirred at 100°C for 12 hr. The reaction mixture was concentrated under reduced pressure to give a residue. The residue was purified by flash silica gel chromatography (120 g Silica Flash Column, Eluent of 0∼7% Methanol/Dichloromethane gradient @ 120 mL/min) to give 48 (5.5 g, 6.4 mmol, ee%=100%) as a yellow solid.

### Compound 49

To a solution of Pd(OH)_2_/C (5 g, 20% purity) in EtOAc (50 mL) was added **48** (5 g, 5.8 mmol, 1 eq) under N_2_ atmosphere. The suspension was degassed and purged with H_2_ for 3 times. The mixture was stirred under H_2_ (15 Psi) at 40°C for 12 hr. Then the reaction mixture was filtered and concentrated under reduced pressure to give a residue to give **49** (4 g, crude) as a yellow solid.

### YNW-1

To a solution of **49** (4 g, 6.7 mmol, 1 eq) in DCM (150 mL) was added TEA (2.0 g, 20.1 mmol, 2.8 mL, 3 eq). Then prop-2-enoyl chloride (1.2 g, 13.4 mmol, 1.1 mL, 2 eq) was added to the mixture at 0°C. The mixture was stirred at 0°C for 1 hr. The reaction mixture was concentrated under reduced pressure to remove solvent. The residue was purified by prep-HPLC (FA condition column: Phenomenex luna C18 250mm × 100mm × 15um; mobile phase: [H_2_O (0.02%FA)-ACN]; gradient: 10%-40% B over 28.0 min) to give **YNW-1** (852 mg, 98.63% purity) as a yellow solid. MS (ESI): m/z = 704.4 [M+H] ^+^

VT ^1^HNMR (400 MHz, DMSO-*d*_6_) *δ* = 8.66 - 8.60 (m, 1H), 8.59 - 8.50 (m, 1H), 7.81 - 7.78 (m, 1H), 7.45 - 7.28 (m, 1H), 6.94 - 6.82 (m, 1H), 6.72 - 6.60 (m, 3H), 6.49 - 6.43 (m, 1H), 6.34 (br d, *J* = Hz, 1H), 6.30 - 6.20 (m, 2H), 6.11 - 6.01 (m, 1H), 5.75 - 5.65 (m, 1H), 5.61 (br d, *J* = 9.4 Hz, 1H), 5.16 - 5.03 (m, 1H), 4.96 - 4.74 (m, 1H), 4.23 - 3.92 (m, 2H), 3.78 (s, 3H), 3.64 - 3.39 (m, 2H), 3.22 - 3.10 (m, 5H), 3.08 - 2.89 (m, 1H), 2.70 - 2.53 (m, 2H), 2.49 - 2.47 (m, 4H), 2.28 - 2.24 (m, 3H), 2.15 - 2.06 (m, 3H), 1.77 - 1.57 (m, 2H), 1.47 - 1.31 (m, 1H)

## Synthesis of YNW-2

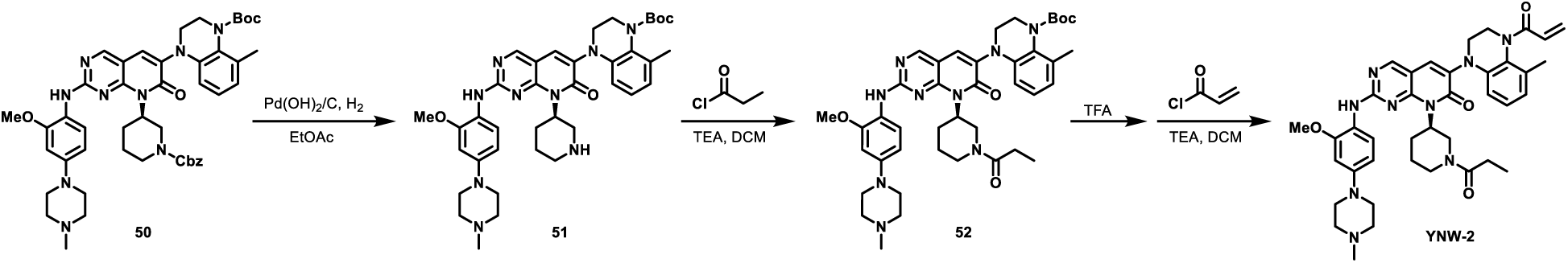

### Compound 51

To a solution of Pd(OH)_2_/C in EtOAc was added **50** (1 eq) under N_2_ atmosphere. The suspension was degassed and purged with H_2_ for 3 times. The mixture was stirred under H_2_ (15 Psi) at 40 °C for 12 hr. The reaction mixture was filtered and concentrated under reduced pressure to give a residue to give **51**.

### Compound 52

To a solution of **51** in DCM was added TEA (3 eq). Then propionyl chloride (2 eq) was added to the mixture at 0 °C. The mixture was stirred at 0 °C for 1 hr. The reaction mixture was concentrated under reduced pressure to remove solvent. Then purified by ISCO (methanol in DCM 0-30%) to give **52**.

### YNW-2

To a solution of **52** in DCM was added TEA (3 eq). Then prop-2-enoyl chloride (2 eq) was added to the mixture at 0 °C. The mixture was stirred at 0 °C for 1 hr. The reaction mixture was concentrated under reduced pressure to remove solvent. Then purified by HPLC to give **YNW-2** as a yellow solid. MS (ESI): m/z = 706.1 [M+H] ^+^

^1^H NMR (500 MHz, DMSO-*d_6_*) *δ* (ppm) = 9.83 - 9.72 (m, 1 H), 8.65 (s, 1 H), 7.85 (s, 1 H), 6.89 (m, 1H), 6.72 (m, 1H), 6.59 (m, 3H), 6.38 - 6.18 (m, 3H), 5.72 (m, 1H), 4.91 - 4.84 (m, 1H), 3.91 - 3.63 (m, 6H), 3.56 (d, *J* = 10.0 Hz, 2H), 3.41 (m, 1H), 3.18 (m, 2H), 2.92 (m, 5H), 2.11-2.02 (s, 6H), 1.56 - 1.28 (m, 8H), 0.93 (m, 4H).

## Synthesis of YNW-3

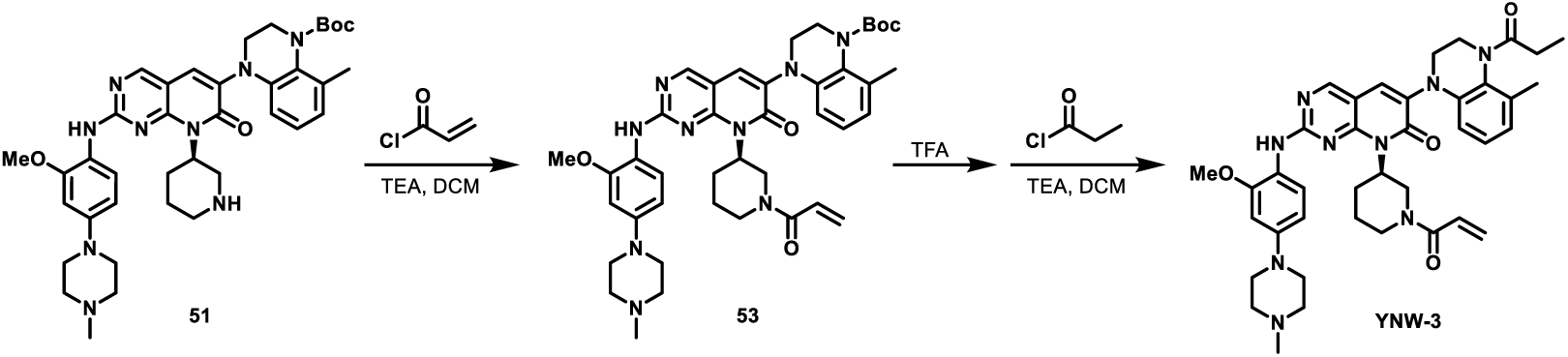

### Compound 53

To a solution of **51** in DCM was added TEA (3 eq). Then prop-2-enoyl chloride (2 eq) was added to the mixture at 0 °C. The mixture was stirred at 0°C for 1 hr. The reaction mixture was concentrated under reduced pressure to remove solvent. Then purified by ISCO (methanol in DCM 0-30%) to give **53**.

### YNW-3

To a solution of **53** in DCM was added TEA (3 eq). Then propionyl chloride (2 eq) was added to the mixture at 0 °C. The mixture was stirred at 0 °C for 1 hr. The reaction mixture was concentrated under reduced pressure to remove solvent. Then purified by HPLC to give **YNW-3** as a yellow solid. MS (ESI): m/z = 706.1 [M+H] ^+ 1^H NMR (500 MHz, DMSO-*d_6_*) *δ* (ppm) = 9.75 (m, 1H), 8.67 (s, 1H), 7.86-7.80 (m, 1H), 6.89 (m, 1H), 6.74-6.65 (m, 3H), 6.53 (m, 1H), 6.37-6.25 (m, 2H), 6.06 (m, 1H), 5.70-5.57 (m, 1 H), 4.85-4.77 (m, 1H), 3.82 (m, 6H), 3.56 (m, 2H), 3.41-3.32 (m, 1H), 3.19 (m, 2H), 2.97-2.83 (m, 5H), 2.21 (s, 3H), 2.16-1.96 (m, 3H), 1.94-1.20 (m, 8H), 1.03 (m, 4H).

